# Colorectal Cancer Stem Cell Subtypes Orchestrate Distinct Tumor Microenvironments

**DOI:** 10.1101/2024.04.25.591144

**Authors:** Linzi Hosohama, Delia F. Tifrea, Kevin Nee, Sung Yun Park, Jie Wu, Amber N. Habowski, Cassandra Van, Marcus M. Seldin, Robert A. Edwards, Marian L. Waterman

## Abstract

Several classification systems have been developed to define tumor subtypes in colorectal cancer (CRC). One system proposes that tumor heterogeneity derives in part from distinct cancer stem cell populations that co-exist as admixtures of varying proportions. However, the lack of single cell resolution has prohibited a definitive identification of these types of stem cells and therefore any understanding of how each influence tumor phenotypes. Here were report the isolation and characterization of two cancer stem cell subtypes from the SW480 CRC cell line. We find these cancer stem cells are oncogenic versions of the normal Crypt Base Columnar (CBC) and Regenerative Stem Cell (RSC) populations from intestinal crypts and that their gene signatures are consistent with the “Admixture” and other CRC classification systems. Using publicly available single cell RNA sequencing (scRNAseq) data from CRC patients, we determine that RSC and CBC cancer stem cells are commonly co-present in human CRC. To characterize influences on the tumor microenvironment, we develop subtype-specific xenograft models and we define their tumor microenvironments at high resolution via scRNAseq. RSCs create differentiated, inflammatory, slow growing tumors. CBCs create proliferative, undifferentiated, invasive tumors. With this enhanced resolution, we unify current CRC patient classification schema with TME phenotypes and organization.

- *Two cancer stem cell subtypes are isolated and determined to be commonly found as mixed populations in human CRC*
- *The heterogeneous SW480 cell line models and unifies two binary cancer stem cell classifications of colorectal cancer*
- *High resolution analysis of xenograft tumors reveals that CRC stem cells are dominant, context-independent organizers of their TME*
- *RSC/YAP/STAT3-positive CRC stem cells create differentiated tumors that attempt tissue generation via an inflammatory, angiogenic, oncofetal program*
- *CBC/MYC-positive CRC stem cells create tumors that are proliferative, undifferentiated, and invasive*

**Graphical Abstract:** 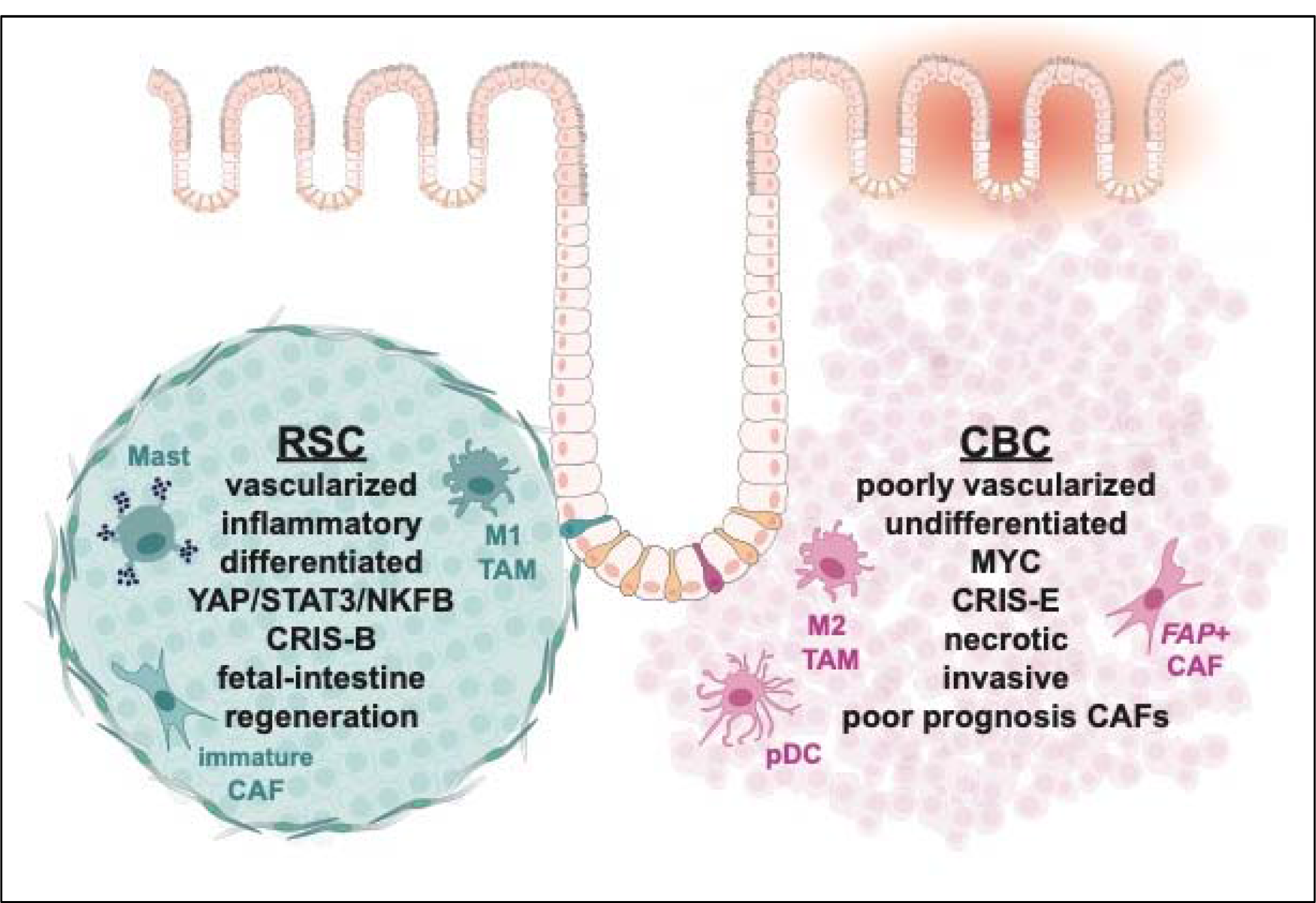

## Introduction

While initiation of cancer is thought to arise from a single transformed cell, progression to a tumor mass results in a population of cancer cells with a great deal of heterogeneity. Canonical thought ascribes genetic mutations to be the main drivers of this process, giving rise to the outgrowth of stable clonal subpopulations. However, the establishment of intra-tumor heterogeneity can also derive from transcriptionally distinct cellular states in isogenic cancer cells triggered by pressures in the tumor microenvironment^1^. To understand the complexity of heterogeneous tumors, several groups have used transcriptomic data from colorectal cancer (CRC) patients or PDX biopsies to develop classification structures and categorize this heterogeneity. For example, the CMS and CRIS systems each utilize transcriptomic signatures of signal transduction pathways including Wnt, inflammation, and others to define 4 and 5 CRC subtypes, respectively^2,3^. More recently, other groups have reduced the number of subtypes to “binary” categorizations of CRC based on either the Hippo Signal Transduction pathway (“YAP^ON^” versus “YAP^OFF^”)^4,5^, a revision-reduction of the CMS classification system to iCMS2 versus iCMS3^6^, or by association to two intestinal stem cell states, Regenerative Stem Cell (RSC) versus Crypt Based Columnar (CBC) cells^7–11^. How these different subtype signatures compare with each other, whether they are co-expressed in the same cells or in distinct populations, and how each subtype influences the tumor microenvironment (TME) is not yet known.

These CRC classification structures are based on bulk RNA sequencing of patient tumors or patient-derived organoids, even though there is wide acknowledgment of intra-tumor heterogeneity with respect to the signatures used to classify them^12^. Indeed, we and others have noted how intra-tumor heterogeneity is evident at the cell-to-cell level in CRC and even in tumor xenografts created from human colon cancer cell lines such as SW480^13–15^. To account for this, Vazquez et al. recently proposed that CRC tumors comprise a varying “admixture” of RSC-like and CBC-like cancer stem cells^7^. In their schema (CBC) cells are the stem cells that are LGR5+ and Wnt^HI^, and rapidly cycling to maintain crypt homeostasis (Figure 1A, e.g. *LGR5, SOX4, PROX1, FGFR4*)^11,16,17^. RSC refers to the less abundant population of quiescent stem cells that are activated upon injury to repair damaged intestinal crypts^18^. This stem cell population expresses gene programs of wounding and fetal-intestine (Figure 1A, e.g. *WNT5A, TACSTD2, CLU, GSN, IFITM1*)^8,9,19,20^. Whether the admixture classification model can account for heterogeneity at the individual cell level in CRC and whether it aligns with the other signaling-based classification schema is not known. Beyond the observation that CRC tumors are heterogeneous, there is much that is not understood about how different cancer stem cell subtypes influence the TME and overall phenotype. Tumor modeling of distinct CRC cancer subtype populations could greatly improve our understanding, but to achieve this, new methods that enable the capture, characterization, and culture of individual cancer cell populations are needed.

**Figure 1:**
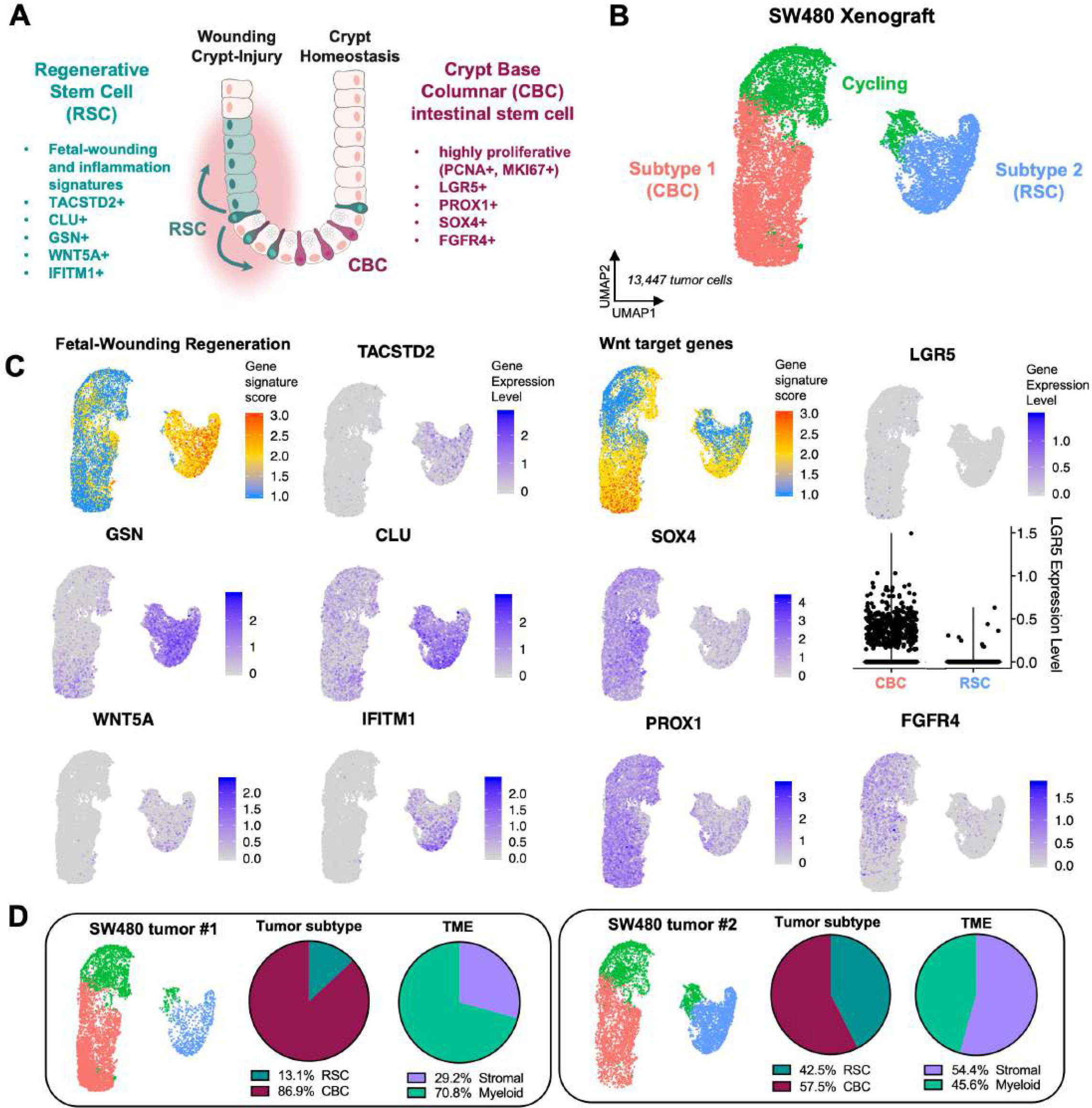
Two cancer stem cell subtypes comprise SW480 xenograft tumors. A: Schematic diagram showing location of different stem cell pools in the colon crypt during homeostasis and injury. B: UMAP clustering of human tumor cells from SW480 xenografts reveals two populations, Subtype 1 (CBC) and Subtype 2 (RSC). C: Gene expression plots for CBC biomarkers (Wnt target genes, *LGR5, SOX4, PROX1, FGFR4)* and RSC biomarkers (Fetal-Wounding Regeneration signature, *TACSTD2, GSN, CLU, WNT5A, IFITM1).* D: SW480 xenograft replicates contain different proportions of CBC and RSC tumor cells with higher RSC enrichment correlating with more stromal cells in the TME.

In this study we address these unknowns using analysis of single cell RNA-sequencing (scRNA-seq) datasets from CRC patients and performing scRNA-seq of heterogeneous SW480 tumor xenografts. We report that the binary classification scheme of RSC versus CBC is highly relevant to patient CRC and that patient tumors containing both populations can be modeled using SW480 CRC xenografting. The property of heterogeneity is intrinsic to the SW480 culture as it is comprised of RSC and CBC cancer stem cells with identical transforming genetic mutations (*APC, TP53* and *KRAS)*, but transcriptionally distinct gene programs. We present a method to isolate and separately xenograft RSC and CBC subtypes from the SW480 cell line and show that these subtypes exert striking, distinct influences on tumor progression and invasion. CBC-based tumors develop into large, highly proliferative, undifferentiated, aggressive tumors that invade and extensively wound neighboring epithelial tissue. CBC tumors are stroma poor overall, but the resident population of cancer associated fibroblasts (CAFs) is predominantly of the mature cancer-promoting phenotype and is highly prognostic for human CRC. The innate immune compartment has a large presence of plasmacytoid dendritic cells (pDCs). In contrast, RSC-based tumors are stroma-rich and differentiated with a *Wnt5a*- expressing environment encapsulated by a thick collagen matrix. RSC tumors exert a different signature of injury-resolution in neighboring mucosae with genes activated in epithelial enterocytes that promote wound healing. The innate immune compartment of RSC tumors is expanded with the presence of mast cells expressing genes for granzymes, serotonin production, and genes normally expressed in T lymphocytes.

Importantly, each subtype shares significant alignment with distinct human colorectal cancer CRIS molecular classifications and the YAP^ON^/YAP^OFF^ classification structure, and furthermore analysis of scRNA-seq datasets from patient tumors confirm that the RSC/CBC admixture model is highly relevant. Our study unifies multiple classifications of human CRC and demonstrates through subtype co-existence in a single culture as well as in patient biopsies how stable co-variants direct unique contributions to tumor phenotype.

## Results

### Binary patterns of CRC stem cell heterogeneity

The patterns of heterogeneity in xenografted CRC tumors that we and others have previously reported suggests that distinct cellular states might be a fundamental property reflected in the transcriptome of individual tumor cells^13–15^. To determine if this is the case, we performed scRNA-seq of xenografted SW480 subcutaneous tumors developed in NSG mice (Figure S1). Mouse and human cells sorted from the tumors that passed quality control filtering were clustered and visualized by uniform manifold approximation and projection (UMAP).

Analysis of human tumor cells revealed two distinct populations (Subtype1, Subtype2), each with a respective cycling population and differentially expressed genes (DEGs) with respect to the Wnt signaling pathway, intestinal inflammation and regeneration (Figure 1B-C; Figure S1E- G). The more abundant Subtype 1 cells expressed a Wnt target gene signature enriched in CBC intestinal stem cells (Figure 1C; *LGR5, SOX4, PROX1, FGFR4*) and we therefore assigned this subtype to be the LGR5+/Wnt^HI^ CBCs. Subtype 2 exhibited strong expression to a Fetal- Wounding and Regeneration signature and inflammatory genes (Figure 1C; *WNT5A, TACSTD2, CLU, GSN, IFITM1*) suggesting that these cells are the RSC subtype described by Vazquez et al.^7^. Interestingly, of the two xenograft replicates, the tumor with a greater abundance of RSC- Subtype 2 cells contained almost double the stromal compartment suggesting that the two subtypes might correlate with different tumor microenvironments (Figure 1D).

We hypothesized that CBC and RSC signatures delineate tumor heterogeneity in CRC but that the prevalence of one or the other subtype correlates with TME compositions. To test this hypothesis, we utilized publicly available scRNA-seq datasets from 29 CRC primary tumors (Figure 2A-C)^21^. We found that while many patients exhibited intra-tumor heterogeneity of both CBC and RSC phenotypes aligning with our SW480 Subtype 1 and Subtype 2 signatures, respectively, the predominant subtype signature in the tumor cells of each patient indeed correlated with differences in TME composition (Figure 2D-E). For instance, patient tumors that as a population scored more highly for the SW480 Subtype 1/CBC signature were predominantly composed of epithelial tumor cells (i.e. CBC-enriched patients; SMC01, SMC09, SMC16, SMC18, SMC21, SMC22), while patients that scored higher for the SW480 Subtype 2/RSC signature contained 5-7-fold more stromal and myeloid cells in their TME (i.e. RSC-enriched patients; SMC03, SMC14, SMC17, SMC20; Figure 2D-E; Figure S2). Interestingly, the significant differences in immune cell populations of the TME are among myeloid populations with no significant differences in the overall abundance of T and B lymphocytes or NK cells between tumor subtypes. We also confirmed that the additional CBC and RSC markers (Figure 1A) that are related to inflammation and regeneration (*TACSTD2,* Fetal-Wounding Signature) in RSC cells and proliferation (*PCNA, MKI67*) in CBC cells similarly distinguish the two cancer stem cell populations in patients (Figure S3).

**Figure 2:**
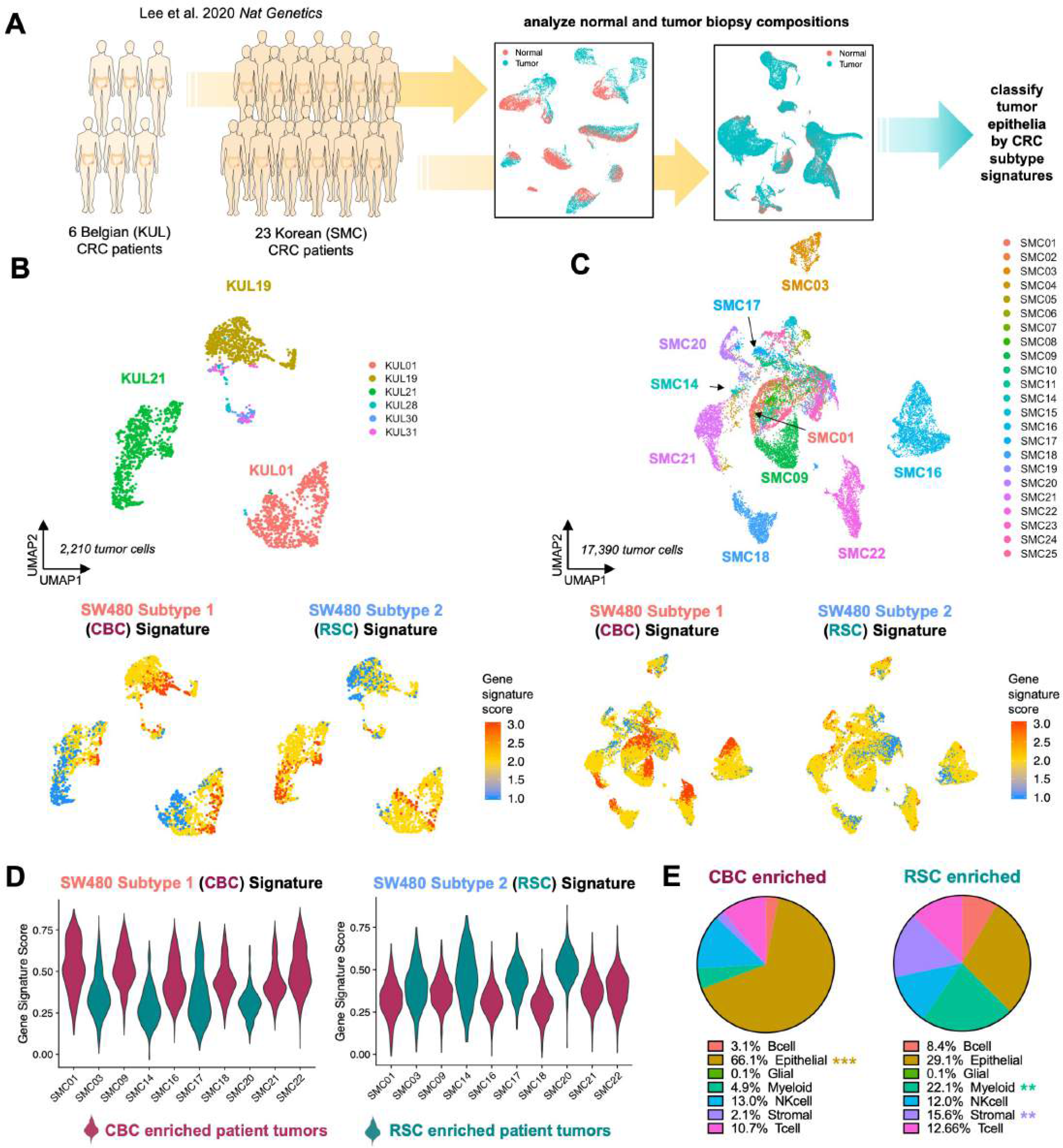
CBC and RSC cancer stem cell signatures link CRC patient-specific patterns of intra-tumor heterogeneity to distinct tumor microenvironment compositions. A: scRNA-seq analysis of two CRC patient cohorts from Lee et al., 2020. B,C: UMAPs of tumor cells from the Belgian (KUL) and Korean (SMC) CRC cohorts, with gene signature plots of SW480-derived CBC and RSC expression signatures. D: Violin plots showing the distribution of CBC- and RSC-enriched signatures in the population of the tumor epithelial cells. E: Pie charts illustrate the tumor microenvironmental compositions associated with CBC- and RSC­ enriched tumors from the SMC patients in D. P values=**< 0.01, *** < 0.001 (Student’s t test).

### SW480 cancer cell subtypes model binary classes of CRC

Analysis of the DEG lists for each subtype enabled the identification of *Roundabout homolog 1, ROBO1*, as a specific biomarker of SW480-CBC cells and *Epidermal Growth Factor receptor, EGFR*, as a biomarker of SW480-RSC cells (Figure S1H). Using antibodies to these receptors for FACS-based identification and separation of the two subtypes we identified distinct cellular morphologies of ROBO1± sorted cells that persisted across passages and single cell clonal derivation (Figure S4). Subtype 1/CBC cells exhibited a spherical “Rounded” morphology and Subtype 2/RSC cultures were a mixture of strongly adherent cells of cuboidal and mesenchymal morphologies (Figure S4). These distinct Rounded and Adherent morphologies of cells in the SW480 cultures are consistent with previous reports suggesting that the heterogeneity in the SW480 cell line is intrinsic and stable^22–24^. We confirmed this through genomic analysis: both subtypes carry identical oncogenic mutations previously identified in this patient (*APC/KRAS/TP53*) and identical Short Tandem Repeat profiles (Figure S5; Supplemental Data 1).

Bulk RNA sequencing analysis of freshly sorted ROBO1+/CBC and ROBO1-/RSC cells identified thousands of significant DEGs enriched in each subtype, including *LGR5*, which was 100-fold upregulated in the CBC subtype (Figure S6; Supplemental Data 2). Gene Set Enrichment Analysis (GSEA) confirmed significantly enriched gene signatures in the RSC subtype including the intestinal Fetal-like Wounding “Regeneration” signature^8,9^, YAP target genes^25,26^, and inflammatory interferon and STAT3^27^ signaling pathways. The CBC subtype is highly enriched for MYC signatures as well as proliferative Duodenal Transit Amplifying (TA) cells^28^. Their metabolic profiles are also drastically different with RSC cells expressing glycolytic and hypoxic gene signatures whereas CBC cells express a strong oxidative phosphorylation gene program, signatures that align with observations of their metabolism in culture. GSEA analysis also confirmed each subtype significantly aligns to different human CRIS molecular subtypes with CBC cells matching the crypt-bottom associated CRIS-E subtype, while the RSC subtype matches the two inflammatory CRIS-A and CRIS-B signatures as well as the EGFR+ CRIS-C subtype (Figure S6).

### RSC and CBC subtypes create divergent tumor landscapes

Given that enrichment of RSC or CBC signatures correlated with TME compositional differences in patients, we hypothesized that each SW480 subtype might direct the formation of distinct tumor initiation potentials and phenotypes. We tested this hypothesis via subcutaneous (SubQ) xenografting of single-cell clonally derived cultures. Limiting dilution analysis revealed that while CBC and RSC tumors were drastically different in size, there was no difference in tumor initiating potential (Figure S7). RSC xenografts developed as small, vascular-rich tumors with regions of ossified extracellular matrix (ECM), confirmed by Alizarin Red staining (Figure 3A-C; Figures S8-9). In contrast, CBC cells created much larger tumors (>10-fold) with a highly necrotic TME and adipocyte rich core, confirmed by BODIPY staining (Figure 3A-B; Figure S8).

**Figure 3:**
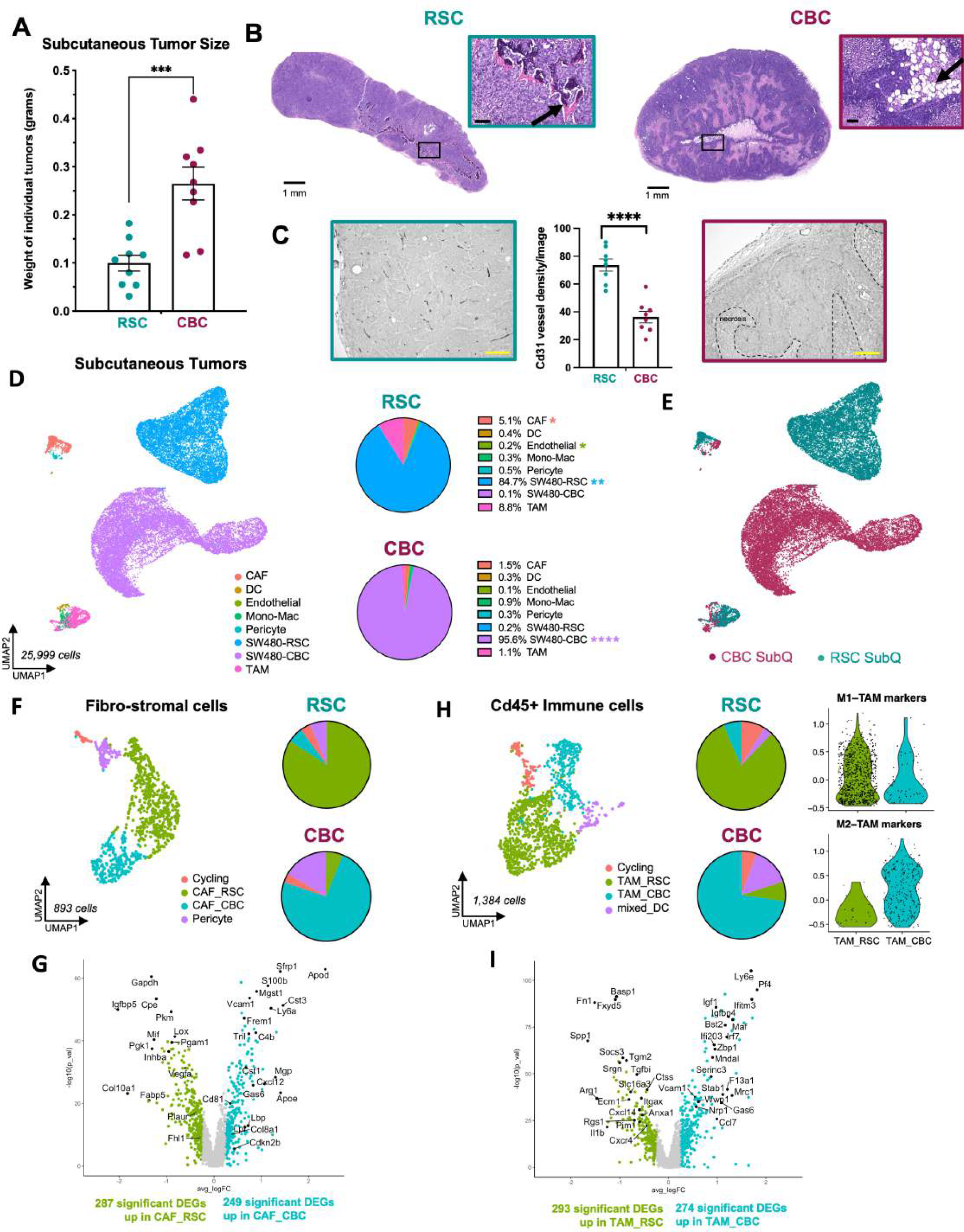
CBC and RSC subtypes generate distinct tumor microenvironments in subcutaneous xenograft tumors. A: Average size of subcutaneous tumors harvested after 4 weeks *in vivo* with representative images. B: H&E staining of RSC and CBC subcutaneous tumors, with arrows pointing out TME features of ossification (RSC) and adipocytes (CBC), scale= 1mm and 100µm (inset). C: Vasculature density as measured by Cd31 immunohistochemistry in subcutaneous RSC (left) and CBC (right) tumors, scale = 200µm. D: UMAP clustering of all mouse TME and human tumor cells from RSC and CBC subcutaneous xenografts with average proportions of cell types, n=2. E: UMAP clustering of all cells from the subcutaneous (SubQ) xenografts colored by subtype-specific xenograft. F: UMAP and pie chart of mouse stroma with cancer-associated fibroblasts (CAFs) colored by enrichment in tumor subtype-specific xenografts. G: Volcano plot of differentially expressed genes in mouse CAF populations enriched in RSC tumors (CAF_RSC) versus CBC tumors (CAF_CBC). H: UMAP and pie chart of mouse tumor-associated macrophages (TAMs) colored by tumor subtype-specific enrichment. I: Volcano plot of differentially expressed genes in mouse TAMs enriched in RSC tumors (TAM_RSC) and CBC tumors (TAM_CBC). P values=*< 0.05, ** < 0.01, **** < 0.0001 (Student’s ttest).

Given these differences in histology and tumor size, we performed scRNA-seq to define their TME at high resolution. CBC and RSC tumors were co-developed in individual mice by left and right rear flank injections (Figure S10A). Tumors were excised and halved for simultaneous H&E staining and single cell dissociation for scRNA-seq. Sequencing data was analyzed after mapping to both human and mouse genomes to distinguish human cancer cells from mouse host cells (Figure 3D, Figure S10B-D). CBC and RSC tumor cells remained distinct populations and expressed the same marker genes as when in mixed SW480 xenografts (Figure 3E; Figure S10E-G). However, we identified stark differences in the abundance and transcriptional signatures of stromal and immune populations.

The overall amount of TME in CBC tumors was 3-fold less than RSC SubQ tumors (5% vs 15%; Figure 3D). Immune and mesenchymal fibro-stromal cells were remarkably transcriptionally distinct with populations of tumor associated macrophages (TAMs) and cancer associated fibroblasts (CAFs) more abundant in RSC tumors (Figure 3E-I). Pericytes also formed distinct populations associated with tumor subtype as evidenced by differential expression of bone associated genes (*Acta2, Ctgf)* and fat associated genes (*Fap*, *Lpl, Pparg*) in RSC and CBC tumors, respectively (Figure S11D). These distinguishing features align with observations of bone and fat ECM staining in the two tumor subtypes (Alizarin Red and BODIPY, respectively; Figure S8), and implicate pericytes as a potential mesenchymal progenitor that mediates crosstalk between tumor and stroma.

*Cd45*+ immune cells could be clustered into multiple subgroups including subpopulations of TAMs and Tissue Resident Macrophages (TRM). The fast-growing CBC SubQ tumors contained nearly all of the tumor-promoting M2 subtype of TAMs (TAM_M2) and TRMs expressing interferon stimulated genes (ISGs; TRM_Inflam; Figure S12). In contrast, RSC SubQ tumors contained a greater complexity of transcriptional states amongst *Spp1+* M1- TAMs with subclusters expressing distinct sets of inflammatory biomarkers (Figure S12).

Overall, the inflammatory M1-TAM and *Spp1*+ populations were nearly exclusive to RSC tumors, while the tumor-promoting M2-TAM population was near exclusive to CBC tumors. Comparing all macrophages from RSC tumors (TAM_RSC) and CBC tumors (TAM_CBC), we identified roughly 300 DEGs with TAM_RSC expressing ECM, apoptosis and chemokine gene programs, while TAM_CBC highly express ISGs, anti-inflammatory M2 biomarkers and regulators of necrosis (Figure 3H-I). Dendritic cells (DCs) were a minor population in the SubQ setting, but we identified multiple subpopulations including an enrichment of mature, migratory DCs (migDC) and plasmacytoid DCs (pDC) in CBC tumors (Figure S12E).

### Orthotopic xenografts develop a more diverse, complex tumor microenvironment

Comparing tumor subtypes in an orthotopic setting, we again observed that injection of CBC cells created large, stroma-poor, necrotic tumors that were highly aggressive with destructive invasion into the epithelium, lymph, and muscle compartments (Figure 4A-C; Figure S13). RSC orthotopic tumors were again smaller yet rich in stroma and densely wrapped with collagen fibers (indicated by Trichrome staining; Figure 4A-C). Pathological scoring of H&E as well as smooth muscle actin immunohistochemistry (IHC) staining of both tumor subtypes characterized CBC tumors as having an undifferentiated morphology, while RSC cancer cells were observed to organize into elongated rosette clusters, bounded by ECM proteins - reminiscent of quasi-differentiated epithelial crypt structures (Figure 4C).

**Figure 4:**
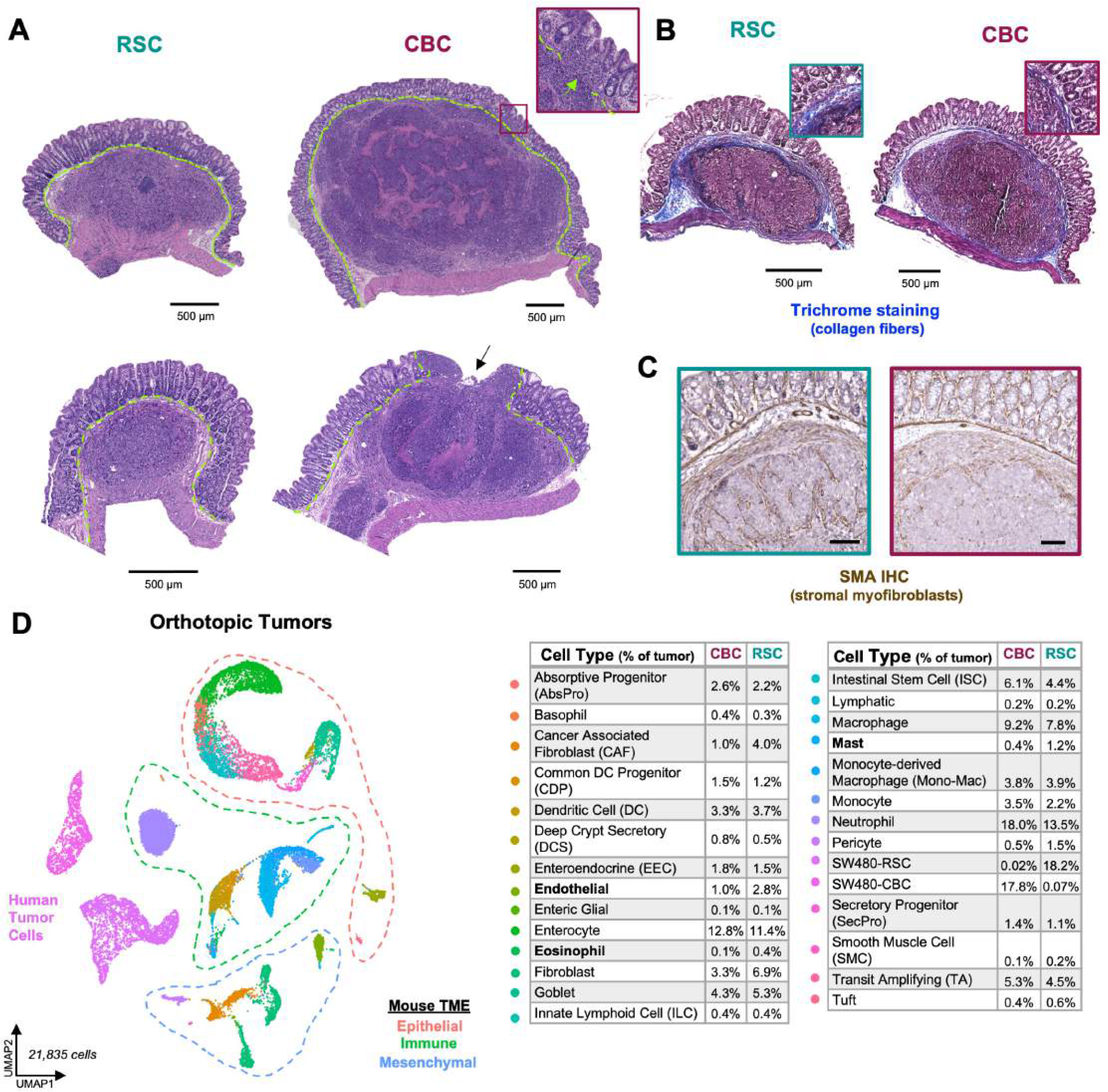
Orthotopic xenograft tumors host a more complex and diverse TME. A: H&E staining of RSC and CBC orthotopic tumors, with arrows pointing to tumor cell invasion through the muscularis mucosae (green dashed line), scale= S0Oµm. 8: Trichrome staining of orthotopic tumors, collagen fibers are stained blue, scale = S00µm. C: lmmunohistochemistry of SMA protein in orthotopic RSC (left) and CBC (right) tumors, scale = 200µm. D: UMAP clustering of all mouse and human tumor cells captured in orthotopic xenografts with legend reporting the average cell type proportions in CBC and RSC tumors, n = 4 per subtype. Balded cell types denote significant TME differences between tumor subtypes; P values < 0.05 (Student’s t test).

While scRNA-seq of SubQ tumors revealed many TME differences associated with tumor subtype, TME content was relatively limited, especially for fast-growing CBC tumors (only ∼5% TME). We therefore investigated whether scRNA-seq of orthotopic tumors developed in the intestinal wall of the cecum, a more physiologic setting, would recapitulate the pro-inflammatory versus immune-depleted TME features observed in SubQ tumors. Tumors harvested for scRNA-seq analysis (n = 4 mice each subtype) again showed each tumor subtype as distinct populations, and the expected TME cell types of surrounding murine tissues were identified (Figure 4D; Figure S14). In contrast to the SubQ setting, human tumor cells comprised only ∼18% of the orthotopic TME (Figure S14C). A rich variety of immune cell types not captured in SubQ tumors (e.g neutrophils, mast cells, eosinophils, basophils) were identified including an expansion of innate progenitors and dendritic cell subtypes (Figure 4D; Figure S14). Additionally, neighboring epithelial populations from normal mucosa that encapsulate the developing tumors were also captured.

Analysis of the TME in our orthotopic model revealed several important distinctions between tumor subtypes with the largest difference comprising a 4-fold greater abundance of cancer associated fibroblasts (CAFs) in RSC tumors, with significant enrichment of an intermediary CAF_1 phenotype (Figure 5; Figure S15B). UMAP clustering of *Col1a1*+ fibroblasts grouped crypt associated and cancer associated fibroblast populations which were further distinguished using marker genes expressed by intestinal fibroblasts(Figure S15C-D)^29^. The CAF marker *Thy1*^30^ revealed two transitional but distinct CAF populations wherein the CAF_2 population comprised the majority population in the aggressive CBC tumors (75%) versus its presence as a minority population in RSC tumors (40%; Figure 5B-C; Figure S15E). CAF_2 cells express high levels of *Fap* and *Tgfb1* as well as gene signatures associated with angiogenesis, cell migration, and wounding (Figure 5D; Figure S15E). Notably, high level expression of the CAF_2 signature is prognostic for worse survival in CRC patients (Figure 5E), a measure that highlights the relevance of this signature to human CRC. RSC tumors contain an enrichment of the CAF_1 population which expresses gene programs involved in tissue development and morphogenetic processes including *Wnt5a* (Figure 5D; Figure S16), signatures that are ontologically congruent with the Fetal-like “Regeneration” signature expressed by the RSC subtype. Our dataset also contained a large quantity of normal intestinal fibroblasts which enabled us, via gene signature similarities, to predict the origins of the poor-prognosis CAF_2 population to fibroblasts that surround the stem cell zone at the bottom of the intestinal crypt (i.e. CBF_2), and the regenerative CAF_1 population to fibroblasts that associate with the top half of the crypt where epithelial cells are differentiating (i.e. CTF; see Discussion and Figures S16-17).

**Figure 5.**
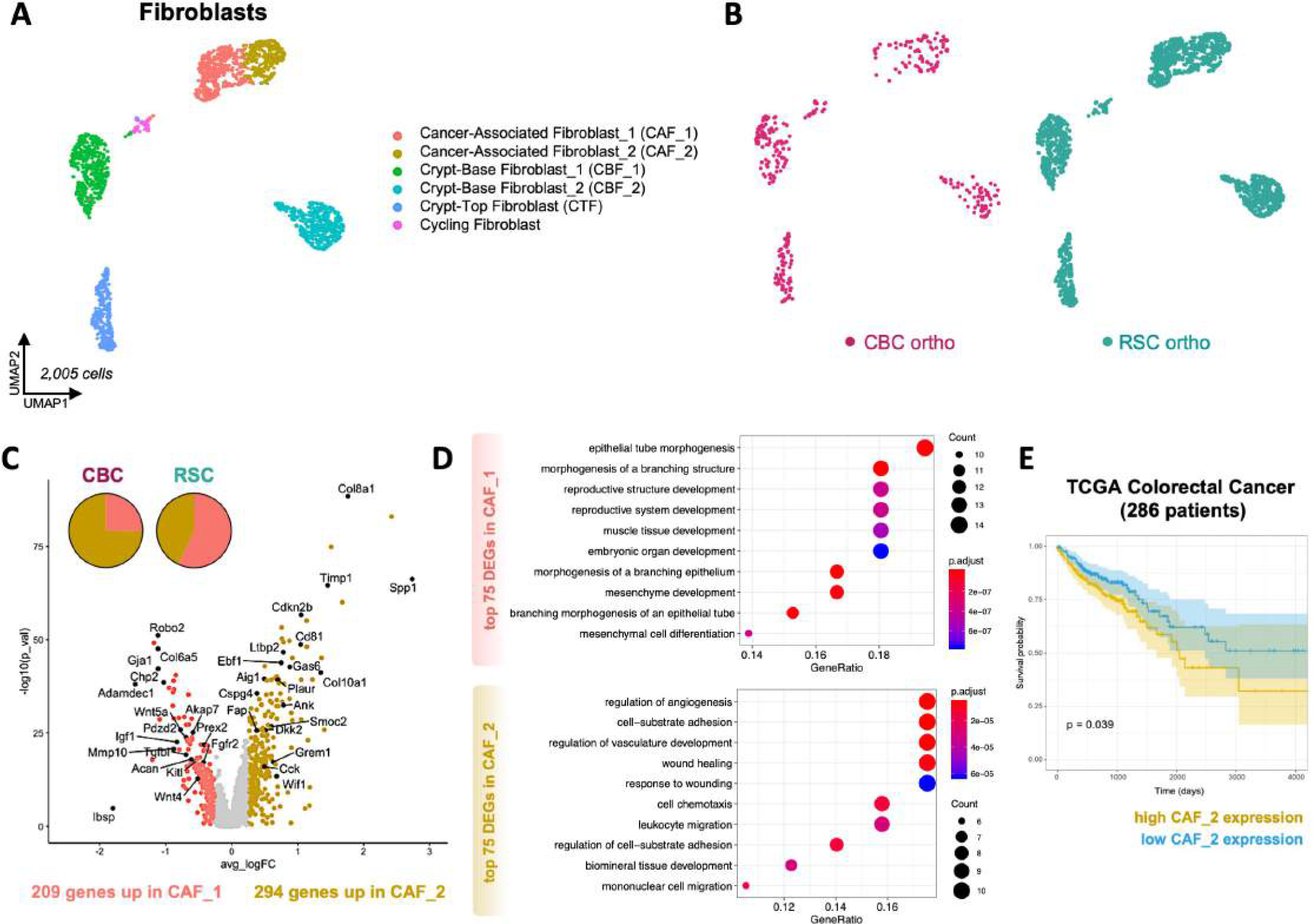
CBC orthotopic tumors are enriched in a CAF subtype that correlates with worse patient prognosis. A: UMAP clustering of mouse normal crypt-associated and cancer-associated fibroblast (CAF) populations in orthotopic xenografts. B: UMAP clustering of all fibroblasts colored by tumor subtype-specific xenograft. C: Volcano plot of differentially expressed genes (DEGs) in CAF subtype populations (CAF_1 and CAF_2). D: DotPlot showing functional enrichment results associated with CAF_1 and CAF_2 top DEGs. E: Survival curve of TCGA colorectal cancer patients (n = 286) that express low and high expression signatures of CAF_2 marker genes.

The immune landscape in orthotopic xenografts was a more diverse and abundant presence compared to the SubQ TME (Figure 6; Figure S18A). We identified multiple types of granulocytes (neutrophils, mast cells, basophils, eosinophils) that were absent from the SubQ setting, and these populations were some of the most significantly different between the two tumor subtypes including the relative abundances of mast cells and eosinophils (Figure 6A; Figure S19C). For example, one of the three subpopulations of mast cells was completely absent in CBC tumors (RSC-specific Mast cell (RSC-MC)). This population exhibited strong and distinct expression of secreted proteases (*Mcpt9*, *Gzma, Gzmb*), serotonin synthesis enzymes (*Tph1, Ddc*), and adhesion complexes specific for E-cadherin *(Cdh1:Jup, Itgae:Itgb7*), suggesting that this population could play a unique role in tumor-TME crosstalk (Figure 6C-D; Figure S19D-E). RSC tumors also contained a greater number of tumor-associated mast cells (TAMC)^31^ that express *Vegfa* (Figure S19D-E).

**Figure 6:**
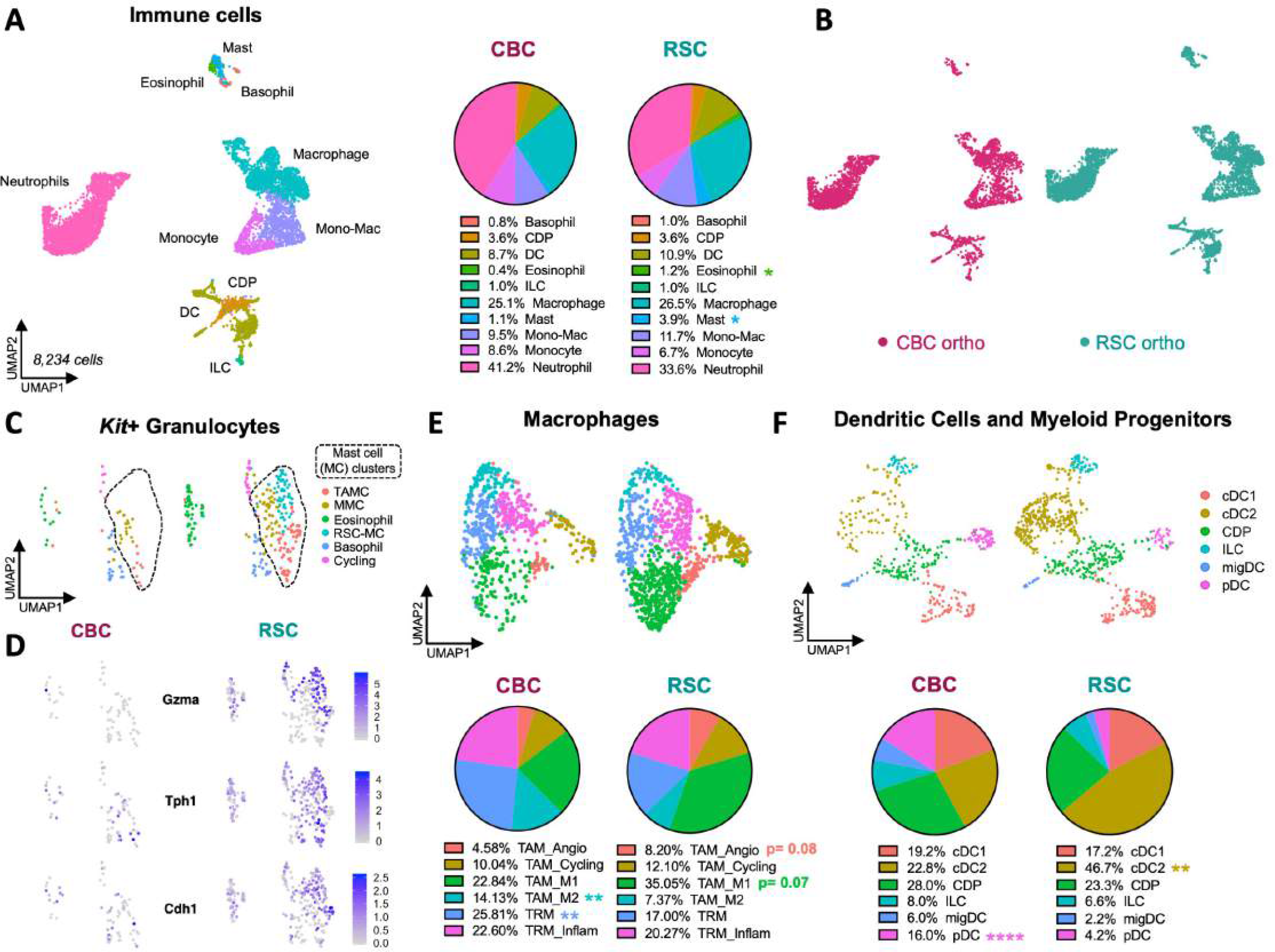
Pro-inflammatory and immune-suppressive cell types distinguish the TME of RSC and CBC orthotopic tumors. A: UMAP clustering of Cd45+ immune cells in orthotopic xenografts with pie charts showing cell type proportions in each tumor subtype. B: UMAP clustering of immune cells colored by tumor subtype-specific xenograft. C: UMAP clustering of *Kit+* granulocytes identifies mast, basophil, and eosinophil populations. D: Gene expression plots predominantly expressed by mast cells in RSC tumors (RSC-MC). E: UMAP clustering of tumor-associated (TAM) and tissue-resident (TRM) macrophages, with pie charts showing proportional compositions of macrophage states. F: UMAP clustering of dendritic cells (DCs) and myeloid progenitor cell types, with pie charts showing proportional differences of DC heterogeneity. P values=*< 0.05, ** < 0.01, **** < 0.0001 (Student’s t test).

In addition to tumor-specific signatures in *Kit*+ granulocytes, we identified significant differences in the prevalence of *Adgre1*+ macrophages (Figure S20). Both tumor subtypes contained these populations, but the pro-inflammatory TAM_M1 state was more prevalent in RSC tumors while the tumor-promoting TAM_M2 state was two-fold more abundant in CBC tumors (Figure 6E; Figure S20E). The opposing proportions of tumor-promoting and tumor-suppressing macrophage populations align with the overall tumor phenotypes where the smaller, slower growing RSC tumors have a higher proportion of pro-inflammatory M1 macrophages and the faster growing, invasive CBC tumors have a higher proportion of tumor-promoting M2 macrophages.

Despite NSG mice lacking an adaptive immune system, an examination of the dendritic cell (DC) population, which was more abundant in the orthotopic setting, allowed us to investigate further differences between tumor subtypes. DCs are a heterogeneous cell type with many different functions that are known to bridge innate and adaptive immune responses. We identified six DC populations and progenitor states (Figure 6F), with higher resolution clustering resolving into nine subpopulations (Figures S21-22). Conventional DCs (cDCs) were the most diverse DC subtype with the cDC2 subpopulation enriched in RSC tumors (Figure 6F). This population expresses pro-inflammatory genes and enzymes central to prostaglandin synthesis (Figure S22; *Nr4a3, Ptgs2, Hilpda*)^32^. We also identified an innate lymphoid progenitor (ILC) population that expresses NK cell (*Ncr1*) and T-cell markers (Figure S21; *Pdcd1, Tox, Il2rb, Tcf7*). The hybrid co-expression of innate and adaptive associated genes in an ILC population suggests that this population is a bona fide connector of the two arms of the immune system in NSG mice^33^. Taken together, RSC tumors have a much greater abundance of innate cell types associated with inflammation, prostaglandin synthesis and serotonin signaling. In contrast, immune-suppressive plasmacytoid DCs (pDCs), marked by *Siglech, Bst2, and Bcl11a*^34^, were 4- fold more abundant in the proliferative, invasive CBC tumors (Figure 6F; Figure S21). Previous studies have shown pDCs can activate T-regulatory cells, a potential contribution to the immune-suppressive environment of CBC tumors^35,36^.

### Differential wounding patterns in mucosal epithelial cells

The orthotopic TME is encapsulated by mouse intestinal epithelium that is co-excised during tumor harvest. Our scRNA-seq analysis identified all the known stem cell, secretory, and absorptive populations from the mucosal epithelia (Figure 7A; Figure S23A-B). As both tumor subtypes differed in their degree of invasion into the normal mucosa, we also discovered marked differences in the way each subtype wounded intestinal epithelial tissue. For instance, we could ascribe distinct patterns of wounding associated with tumor-induced injury and healing, and we developed a novel signature for damaged mouse epithelia in both tumor subtypes (Figure S24; *Gadd45a+Plaur+Sprr1a+F3.1*+*Gm26825-*). Interestingly, we identified distinct signatures and abundances of wounded epithelial cells in each tumor subtype that signify early versus late stages of tumor-induced damage. For example, the invasive CBC tumors extensively damage neighboring mucosae such that the mature *Krt20*+ enterocyte (Ent_mature) population was depleted and replaced by a population of enterocytes expressing interferon stimulated genes (Ent_ISG; Figure 7A-C; Figure S23C). The mucosae overlying RSC tumors incurred less damage, containing both normal enterocytes (Ent_mature) and a population expressing the pro-resolving wound healing factor *Anxa1* (Ent_Anxa1; Figure 7A-B; Figure S23)^37^.

**Figure 7:**
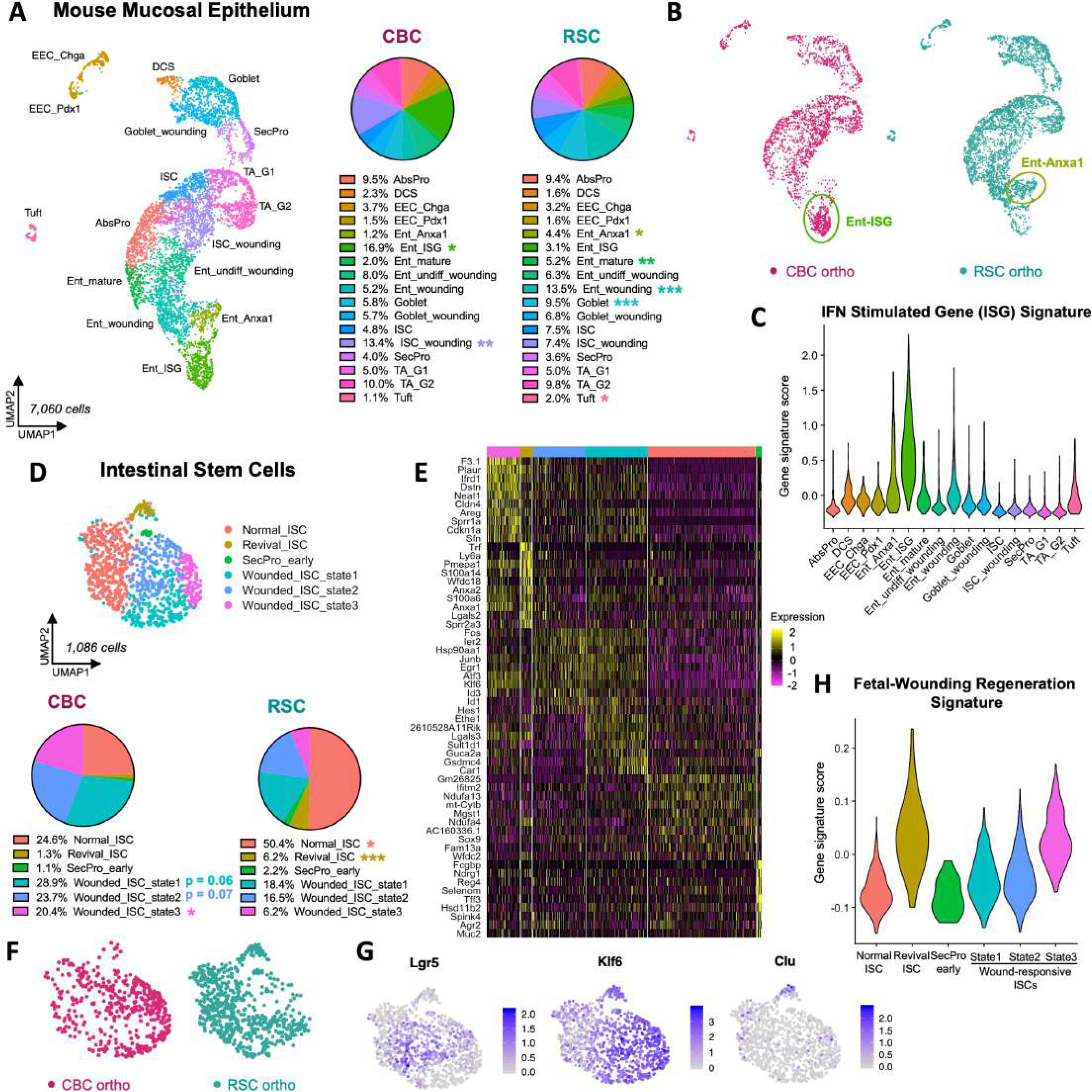
Mouse epithelial cells exhibit subtype-specific patterns of wounding in CBC and RSC orthotopic tumors. A: UMAP clustering of *Epcam+* mouse mucosal epithelium in orthotopic xenografts with pie charts showing proportional composition of epithelial cell types. B: UMAP clustering of mouse epithelium colored by tumor subtype xenograft; two different populations of wounded enterocytes (Ent-lSG and Ent-Anxa1) are indicated. C: Violin plot of Interferon Stimulating Gene (ISG) Signature scoring across mouse epithelial cell types. D: UMAP clustering of mouse intestinal stem cell (ISC) states with pie charts showing proportional composition between subtype-specific xenografts. E: Heatmap of marker genes expressed by normal and wounded ISC states. F: UMAP clustering of ISCs colored by subtype-specific xenograft. G: Gene expression plots of biomarkers of revival stem cell *(C/u)* and wounded ISC *(K/f6)* populations. H: Violin plot scoring of the Fetal-Wounding Regeneration Signature amongst ISC states. P values = * < 0.05, ** < 0.01, *** < 0.001 (Student’s t test).

These effects of differential bystander wounding by the tumors were particularly evident in the murine intestinal stem cell (ISC) population with multiple subpopulations of stem cells expressing unique signatures of damage (Figure 7D-F). In the less damaged early stage of wounding as observed with RSC tumors, normal mouse ISCs comprised 50% of the stem cell population but in more damaged later stages of wounding from CBC tumors, only 25% of normal ISCs were detected (Figure 7D). *Klf6* is a transcription factor and immediate-early injury response gene, and its expression is triggered in wounded ISCs^4,38^. *Klf6* expression labeled a large proportion of the ISC population, including the *Lgr5*+ population that was uniquely present in CBC tumors (Figure 7G). This co-expression pattern suggests that the *Lgr5*+ stem cell population in the mucosae near CBC tumors is in fact, mostly wounded and there are very few “normal”, homeostatic ISCs in and around the invading tumor. Indeed, the majority of ISCs (75%) isolated from the CBC tumor samples were divided between three transcriptional wounding states with Wounded_ISC_state3 being the most highly enriched compared to RSC tumors. Another important stem cell marker for injury is the *Clu*+ “Revival Stem Cell” population (Revival_ISC; Figure 7G)^18^. This small, quiescent population in normal crypts is triggered by injury to proliferate and replace the ISC compartment. We found this population was significantly depleted in the CBC tumors compared to RSC tumors (Figure 7D). Activated *Clu*+ Revival_ISC cells had the highest enrichment of the Fetal-Wounding “Regeneration” gene signature, and curiously, the Wounded_ISC_state3 also scored highly for this signature (Figure 7H). This gene signature alignment suggested that perhaps the absence of Revival_ISC in CBC tumors was due to their activation, proliferation, and constitution of the Wounded_ISC_state3 population.

Taken together, the significant enrichment of Wounded_ISC_state1,2,3 populations in CBC tumors suggests a greater degree of wounding, and perhaps even a more prolonged, chronic state of wounding since the Revival_ISC population is nearly depleted in its entirety.

## Discussion

The cellular basis for tumor heterogeneity and its consequences in the tumor microenvironment are critical, fundamental questions for understanding how tumors develop and for developing better treatments. While several of the classification structures that define heterogeneity in CRC were derived from bulk RNA sequencing of patient samples, these studies lacked single cell resolution to determine the true basis of intra-tumor heterogeneity. Here we close that important gap by utilizing scRNA-seq of CRC patient biopsies together with xenograft models to provide an in-depth transcriptomic analysis of two colorectal cancer stem cell subtypes: Regenerative Stem Cells (RSCs) and Crypt Base Columnar cells (CBCs). We define how each subtype orchestrates its tumor microenvironment at the single cell level, underscoring that these differences were identified even when tumors were grown in the same mouse. Our strategy to isolate RSCs and CBCs from the heterogeneous SW480 cell line revealed the these subtypes align with, and unify, two proposed binary classifications of CRC: the YAP^ON^/YAP^OFF^ and the RSC/CBC classification models^5,7^. The transcriptomic and cellular signatures derived from the xenograft models were useful in determining that CBCs and RSCs co-exist in most CRC and can therefore account for much of the naturally occurring intra-tumor heterogeneity in CRC patient tumors. Given the intriguing co-existence of these two subtypes over decades in culture, we predict that they provide mutual benefit for survival and that their coexistence in patient tumors in varying proportions dictates the overall tumor phenotype. For example, Belgian patient KUL19 whose tumor scored higher for the CBC signature contained a large population of immune-suppressive pDCs and fewer CAFs which have gene signatures associated with the poor-prognosis CAF_2 phenotype (Figure S25).

To our knowledge, our study is the first to link CRC stem cell subtypes with unique TME signatures and phenotypes: the RSC subtype promotes a differentiated, highly vascularized, pro-inflammatory environment and CBC tumors are stroma-poor yet faster growing and more aggressive. The immune-depleted CBC tumors also contain cancer associated fibroblasts (CAFs) that align with CAF signatures of poor-prognosis patients (CAF_2; Figure 5E). The enhanced invasive phenotype also induces a unique interferon-driven bystander effect on neighboring epithelial mucosae, as well as a heightened injury signature in the normal intestinal stem cell population. Perhaps prolonged injury from an invading tumor front might subject normal crypts and stem cells to a risk of Field Cancerization.

### Single cell resolution of the tumor microenvironment

To our knowledge, we provide one of the first, in-depth single cell atlases of an NSG mouse, showcasing all the innate immune populations present in xenografted tumors, with surprisingly, some populations expressing genes normally associated with adaptive NK and T-cell functions (e.g. granzymes, PD-1/*Pdcd1*, *Tox*, *Ncr1*). We find that there are important and significant tumor-specific differences in the innate immune cell and fibroblast compartments.

Concordantly, it is these compartments that were also the most significantly different between patient tumor subtypes.

Previous studies have suggested that a quantitative measure of the stromal abundance of CAFs is considered to be a negative prognostic indicator for CRC outcomes^3,39–41^. However, our tumor models highlight that CAF gene expression might be a stronger predictor. In both SubQ and orthotopic models, CAFs were 3.5-4-fold more abundant in the slower growing RSC tumors, making these tumors more fibrotic than the fast-growing, invasive CBC tumors.

Immunohistochemistry (IHC) staining for the myofibroblast biomarker SMA in RSC tumors identified CAFs surrounding clusters of tumor cells as if they were circumscribing quasi-crypt forming units, similar to regenerating epithelium, but managing only abortive attempts at gland formation (Figure 4C). Concordantly, the majority CAF population (CAF_1) in RSC tumors express developmental gene programs reminiscent of embryonic and/or regenerative fibroblasts which are necessary for the formation of intestinal crypts^42^. Even though CAFs were 4-fold less abundant in CBC tumors, the predominance of mature *Fap*+ CAFs (CAF_2) which express multiple anti-inflammatory secreted molecules (*Gas6, Tgfb1, Cxcl12*) was associated with significantly worse prognosis in CRC patients (Figure 5E). Therefore, our data suggests that the transcriptional identity of CAFs is more important for prognosis than the physical abundance. In addition to CAF signatures as prognostic indicators, our data provide novel predictions of their origin (Figures S16-17). The CAF_2 population, enriched in CBC tumors, shares patterns of gene expression with crypt-bottom CBF_2 fibroblasts. This suggests that CBC cells, which epitomize oncogenic homeostatic crypt stem cells, may be influencing CAF differentiation to retain crypt base niche expression signatures. In contrast, RSC tumors have a predominance of CAF_1 fibroblasts that co-express genes with CTFs, which are found at the top of crypts associated with mature, differentiated epithelia. Interestingly, *Wnt5a*, a well-known signal for inflammation, is highly expressed by CTFs, CAF_1 fibroblasts and RSC cancer cells (Figure S26A). Furthermore, we detected WNT5A expression in CAFs from RSC enriched CRC patient tumors (SMC14, SMC17, and SMC20) suggesting that this Wnt ligand is potentially an important and clinically relevant signal in the TME of RSC tumors (Figure S26B).

A study of the innate immune compartment of NSG mice allowed for important discoveries in three populations: mast cells (MCs), tumor associated macrophages (TAMs) and dendritic cells (DCs). We captured a large abundance of MCs in RSC tumors, with a unique population completely absent in CBC tumors that exclusively express genes associated with serotonin production (*Tph1, Ddc*), cytotoxic T-cells (*Gzma*) and tissue remodeling (*Mcpt9*; Figure 6D; Figure S19D). For example, these cells have the strongest signature for secreted proteases including *Gzma*, which is normally produced by cytotoxic T-cells, and *Mcpt9*, a β- chymase protease known to be upregulated during infection and tissue remodeling^43^. Because this population strongly expresses genes for serotonin production, we searched for expression patterns of serotonin receptors (Htr’s) and the uptake transporter *Slc6a4* which reduces environmental concentrations. We found that monocyte, macrophage, and DC populations in RSC tumors can respond to serotonin (*Htr2b*, *Htr7* expression; Figure S20D; Figure S22G).

However, there is no expression of *Slc6a4/SLC6A4* in any cell type in the tumor. This suggests that serotonin production by MCs is a long-lasting environmental signal that contributes to the overall inflammatory TME.

Additionally, we identified patterns of TAM polarization as another defining feature associated with CRC subtype. RSC tumors were enriched for TAM populations expressing inflammatory M1 signatures while CBC tumors were enriched for tumor-promoting M2-TAMs. Eosinophils were markedly depleted in CBC tumors compared to RSC eosinophils which express high levels of Osteopontin (*Spp1*) and the serotonin receptor *Htr7*, meaning that they are likely able to respond to serotonin-secreting MCs (Figure S19F). Finally, the contrasting roles of different DC populations enriched in each tumor subtype further promote the opposing pro-inflammatory (cDC2 cells in RSC tumors) versus anti-inflammatory phenotypes (pDCs in CBC tumors)^44,45^.

### Tumor associated wounding signatures in bystander epithelium

A fascinating aspect of this study was the discovery of different tumor-induced wounding patterns within the epithelial compartment including distinct transcriptional states of wounded stem cells. Using Monocle2 analysis to build cell differentiation trajectories, we observed unique patterns of wounding at different windows of “pseudotime” (Figure S23D). In this analysis, differentiating wounded populations lie closer to the predominantly injured ISCs at the beginning of pseudotime in CBC tumors. Whereas in RSC tumors these wounded differentiating populations are placed farther away from the largely uninjured ISC population. This suggests a wounding trajectory that develops over time as stem cells transition to wounded states. It is therefore possible that the different wounding patterns in the two tumor subtypes provide a model for the effects of early (RSC) versus later stages (CBC) where there is sustained, invasive damage exerted on neighboring, normal epithelia. Finally, we discovered that the lncRNA *Gm26825* is an exclusive biomarker of normal mouse epithelial cells as its expression disappears in wounded populations (Figure S24). This discovery enabled us to develop a novel “Wounding Signature” that broadly captures wounded cells of mouse intestinal epithelium (*Gadd45a*+, *Plaur+, Sprr1a+, F3.1+, Gm26825-*). We found that a human version of this signature (*GADD45A, PLAUR, SPRR1A*) was indeed upregulated in tumor epithelial cells and bordering biopsies of KUL patients compared to their normal epithelial counterpart (Figure S25C).

Developing a system to separately culture and study the two CRC stem cell subtypes that co-exist in the SW480 cell line offered a unique opportunity to define their transcriptomes and directly compare how each subtype orchestrates a different tumor microenvironment. Our discoveries of markedly distinct tumor phenotypes offer a new perspective on intra-tumor heterogeneity illustrating how different stem cell populations promote different hallmarks of cancer. We find that the CBC and RSC subtypes and the tumors they create remain connected to the functions of their normal stem cell counterparts, either the proliferative capacities of CBCs, or the wound healing properties of regenerative Revival Stem Cells (revSCs).

Understanding the mechanisms underlying these inherent stem cell properties of plasticity and differentiation in each subtype will be important for identifying drug sensitivities and preventing the emergence of drug resistance.

### Limitations of the study

It is important to note two limitations of our study. First is the use of immune compromised mice that lack an adaptive immune system. While we have identified multiple innate immune cell types (e.g. M2-TAMs and plasmacytoid DCs) that are immune suppressive in other mouse and human immune competent models, we cannot account for any direct immune-mediated effects in our xenograft model as the adaptive compartment has important influences on innate populations and the overall status of immunity in and around tumors.

Interestingly, our analysis of the TME of CRC patients (Figure 2E) determined that there are significant differences in abundance of innate cell populations such as myeloid cell types but no apparent differences in the abundance of adaptive cell populations: T and B lymphocytes or NK cells. Therefore, while the use of NSG mice in our study is a limitation, the differences we identified in the innate compartment share some similarities to the tumor subtypes in immune-competent patients.

An additional limitation is that our study discovers CRC cancer stem cell subtypes from a cell line. Given that almost every human patient tumor that we examined contained RSC and CBC-like cells (Figure 2), we predict these two subtypes will be detected often in KRAS/TP53/APC MSS tumors. Here we exploited the heterogeneity innate to the SW480 cell line as a useful tool to identify and define two CRC cancer stem cell subtypes. We determined that these cancer stem cells are stable and that they direct the formation of distinct tumor microenvironments. The TME signatures of those environments, such as the poor prognosis CAFs in CBC tumors and bystander wounding of epithelial tissue, should be useful in the analysis of patient tumors and immune-competent mouse models.

## Acknowledgements

We thank all members of the Waterman Lab and collaborators John Lowengrub, Arthur Lander, Peter Donovan, and Grant MacGregor for discussion of this project, and Harry Mangalam for bioinformatic support. We are grateful to Morgan Dragan, Adeela Syed, Jennifer Bates, Melanie Oakes, and Stephanie Hachey for their assistance with experiments. The authors acknowledge the support of the Chao Family Comprehensive Cancer Center Shared Resources: Optical Biology Center, Flow Cytometry Facility, Genomics Research & Technology Hub, and the Experimental Tissue Resource. This work was supported by the National Cancer Institute of the National Institutes of Health (P30CA062203, U54-CA217378, and R03CA223929 to M.L.W.).

The content is solely the responsibility of the authors and does not necessarily represent the official views of the National Institutes of Health. The material is based on work supported by the National Science Foundation (DGE-1839285 to L.H.). Research was supported by an Anti- Cancer Challenge research grant from the University of California, Irvine Chao Family Comprehensive Cancer Center.

## Author Contributions

M.L.W. and L.H. designed and supervised research and wrote the manuscript; L.H. performed all experiments and bioinformatic analyses; D.T. performed all mouse surgeries and xenograft injections; R.A.E. contributed to pathological analysis and maintained animal compliance; S.Y.P. quantified staining; K.N., S.Y.P., J.W., A.N.H., C.V. and M.M.S. assisted with bioinformatic analyses.

## Declaration of Interests

The authors declare no competing interests.

## Methods

### Material availability

Further information and requests for resources and reagents should be directed to and will be fulfilled by Marian L. Waterman.

### Data availability

Raw bulk RNA-seq datasets for ROBO1+ and ROBO1- sorted SW480 cells are available in the NCBI Gene Expression Omnibus (GEO) database using accession number GSE254891.

Original and processed scRNA-seq data from xenograft tumors are available under GEO accession number GSE254890. Script for processing of data can be provided upon request.

### Xenograft Models

All mouse experiments were approved by the University of California Institutional Animal Care and Use Committee (AUP-20-033). In brief, subcutaneous xenografted tumors were generated by injecting 1-2 million cells in 100µL of PBS into the back rear flanks (and shoulder flanks for limiting dilution experiments, with cells suspended in Matrigel) of male NSG mice using a 26 gauge needle and 1 cc syringes. Tumor inoculations were left to develop for 21-28 days, to a maximum diameter of 1.2cm, or until time of harvest. Excised tumors were measured and photographed before tissue processing. For orthotopic cecal xenografts, male NSG mice were placed under anesthesia and a surgical incision was made in the abdomen to visualize the cecum. Using a Hamilton syringe, 100-200K cells/10µL PBS was injected into the submucosal space of the cecum, performing 4-5 injections per side for a total of 8-10 inoculations per mouse cecum. The mouse was placed in a semi-reclined position and the outside cecal surface was rinsed with PBS before insertion back into the abdomen and suturing (to avoid seeding the abdominal cavity). Orthotopic inoculations were left to develop for 28-35 days before harvesting. Individual tumors excised from the cecum were measured and photographed, or the whole cecum tissue was linearized and rolled into a “swiss roll” before tissue processing.

### Cell lines and culturing

SW480 and SW620 cell lines were purchased from ATCC in 2019 and cultured in T-75 flasks (Falcon) with DMEM-complete (DMEM-C: DMEM High Glucose (Hyclone), 10% FBS (Atlas Biologicals), 1X L-glutamine (Gibco), and 1X Antibiotic-Antimycotic Solution (Corning)). Cells were passaged every 3-5 days using trypsin-EDTA by splitting 1:5 to maintain sub-confluency. All experiments were performed within the first 12 passages of a new stock vial from ATCC and/or clonal derivation. Cell cultures were mycoplasma tested with the MycoAlert Mycoplasma Detection Kit (Lonza) every 3-5 months. Confirmation of cell line identity was performed using short tandem repeat (STR) profiling by the University of Arizona Genetics Core and SNV mutations were identified by exome sequencing by the UCI Genomics Research & Technology Hub.

Passaging of ROBO1+ sorted Rounded/CBC cultures every 4-5 days is best done by incubating cells with 0.25% trypsin-EDTA (Gibco) for 1-2 min at RT/37°C, rigorously pipetting the cells with trypsin with a p1000 Pipetman (Gilson) to break up colonies and achieve a single cell suspension of the Rounded/CBC subtype. Suspensions are quickly quenched with DMEM-C and counted for concentration and viability. For subculturing experiments after FACS, growth medium and PBS washes were added back to the trypsinized cell suspension to account for floating Rounded/CBC cells (normally lost during vacuum aspiration of media/wash steps).

Passaging of Adherent/RSC cultures is achieved by incubating cells with 0.25% trypsin-EDTA at 37°C for 5-8 min, quenching with DMEM-C and counting the single cell suspension. Cells cultured in T-75 flasks were seeded at 2-4 million cells/flask in 10 mL DMEM-C. Phase-bright contrast images of cell cultures were obtained using an EVOS FL inverted microscope (Invitrogen #AMF4300).

<u>Alternative passaging strategies:</u> The Rounded/CBC subtype grows as weakly adherent/floating colonies which remain non-adherent to the tissue culture flask for the first two days post-passage making them susceptible to vacuum aspiration. To mitigate Rounded/CBC cell loss from SW480 cultures, passaging every 4-5 days is ideal as colonies become large enough to form weak adhesions with the cell culture plate, enabling slow, careful vacuum aspiration.

However, as they are still only weakly adherent, colonies can also be released from the flask by vigorous pipetting of the growth media with a p1000 pipet (no trypsin). Once released, Rounded/CBC cells can be collected into a 15 mL conical, centrifuged at 300xg for 5 min at 4°C, and the cell pellet resuspended in 0.25% trypsin-EDTA for 2-3 min at RT. Vigorous pipetting to break cell clusters before quenching with DMEM-C is effective to achieve a single cell suspension. Note: incubation with trypsin-EDTA in a 37°C water bath should not exceed 1-2 minutes as the Rounded/CBC subtype is extremely sensitive and easily lysed.

### Fluorescence Activated Cell Sorting (FACS) and Clonal derivation of SW480 subtypes

SW480 cells were washed with 1X PBS-EDTA and incubated with TrypLE Express Enzyme (Gibco) for 5 min at 37°C before quenching with DMEM-C. Cells were centrifuged at 300xg for 5 min at 4°C, washed with FACS buffer (1% BSA in PBS) and centrifuged again before resuspending the cell pellet in enough FACS buffer for 1million cells/100µL. For subtype-specific isolation, antibody incubation was performed on ice in the dark for 30 min (ROBO1-PE; NBP2- 75975PE was custom made using MAB71181 (clone 770502)). After staining, samples were diluted with FACS buffer and centrifuged at 500xg for 5 min at 4°C. Final cell pellets were resuspended in FACS buffer for 2 million cells/100µL, passed through a 35 µm filter cap, and SYTOX blue viability dye was added (1:10,000, Invitrogen) before sorting on a BD FACSAria Fusion cytometer. Sorted samples were collected into 15 mL conical tubes containing DMEM-C or 1% BSA/PBS for subculture or sorted directly into 96-well plates containing DMEM-C for single cell clonal derivations. Single cell clonal derivations were incubated at 37°C for one month before first passage. Passages 4-10 of clonally derived cells were used in the described experiments and mycoplasma tested every 3-5 months.

### Fluorescence Activated Cell Sorting (FACS) of Xenografts

Three weeks post-injection, subcutaneous tumors were processed and sorted for live cells to generate single cell libraries for mRNA sequencing. In brief, xenograft tumors excised from mice were weighed before immediate mechanical and enzymatic dissociation in DMEM-collagenase media for 1 hour in a 37°C shaker at 150 rpm (Hyclone DMEM, 2 mg/mL Collagenase type IV (Sigma), 5% FBS (Atlas Biologicals), 1% Antibiotic-Antimycotic (Gibco); for orthotopic tumors, 20 ug/mL Hyaluronidase (Sigma) was also added). After dissociation, samples were centrifuged at 500xg for 5 min at 4°C, washed with Hank’s balanced salt solution (HBSS) and centrifuged again. Pellets were resuspended in TrypLE Express Enzyme (Gibco) and incubated at 37°C for 6 min, pipetting with a p1000 pipet every 2 min to break cell clumps and adding DNase (100µg/mL = 0.3units/µL) as needed. DMEM-C quenched samples were filtered with a 40µm strainer before centrifugation at 500xg for 5 min at 4°C. Samples were washed with FACS buffer, centrifuged again, and cell pellets resuspended in FACS buffer for 2 million cells/100µL before adding SYTOX blue viability dye (1:10,000). Live cells (SYTOX blue-negative) were sorted using a BD FACSAria Fusion cytometer. FCS data files were analyzed with FlowJo v10 software (BD, FlowJo LLC).

### scRNA-seq generation and Seurat data analysis

Xenograft tumors were prepared as described above and FACS sorted live cells (SYTOX Blue-negative) were washed with 0.04% BSA in PBS twice before a final resuspension to approximately 1,000 cells/µL. Each xenograft sample was generated as an individual scRNA-seq library using the Chromium Single-Cell 3’RNA-sequencing platform with Reagent kit v3 (10X Genomics). cDNA library quantification was performed using a Qubit dsDNA HS Assay Kit (Invitrogen) and KAPA qPCR (Kapa Biosystems). The Illumina NovaSeq 6000 system was used to sequence libraries to a target read depth of 50,000 reads/cell. Fastq files were dual-aligned to both GRCh38 (human) and mm10 (mouse) genomes using Cell Ranger (v3.1.0). Human cells were identified as those containing at least 97% human transcripts per cell (i.e. transcripts beginning with ’GRCh38-’). Mouse cells were identified as those containing at least 97% mouse transcripts per cell (i.e. transcripts beginning with ’mm10-’). Further processing and normalization were performed in R studio using R (v3.6.1) and Seurat packages (v3.1.0). Cells that passed quality control filtering criteria were used for downstream analysis: low quality cells with 100 genes or less, or containing more than 20% mitochondrial genes were removed.

Additionally, a maximum threshold for genes expressed was set to 7500 and doublets were removed based on the number of unique genes. Standard Seurat processing included log normalization and the first 10-20 principal components were selected to perform clustering analysis. Specific marker genes were identified for each cluster using the ’FindAllMarkers’ function setting both the logfc.threshold and min.pct to 0.25. DEG analysis between 2 clusters was performed using the ’FindMarkers’ function to generate a list of significant DEGs for Volcano plot visualization using ggplot functions. Gene scoring of signature gene lists was calculated using the ’AddModuleScore’ function. Gene expression was visualized using DotPlot, VlnPlot and FeaturePlot functions. ClusterProfiler package (github.com/YuLab-SMU/clusterProfiler) was used to find GO term pathways associated with top DEGs with a log-fold change difference >3 enriched in each cell type. Monocle2 analysis was performed as described previously^46^.

### CRC patient data analysis

Single cell RNA sequencing datasets generated from primary CRC tumors was obtained from Lee et al. 2020^21^ and accessed from the Gene Expression Omnibus (GEO) database. The raw UMI read count data for cohorts containing 6 Belgian patients (KUL3 dataset (GSE144735)) and 23 Korean patients (SMC dataset (GSE132465)) were downloaded and used for downstream bioinformatic analysis in R studio using R (v3.6.1). Seurat integration analysis was performed to group cell types from diverse patients, with integration anchors identified across individual patient libraries as previously described^47^. Standard Seurat processing including log normalization and the first 10-15 principal components were selected to perform clustering analysis. Cell types were annotated by identifying marker genes expressed in each cluster using the Seurat function ’FindAllMarkers’, setting fold-change threshold and min.pct to 0.25. For epithelial tumor cell analysis, *EPCAM*+ cells from annotated tumor biopsies were subset and processed by the standard Seurat pipeline. Gene signature scoring was performed using the ’AddModuleScore’ function to identify tumor stem cell subtypes.

### Bulk RNA sequencing

SW480 cultures were FACS sorted for ROBO1+ (CBC) and ROBO1- (RSC) cells and immediately frozen in TRIzol (Invitrogen) and stored at -80°C until all replicates were collected. Replicates were single batch processed for RNA isolation using the Direct-zol RNA Miniprep Kit (Zymo Research) and RNA quality was determined using an Agilent Bioanalyzer DNA High sensitivity chip. 1µg of RNA was used for library generation using Illumina TruSeq Stranded mRNA Library Prep kit (Illumina #RS-122-2103). Sequencing was performed on an Illumina HiSeq 4000 (settings: RTA 2.7.7; HCS3.4.0.38) with 100 bp paired end detection. Adapter sequences were trimmed from paired-end sequencing reads and fastq files were assessed for quality using Base Calling Software (Bcl2Fastq). Data was aligned to the human genome GRCh38 using STAR (version 2.6.0c), converted to bam files and merged (samtools v1.9), and read count files were generated using Enthought Python (v7.3.2). Differential gene expression analysis was performed in R studio (v1.1.423) with R (v3.6.1) using default settings for the DESeq2 (v.1.24.0) pipeline for statistical analysis. Normalized results were log2 transformed and filtered for padj value <0.01 using DESeq2 Wald P-value test for downstream DEG analyses. PCA, volcano and gene count plots were generated in R studio using functions plotPCA and ggplot.

### Gene Ontology Enrichment Analysis

Gene ontology terms were identified by uploading top DEGs into Enrichr (maayanlab.cloud/Enrichr) and using the ClusterProfiler R package (github.com/YuLab-SMU/clusterProfiler). The ’cnetplot’ function was used to plot linked GO term pathways with enriched DEGs. Gene set enrichment analysis (GSEA) using bulk RNA-seq data of SW480 subtypes was performed using the GSEA software (v4.1.0) with Hallmark and GOBP gene lists sourced from the Molecular Signature database (MSigDB)^48,49^. Previously published gene lists and signatures used for gene scoring analyses are listed in Supplemental Data 3.

### Whole Exome Sequencing Analysis

Whole exome profiling of SW480-CBC (C2 clone) and SW480-RSC (E11 clone) samples were first prepared as libraries using the xGen™ Exome Hyb Panel v2 kit (IDT #10005153) following manufacturer instructions. In brief, 300ng of DNA was used for precapture library construction and then sheared by Covaris Acoustic shearing to generate 200bp fragmentsin length. The ends of DNA were repaired and adenylated, followed by overnight ligation of the IDT-STUB adapter onto the ends. The adapter ligated product was cleaned using AMPure XP beads and then amplified for dual indexing using 5 cycles of PCR. The resulting precapture library was cleaned with AMPure XP beads followed by quality control assays using a Bioanalyzer 2100 DNA High Sensitivity Chip and quantification by Qubit DNA High Sensitivity. Precapture libraries were pooled using 500ng of each library, and then hybridized overnight with Xgen biotinylated custom probes from IDT. Following hybridization, streptavidin beads were used to capture the exome libraries and a post capture PCR amplification of 8 cycles was performed. The final exome library was quantified by Kapa Sybr Fast universal and then sequenced on an Illumina NovaSeq 6000 platform generating 100bp paired end reads. Raw reads were transferred and quality analyzed using fastQC tool (v0.11.9). Low quality bases and adapter sequences were trimmed using Trimmomatic (v0.39). Trimmed reads were then aligned to the human reference genome (build hg19) using the Burrows-Wheeler Aligner (BWA mem, v0.7.12). Duplicated reads were removed using Picard tools MarkDuplicates (v1.130). Local realignment and base quality recalibration was done on each chromosome using the Genome Analysis Toolkit (*GATK* v4.0.4.0) using best practices. SNVs and indels were detected and filtered using GATK with HaplotypeCaller function. The output files were generated in the universal variant call format (VCF). Variant functional annotation was done using CLC Genomics workbench.

### Kaplan-Meier Survival Analysis

TCGA integration of DEGs including datasets from differentially expressed pathway genes, curated pan-TCGA data, as well as all scripts used to analyze are available at: https://github.com/cvan859/TCGA-Wnt-signaling-in-CRC. Briefly, panTCGA data was accessed from UCSC Xena on November 3, 2022 where TCGA-COAD (colon cancer) was used. Differentially expressed genes (DEGs) were accessed from TCGA somatic SNIPER results or normalized RNA-seq counts. LOF mutations were filtered for those containing missense mutations or premature stop codons were compared to individuals where no similar disruptive mutations were observed in those genes. Gene expression was subdivided in each individual receiving counts of DEGs which are above or below the mean expression for each gene. Individuals were then divided into expression categories based on the sum of low vs high counts of DEGs and used for survival analyses. Significance of survival differences between groups was calculated using a log-rank test. Survival curves were generated and analyzed using R packages survival and survminer.

### Immunohistochemistry staining

Xenograft tumors were fixed in 3.7% formaldehyde overnight at RT and washed through an increasing ethanol series (25% for 10 min then 50% for 10 min) before storage in 70% ethanol for tissue processing. Tissue embedding, FFPE processing, and H&E staining was performed by UCI Department of Pathology Experimental Tissue Resource. Unstained slides for antigen staining were de-paraffinized in xylenes and re-hydrated to PBS. Antigen retrieval was performed in boiling 10mM sodium citrate buffer pH 6.0 for 3-5 min. Tissue sections were peroxidase quenched with 3% H_2_O_2_ for 15 min at RT and blocked (10% normal goat serum/PBS) for 1 hour at RT before Avidin-Biotin blocking (Vector Laboratories, Inc). Primary antibodies were diluted in 5% serum(FBS)/4% BSA/0.3% Triton-X/PBS and incubated for 30 min at RT (CD31 @ 1:100; SMA @ 1:250). Slides were washed in 0.1% PBST (1XPBS/0.1% Tween-20) for 5 min, 3X before secondary HRP antibody incubation for 1 hour at RT diluted in 0.05% PBST. Slides were washed 3X with 0.1% PBST before incubation with Vectastain Elite ABC-HRP kit (Vector Laboratories, Inc). Finally, sections were developed with DAB Chromogen (ThermoFisher Scientific). Slides were mounted with Permount (Fisher Sci) and left to cure O/N at RT before imaging on a Keyence BZ-X710 microscope with BZ-X analyzer software (Keyence, Inc).

### Other staining protocols

Masson Trichome staining kit (ThermoFisher Scientific) was used on unstained sections following manufacturer instructions and mounted with Permount (Fisher Sci).

Alizarin Red staining was performed using a 2% Alizarin Red solution prepared in distilled water and pH adjusted to 4.1-4.3 with 10% KOH. In brief, paraffin sections were first de-paraffinized in xylenes and re-hydrated to distilled water via a decreasing ethanol series. Sections were then stained with Alizarin Red S (2% w/v) for 30-60 seconds, and excess dye drained and blotted from sections before drying. Sections were dehydrated in 100% acetone for 20 dips, then acetone:xylenes (1:1) for 20 dips. Sections were cleared in xylenes and mounted with Permount (Fisher Sci).

For BODIPY staining, harvested tumors were flash frozen in liquid nitrogen, before embedding in Tissue-Tek OCT in an isopentane-liquid nitrogen bath and stored at -80C. Frozen samples were cut to 5 µm thick sections in a cryostat and placed on SuperFrost Plus slides, and stored at -80°C until staining. Frozen sections were fixed with 4% paraformaldehyde for 15 min at RT and washed 3X with PBS before incubation with BOPIDY FL dye (1:3000; ThermoFisher Scientific) for 30 min in the dark at RT. Stained sections were washed 3X with PBS and mounted with ProLong Diamond with DAPI (Molecular Probes). BODIPY images were acquired using a Keyence BZ-X710 microscope and BZ-X analyzer software (Keyence, Inc).

### RT-qPCR

Cells were collected in TRIzol Reagent (Invitrogen) and stored at -80°C. RNA was isolated with the Direct-zol RNA Miniprep Kit (Zymo Research) and then converted to cDNA using the High-Capacity cDNA Reverse Transcription kit (Applied Biosystems). Primers were ordered from Integrated DNA Technologies and qPCR was performed using 2X qPCR master mix (ThermoFisher Scientific) and 10 µM primers (ROBO1-Forward: ATGGTTGGGGAACGTGAGAG; ROBO1-Reverse: ACTGCCAAGTTACTGGGTCTC). Ct values were normalized by delta-delta-Ct analysis using 18S Ct values as control expression (cDNA was diluted 1:1000 for 18S primer reactions). Statistical evaluation involved fold change Ct values in Prism 9 (v10) using an unpaired Student’s t-test.

### Statistics

All statistics were performed in Prism 9 (v10; GraphPad software) using an unpaired Student’s t-test to determine significance between two groups compared (CBC versus RSC). Data are presented as mean ± standard error of the mean (SEM). P-values < 0.05 were considered as statistically significant.

## SUPPLEMENTAL FIGURES

**Figure S1:**
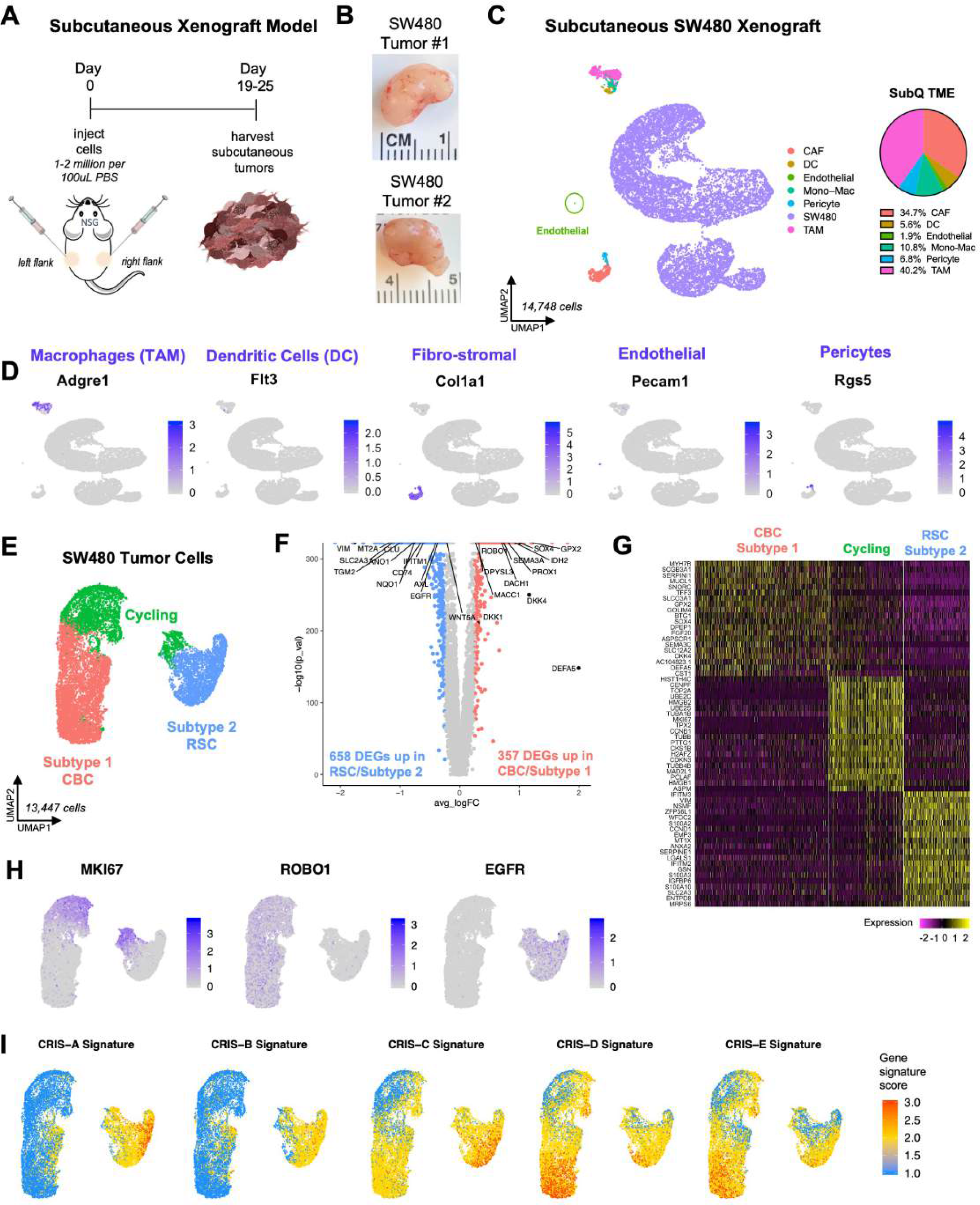
Related to Figure 1**: The human tumor compartment of SW480 subcutaneous xenografts is comprised of two subtypes.** A: Schematic diagram of the subcutaneous xenograft model and tumor development. B: Images of the two SW480 subcutaneous tumors before tissue dissociation and scRNA-seq. C: UMAP clustering of all mouse and human cells captured in SW480 subcutaneous (SubQ) xenografts with pie chart showing the average proportion of cell types in the SubQ TME. D: Gene expression plots of cell type biomarkers in the SubQ TME. E: UMAP clustering of human tumor cells reveals two distinct populations, Subtype 1 (CBC) and Subtype 2 (RSC). F: Volcano Plot highlights significant differentially expressed genes in CBC/Subtypes 1 and RSC/Subtype 2. G: Heatmap of marker genes distinguishing tumor subtype clusters. H: Gene expression plots of subtype-specific biomarkers *(ROB01* marks the CBC phenotype, *EGFR* marks the RSC subtype, MKl67 marks the cycling cells in both subtypes). I: Gene signature plots of CRC patient­ derived CRIS molecular subtype signatures.

**Figure S2:**
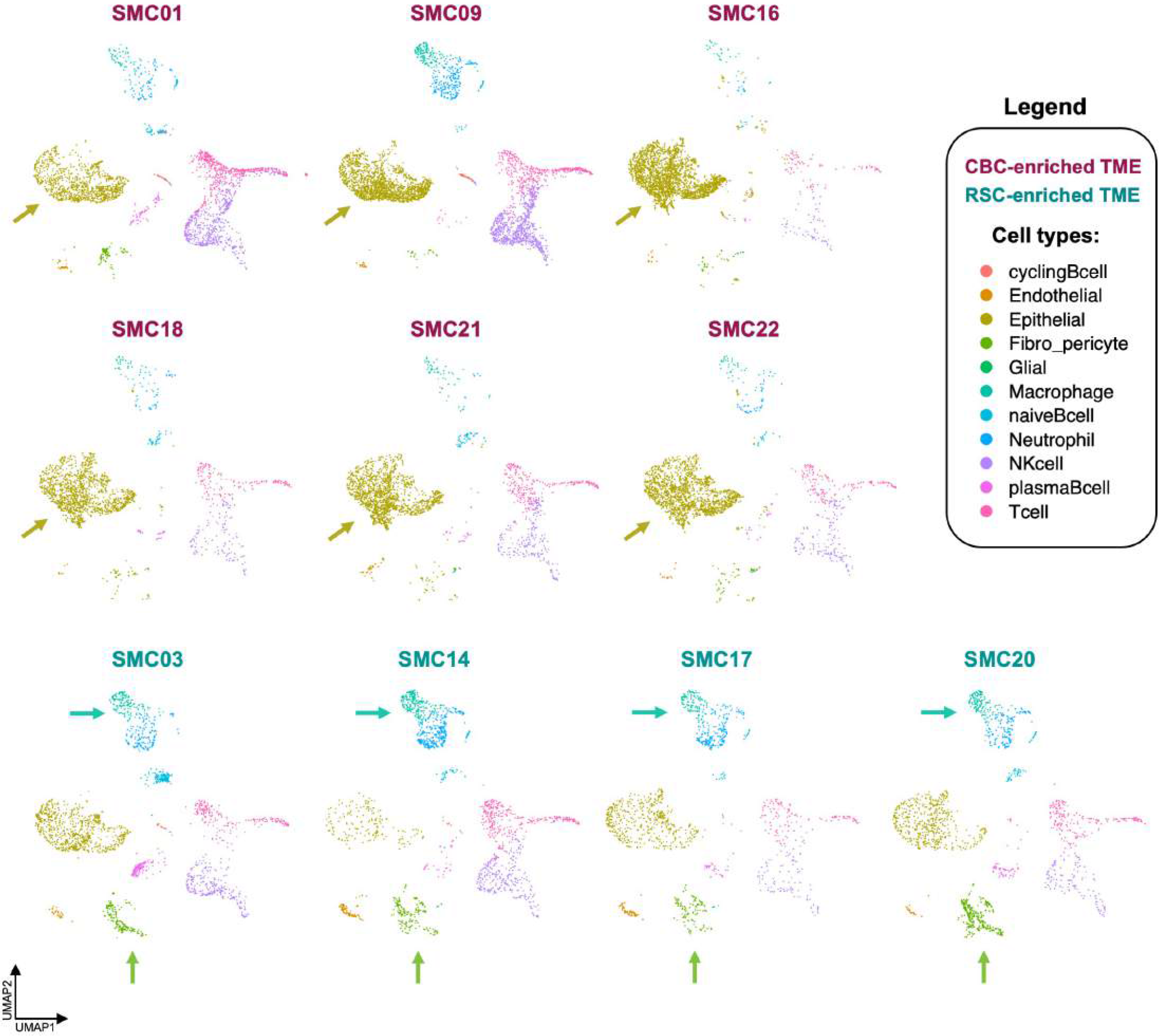
Related to Figure 2**: CRC patients exhibit different TME compositions associated with CBC and RSC phenotypes.** UMAP clustering of scRNA-seq data from Korean (SMC) patient tumor biopsies. Patients in maroon have tumors with strong CBC signatures and contain a larger proportion of epithelial cells (gold arrow). Patients in teal have strong RSC signatures and contain a larger proportion of stromal fibroblast-pericytes (green arrow) and myeloid immune cell types (macrophages and neutrophils, blue arrow).

**Figure S3:**
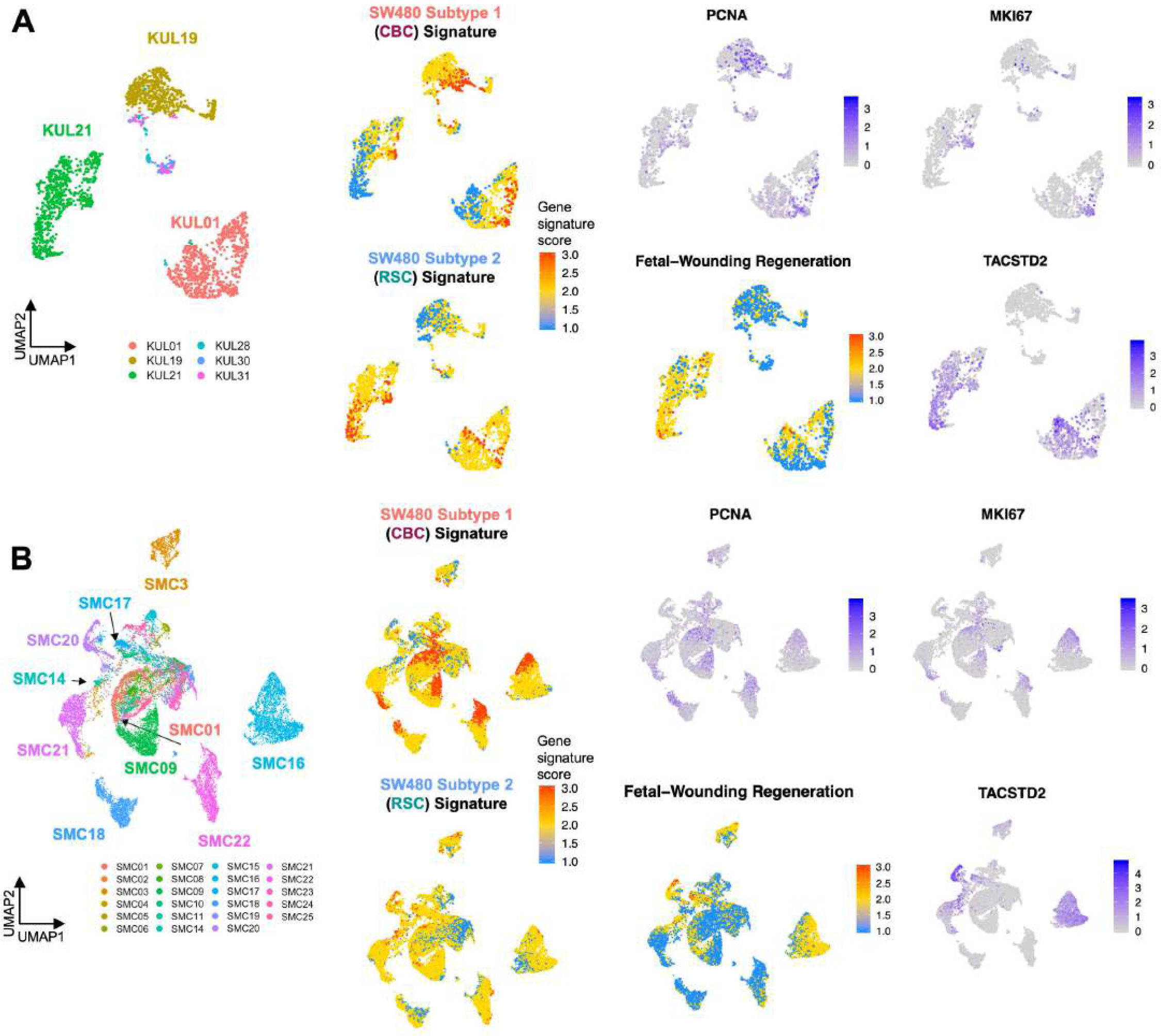
Related to Figure 2**: CRC stem cell subtype signatures co-exist in most patients in varying proportions.** UMAPs of Belgian/KUL {A) and Korean/SMC (B) patients published in Lee et al., 2020, with gene expression and signature plots distinguishing stem cell subtypes.

**Figure S4:**
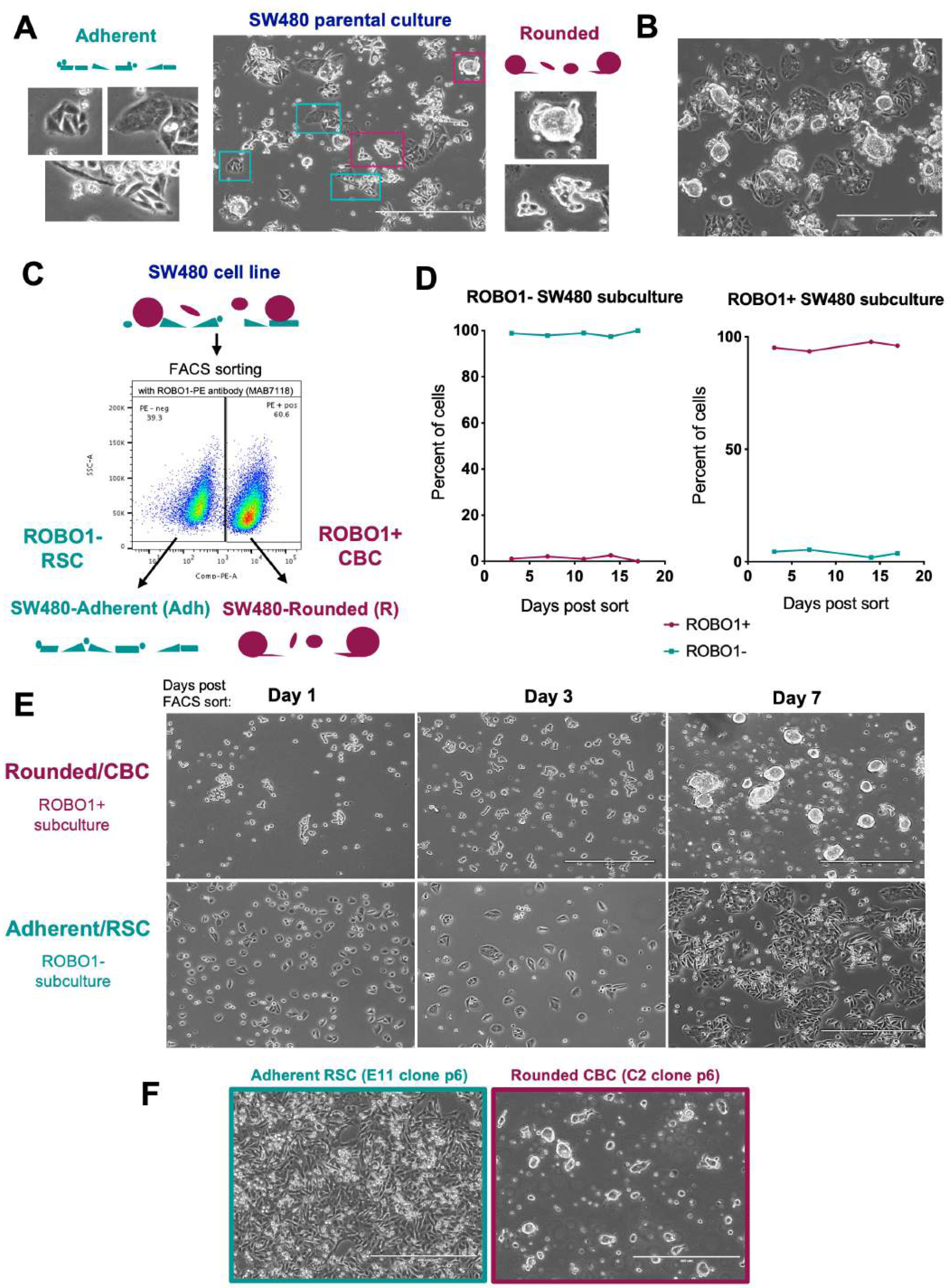
Related to Figure 3**: Use of ROBO1 expression as a novel sorting strategy to isolate SW480 stem cell subtypes.** A-B: Phase contrast images of the SW480 culture with magnified insets showing a mixture of cellular morphologies, scale= 400µm. C: Fluorescence Activated Cell Sorting (FAGS) using ROBO1-PE antibodies separates two populations of morphologies (Adherent versus Rounded). D: Flow Cytometry Analysis of subcultured ROBO1+ and ROBO1- sorted cells retain positive and negative cell surface expression of ROBO1, respectively. E: Phase contrast images of ROBO1+ Rounded/CBC and ROBO1-Adherent/RSC subcultures days after FAGS sorting from the parental SW480 culture, scale= 400µm. F: Phase contrast images of single cell clonal derivations of Adherent/RSC (E11 clone) and Rounded/CBC (C2) cultures; scale= 400µm.

**Figure S5:**
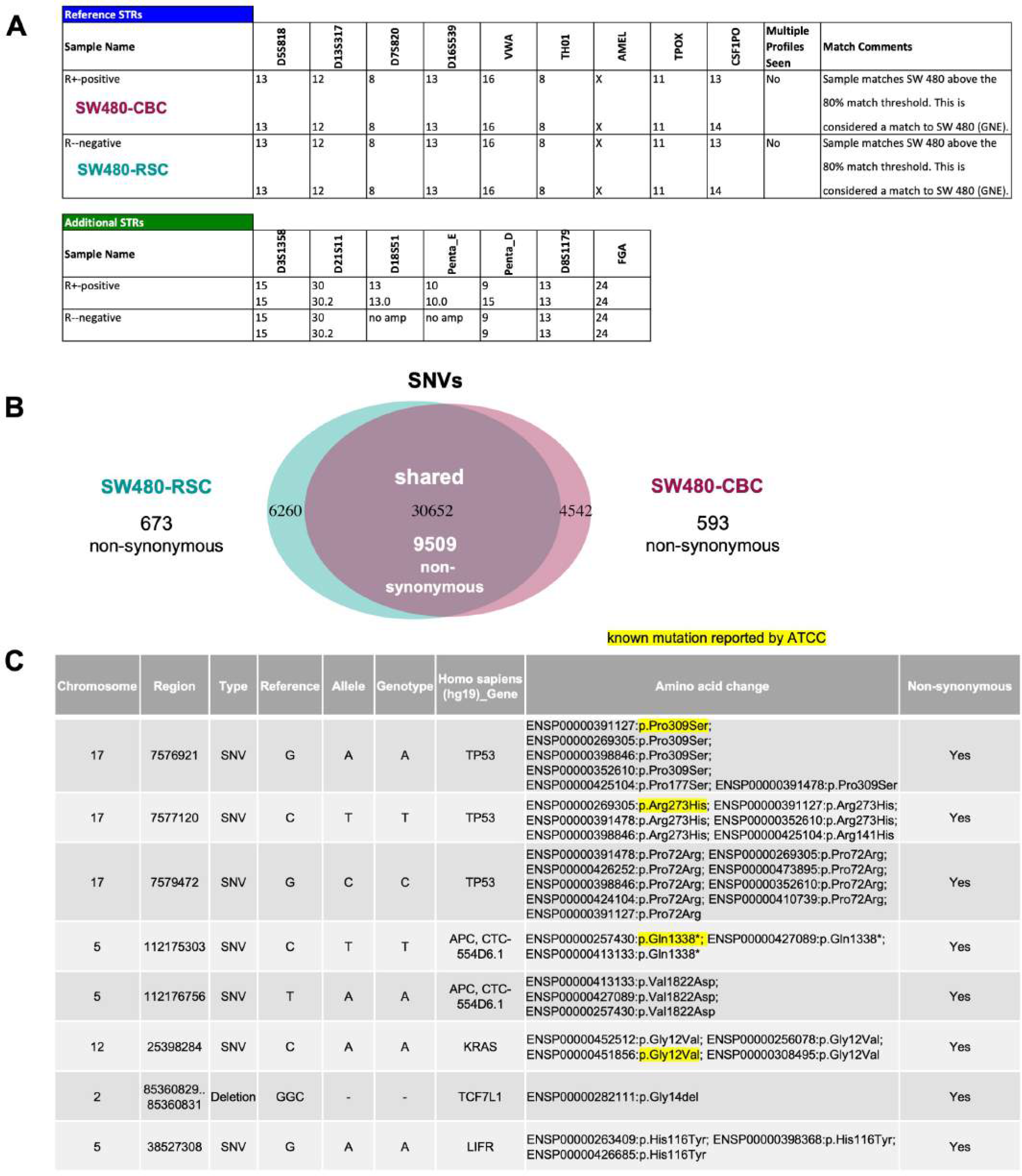
related to Figure S4: STR profiling and exomic sequencing confirm a common SW480 origin with identical cancer driver mutations in CBC and RSC subtypes. A: Cell line authentication report of short tandem repeat (STR) profiling in ROSO1+ sorted (R+-positive) SW480- CBC and ROBO1- sorted (R-negative) SW480-RSC cells. B: Venn diagram summary of the total number of single nucleotide variations (SNV) and of those that encode non-synonymous protein mutations, generated from exomic sequencing of SW480-RSC and SW480-CBC subtypes. C: Table of shared SNV mutations in both SW480 subtypes, both RSC and CBC cells contain the same cancer driver mutations in *TP53, APC,* and *KRAS* gene loci.

**Figure S6:**
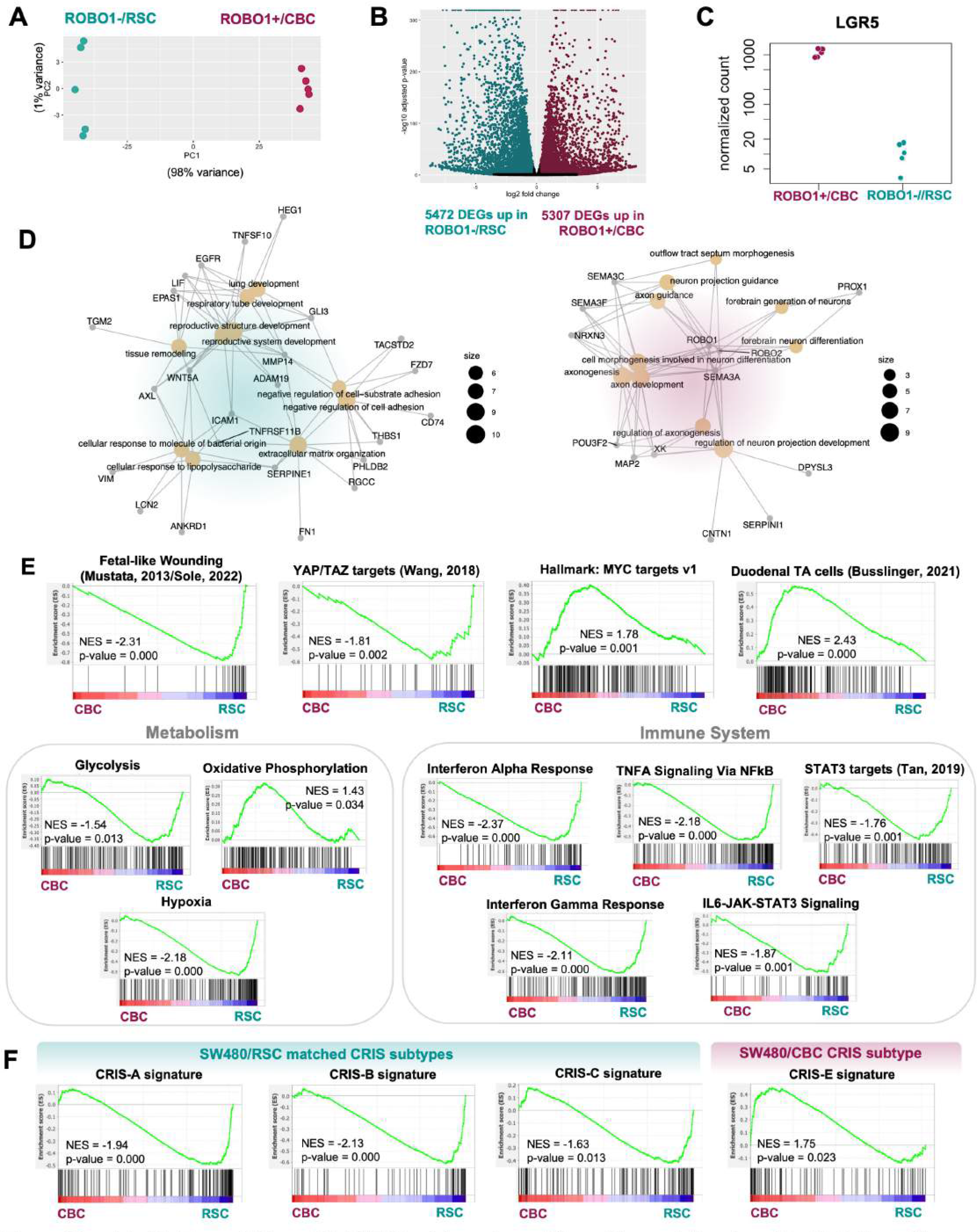
Related to Figure 3**/Figure S4: SW480 subtypes have different transcriptional profiles that align with distinct binary classifications and human colorectal cancer CRIS subtypes.** A: Principal component analysis plot comparing transcriptomes of ROBO1+/CBC and ROBO1-/RSC sorted cells by bulk RNA sequencing, n = 5. B: Volcano Plot highlights thousands of significant DEGs in RSC/ROBO1- sorted cells and CBC/ROBO1+ sorted cells. C: LGR5 mRNA expression in each sorted replicate, n=5. D: Top marker genes (logFC>3) expressed in RSC/ROBO1- cells (left) and CBC/ROBO1+ cells (right) are linked to different GO biological processes; top DEGs in RSC cells were linked to biological processes such as extracellular matrix production, cell adhesion and cytoskeleton regulation while CBC DEGs align with neuronal processes such as axon guidance and neuronal differentiation. E: Gene Set Enrichment Analysis (GSEA) using MSigDB hallmarks, GO biological processes, and published gene signatures distinguish CBC from RSC cells. F: Gene Set Enrichment Analysis (GSEA) of the CRC patient-derived gene signatures show subtype-specific alignment to distinct CRIS molecular subtypes (see Supplemental Data 3 for gene lists).

**Figure S7:**
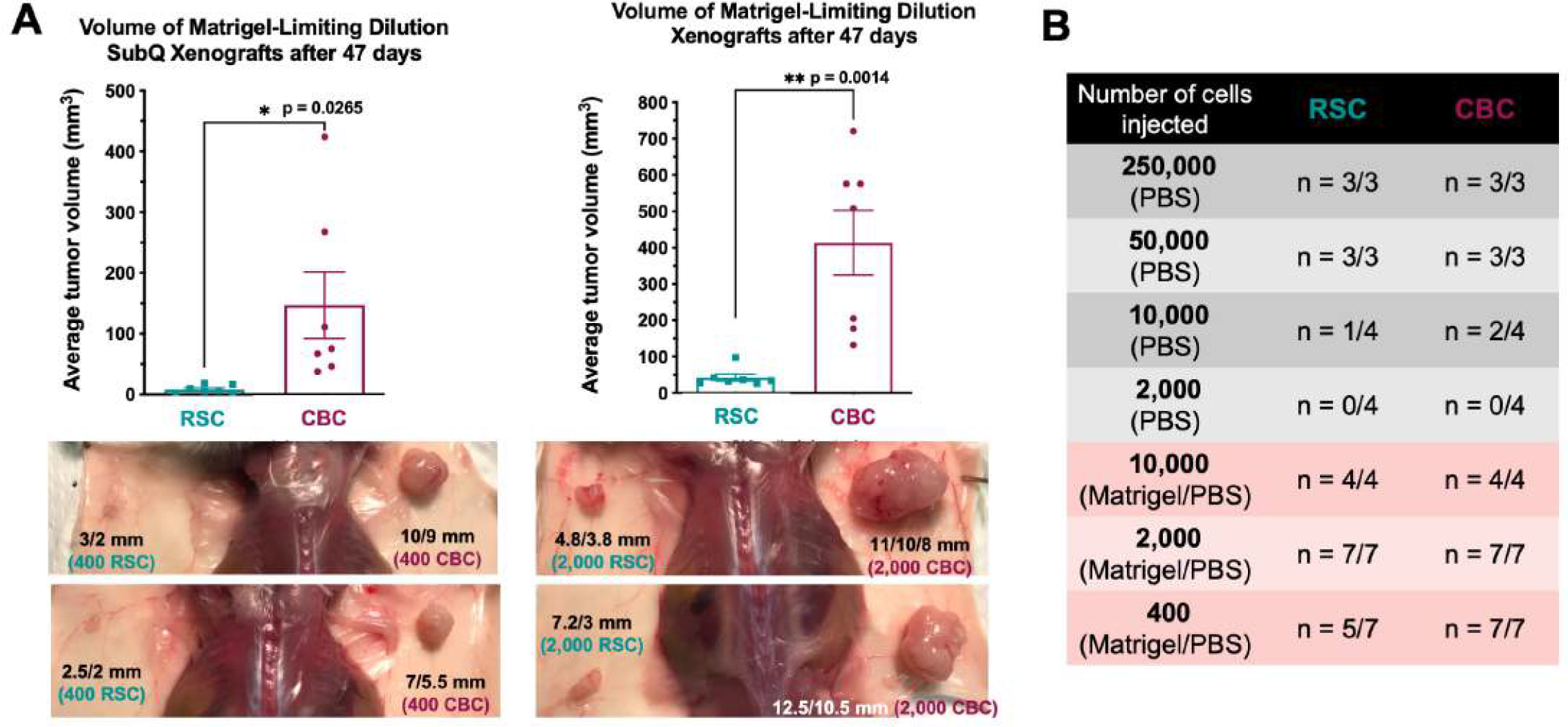
Related to Figure 3**: SW480 subtypes have similar tumor initiating potential but CBC cells form larger tumors.** A: Representative images and weights of limiting dilution subcutaneous (SubQ) xenograft experiments injecting 400 cells (A) or 2,000 cells (8) per inoculant. B: Summary of limiting dilution experiments injecting smaller numbers of cells down to a minimum number in PBS or Matrigel/PBS.

**Figure S8:**
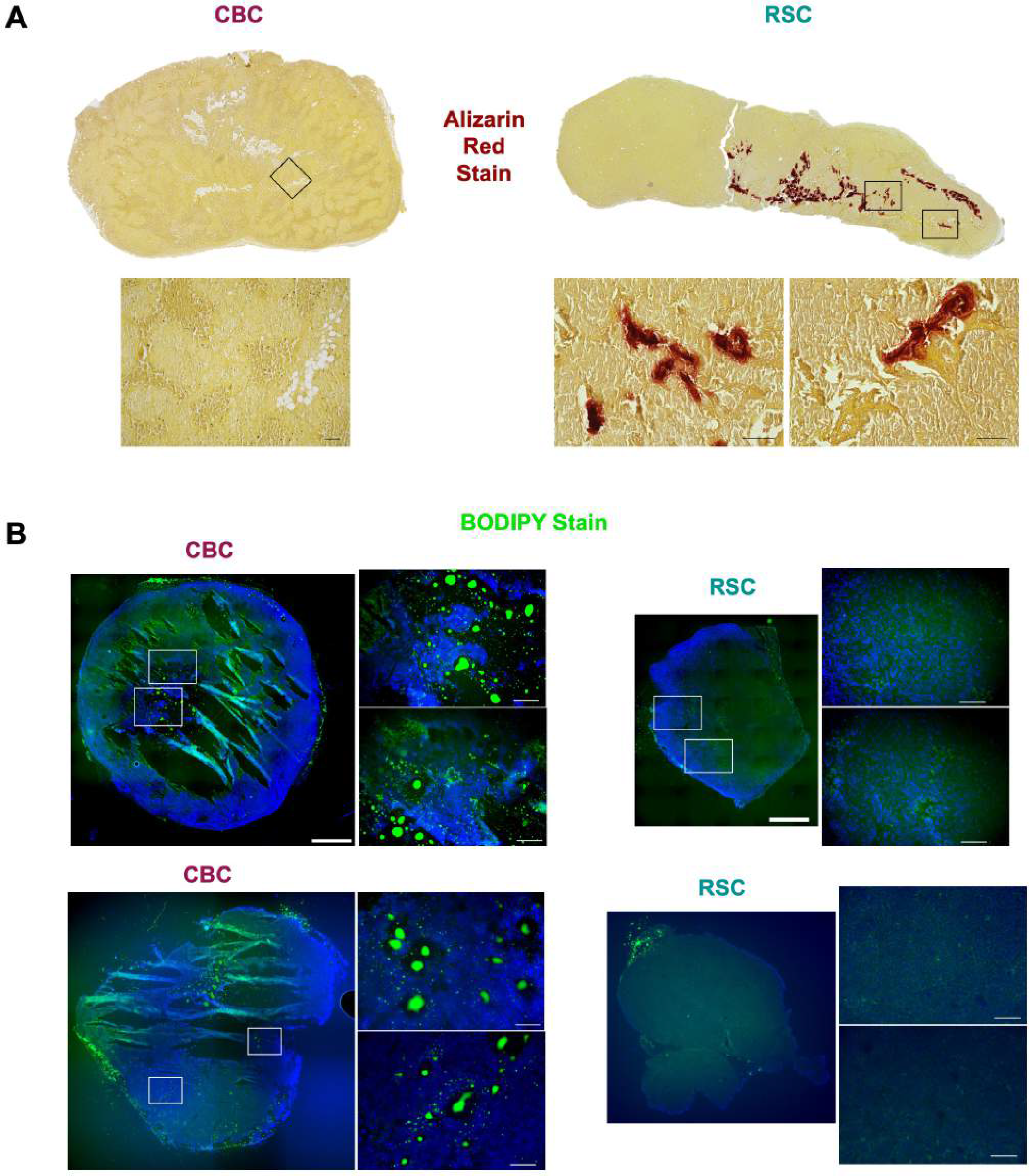
Related to Figure 3**: Diverse stromal features of osteogenesis in RSC tumors and adipogenesis in CBC tumors.** A: Alizarin Red staining marks areas of calcium-deposited tissue (a surrogate for bone-associated extracellular matrix) found only in SW480-RSC tumors, with magnified images of tumor core below (CBC inset = 1OX and RSC insets= 20X), scale= 100µm. B: BODIPY staining illuminates lipid droplets (a biomarker of adipocytes) inside the tumor core of CBC tumors, with 20X magnified images on right, scale= 1mm and 100µm (20X).

**Figure S9:**
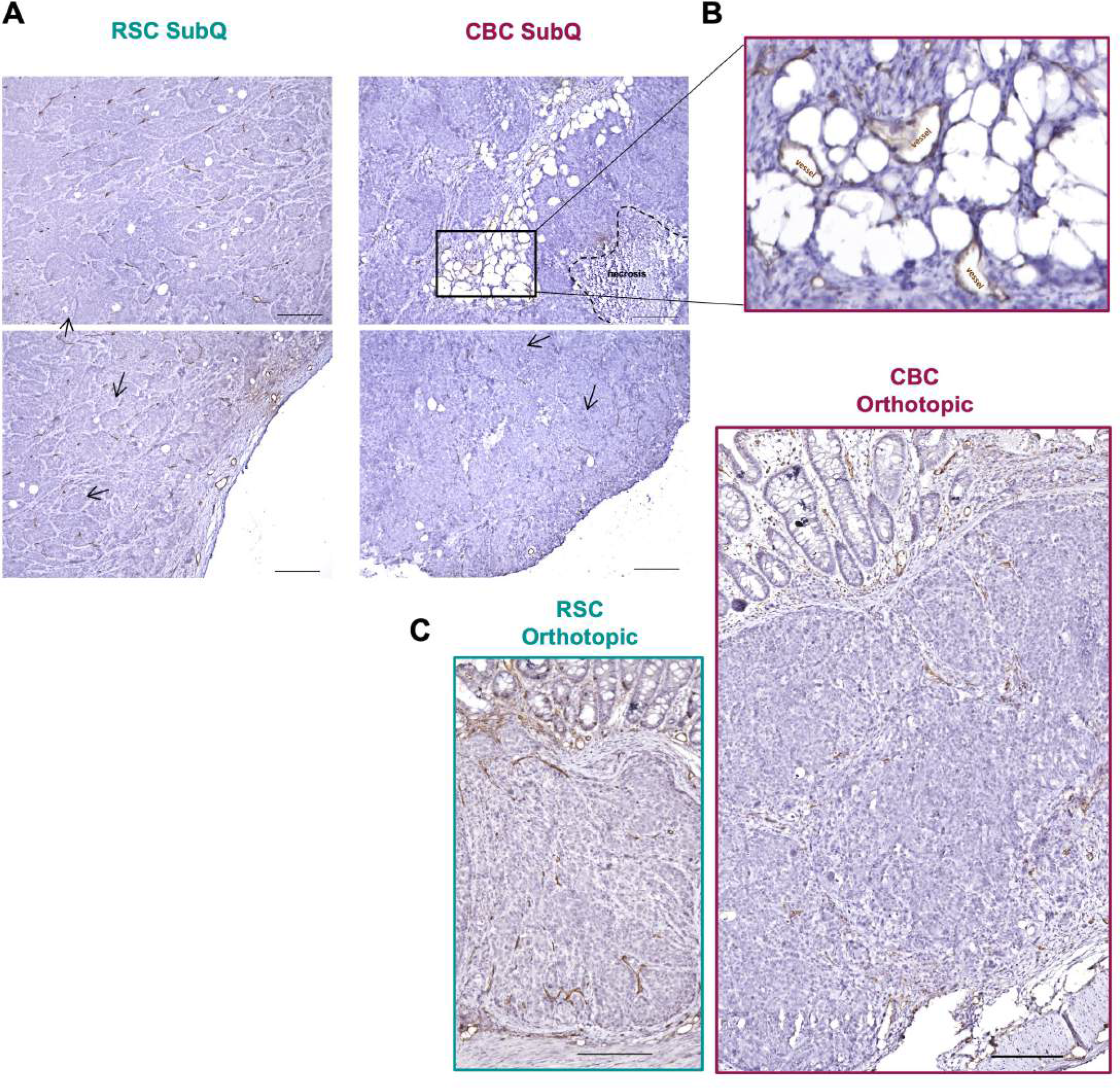
Related to Figures 3 **and 4: RSC tumors demonstrate increased vascularization in both xenograft models.** A: lmmunohistochemistry staining for Cd31+ vasculature in subcutaneous (SubQ) tumors, arrows point to examples of vessels, scale= 200µm. B: Magnified image of the CBC tumor core identifies few Cd31 vessels within an adipogenic core. C: lmmunohistochemistry staining for Cd31+ vasculature in orthotopic tumors marks thicker vessels infiltrating RSC tumors, scale = 200µm.

**Figure S10:**
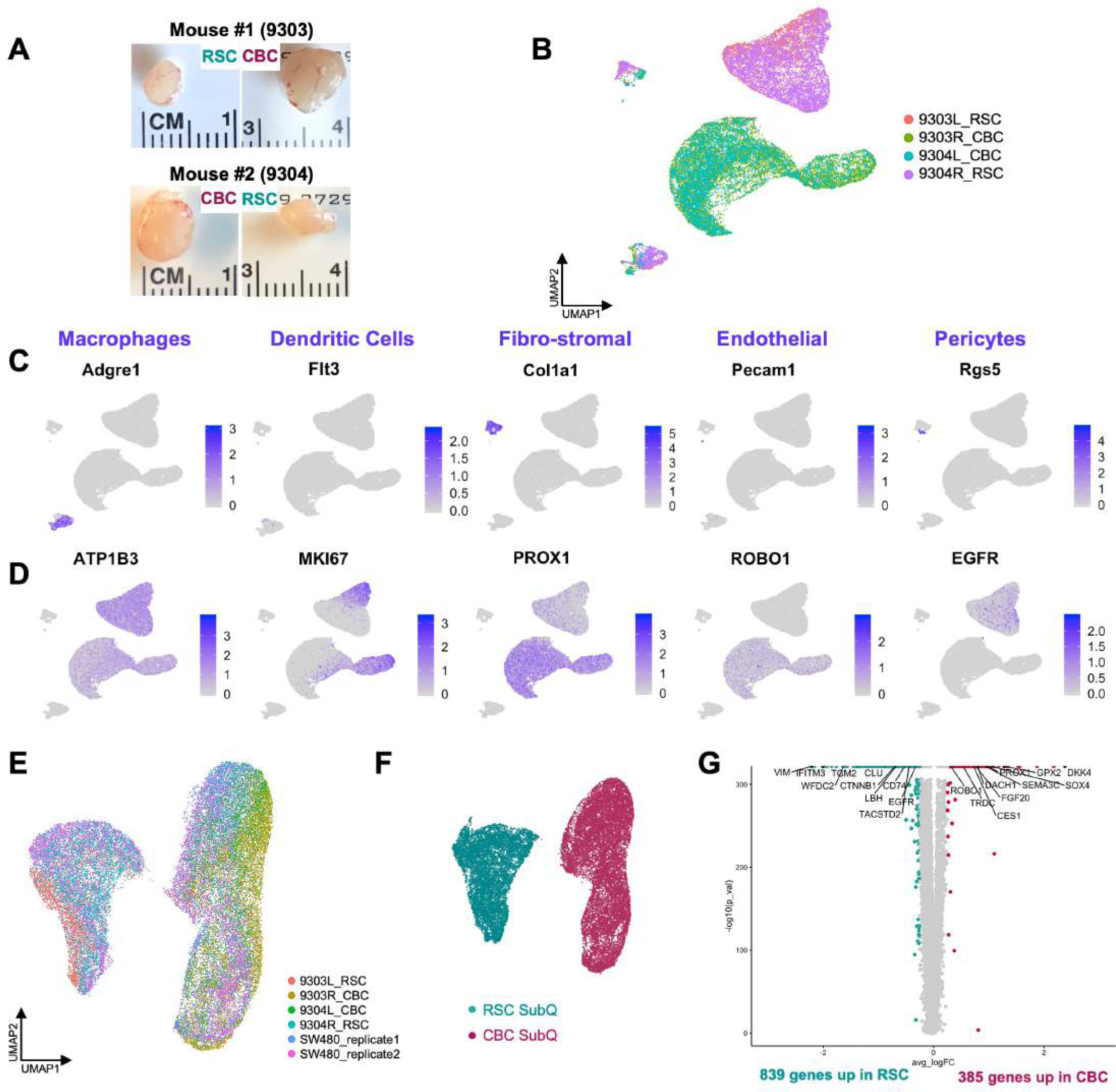
Related to Figure 3**: Stromal, immune and human tumor populations are transcriptionally distinct in RSC and and CBC subcutaneous tumors.** A: Images of subcutaneous (SubQ) xenograft tumors prior to tissue dissociation for scRNA-seq. B: UMAP clustering of all mouse and human cells from CBC and RSC SubQ tumors, n = 2 each subtype. C: Gene expression plots of marker genes for various cell types found in the SubQ TME. D: Gene expression plots of human biomarkers (human specific antigen CD298 *(ATP183),* cell cycle marker *(MK/67+)* and subtype-specific biomarkers *(PROX1, ROB01, EGFR)).* E: UMAP clustering of human tumor cells from SW480 subtype-specific and mixed xenografts (from Figure S1), colored by tumor ID. The clustering of CBC and RSC cells in SubQ tumors propagated as separate xenografts (i.e. as right/left flank tumors in the same mouse, e.g. mice 9303 and 9304) were nearly identical to their transcriptomic signatures when mixtures of both cell subtypes were propagated in a single xenograft (e.g. SW480_replicates), suggesting tumor signatures are strongly intrinsic. F: UMAP of human tumor cells, colored by subtype-specific xenograft. G: Volcano Plot highlights significant differentially expressed genes (DEGs) in RSC versus CBC cells in SubQ xenografts (positive logFC values represent genes upregulated in CBC cells, negative logFC values represent genes upregulated in RSC cells; for top 100 DEG lists for each subtype see Supplemental Data 3).

**Figure S11:**
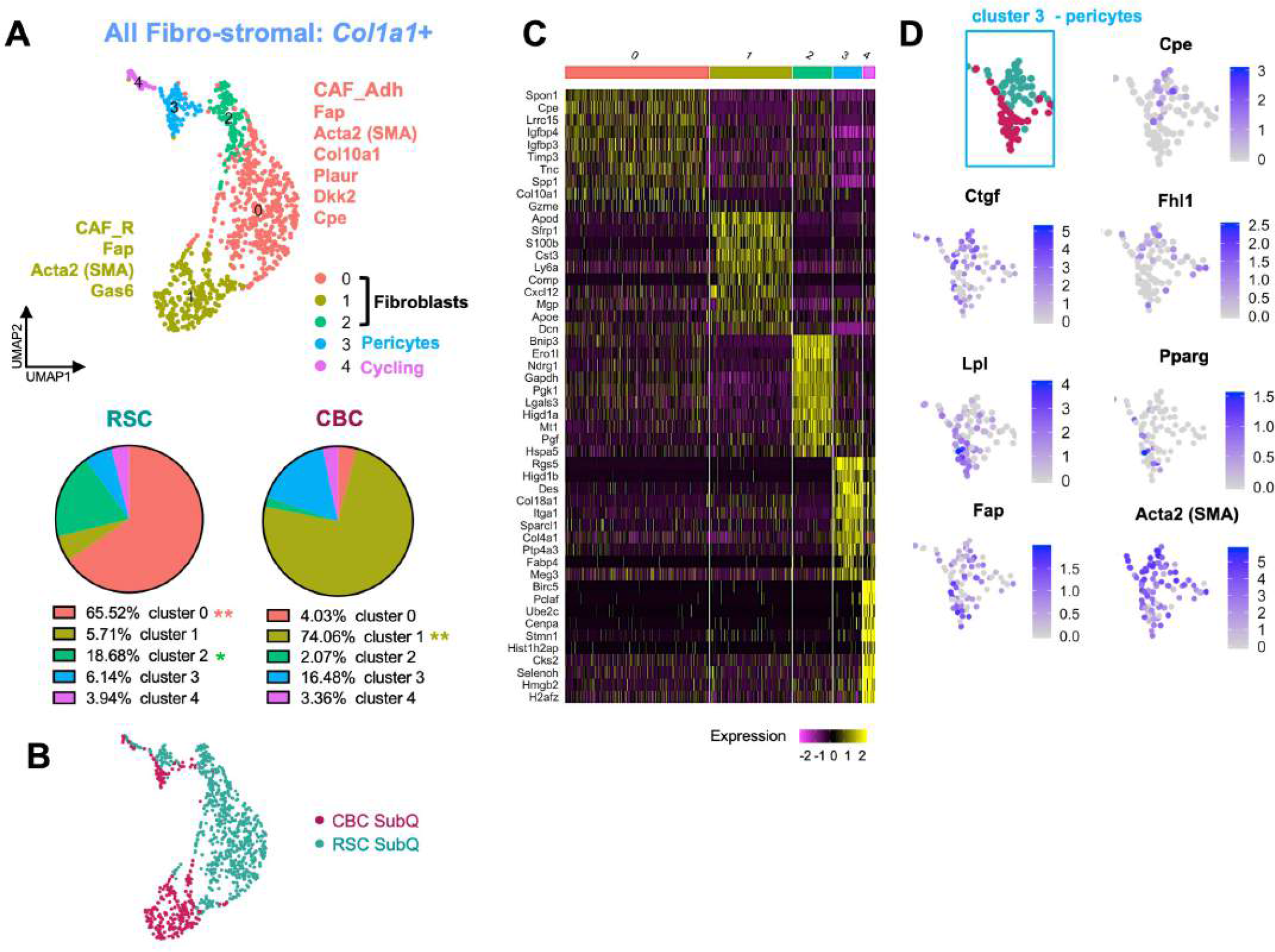
Related to Figure 3**: Fibroblasts and perlcytes have distinct transcriptional profiles** in **CBC and RSC subcutaneous tumors.** A: UMAP clustering of *Co/1a1+* expressing stromal cells in subcutaneous (SubQ} tumors, with pie charts showing significant differences in proportional composition between tumor subtypes. B: UMAP of fibro-stromal cells colored by subtype-specific xenograft. C: Heatmap of marker genes expressed in the different fibro-stromal clusters. D: UMAP crop of the pericyte cluster colored by tumor subtype xenograft and with corresponding gene expression plots of fibroblast *(Fap, Acta2),* adipocyte *(Lpl, Pparg),* and osteoblast *(Ctgf, Cpe, Fh/1)* related marker genes. P values=*< 0.05, ** < 0.01 (Student’s t test).

**Figure S12:**
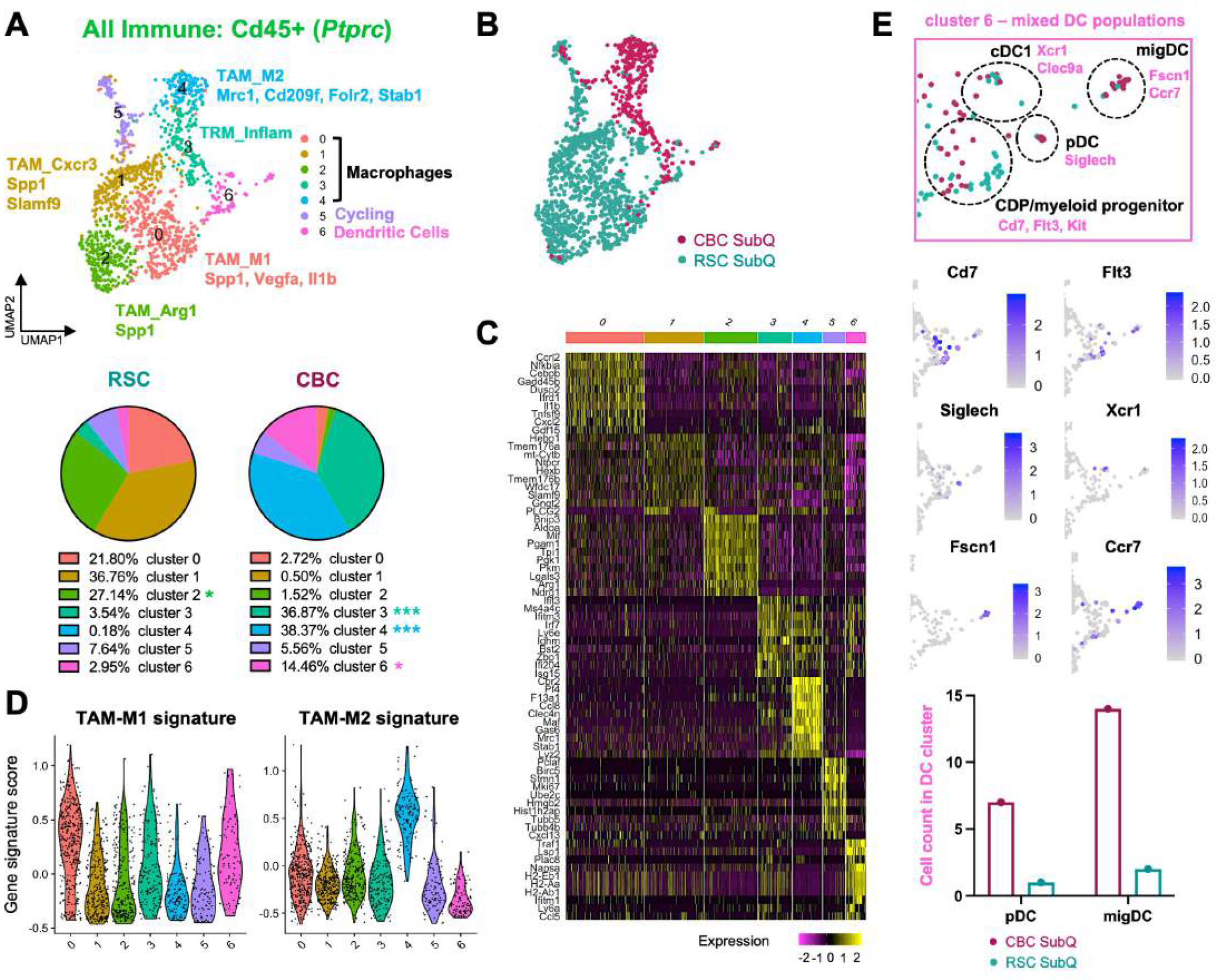
Related to Figure 3**: Tumor associated macrophages and dendritic cell populations have distinct profiles in CBC and RSC subcutaneous tumors.** A: UMAP clustering of *Cd45+* immune cells in subcutaneous (SubQ) tumors, with pie charts showing significant proportional differences in clusters between tumor subtypes; Tumor associated macrophage (TAM) states are distinct clusters from Tissue-resident macrophages (TRM). B: UMAP of immune cells colored by subtype-specific xenograft. C: Heatmap of marker genes for immune clusters. D: Violin plot of gene signature scoring for M1 and M2 associated TAM marker genes amongst immune clusters. E: UMAP crop of the dendritic cell (DC) cluster, colored by subtype­ specific xenograft, with gene expression plots of biomarkers of DC sub-populations: common DC progenitor (CDP), plasmacytoid DC (pDC), conventional DC 1 (cDC1), and migratory DCs (migDC). Bar plot highlights enriched populations of pDCs and migDCs in CBC xenografts. P values = * < 0.05, *** < 0.001 (Student’s t test).

**Figure S13:**
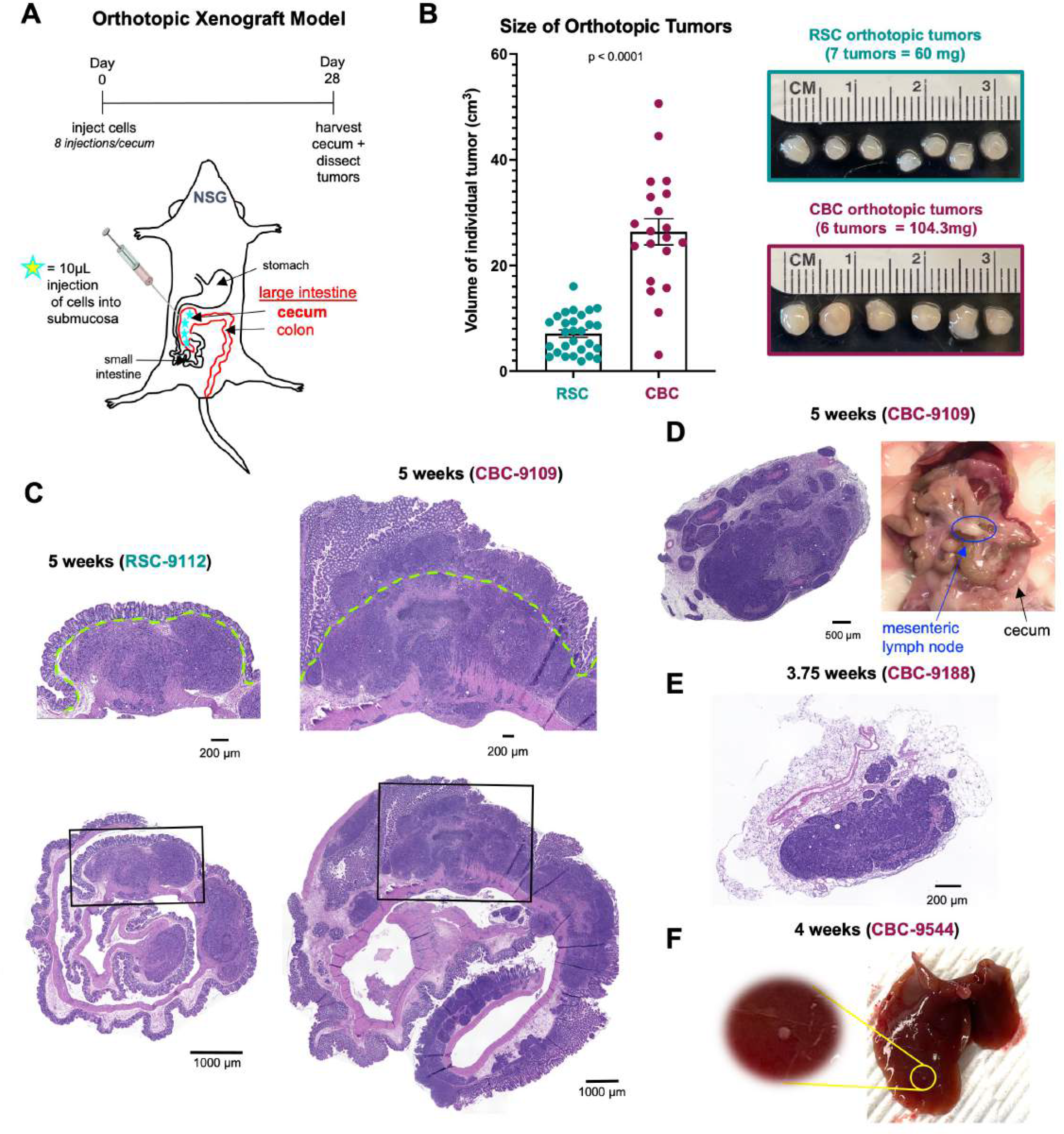
Related to Figure 4**: The CBC subtype displays a more invasive phenotype in orthotopic xenografts.** A: Schematic of the orthotopic tumor model diagramming multi-injections into the cecum of NSG mice and duration of experiment. B: Average size of individual orthotopic tumors with representative images of tumors harvested from one mouse cecum. C: Representative H&E snapshots of RSC and CBC orthotopic tumors after 5 weeks *in vivo,* magnified individual tumors (top) and whole-cecum “swiss rolls” (below), scale= 200µm and 1000µm, green line marks the muscularis mucosae. D: H&E staining and image of mesenteric lymph node filled with tumor cells found in the CBC orthotopically injected mouse from (C), scale bar= 500µm. E: H&E staining of a mesenteric lymph node filled with tumor cells, found in another CBC-injected mouse, scale= 200µm. F: Image of liver dissected from a CBC orthotopically injected mouse (in the scRNA-seq cohort) with likely indication of metastatic tumor growth.

**Figure S14:**
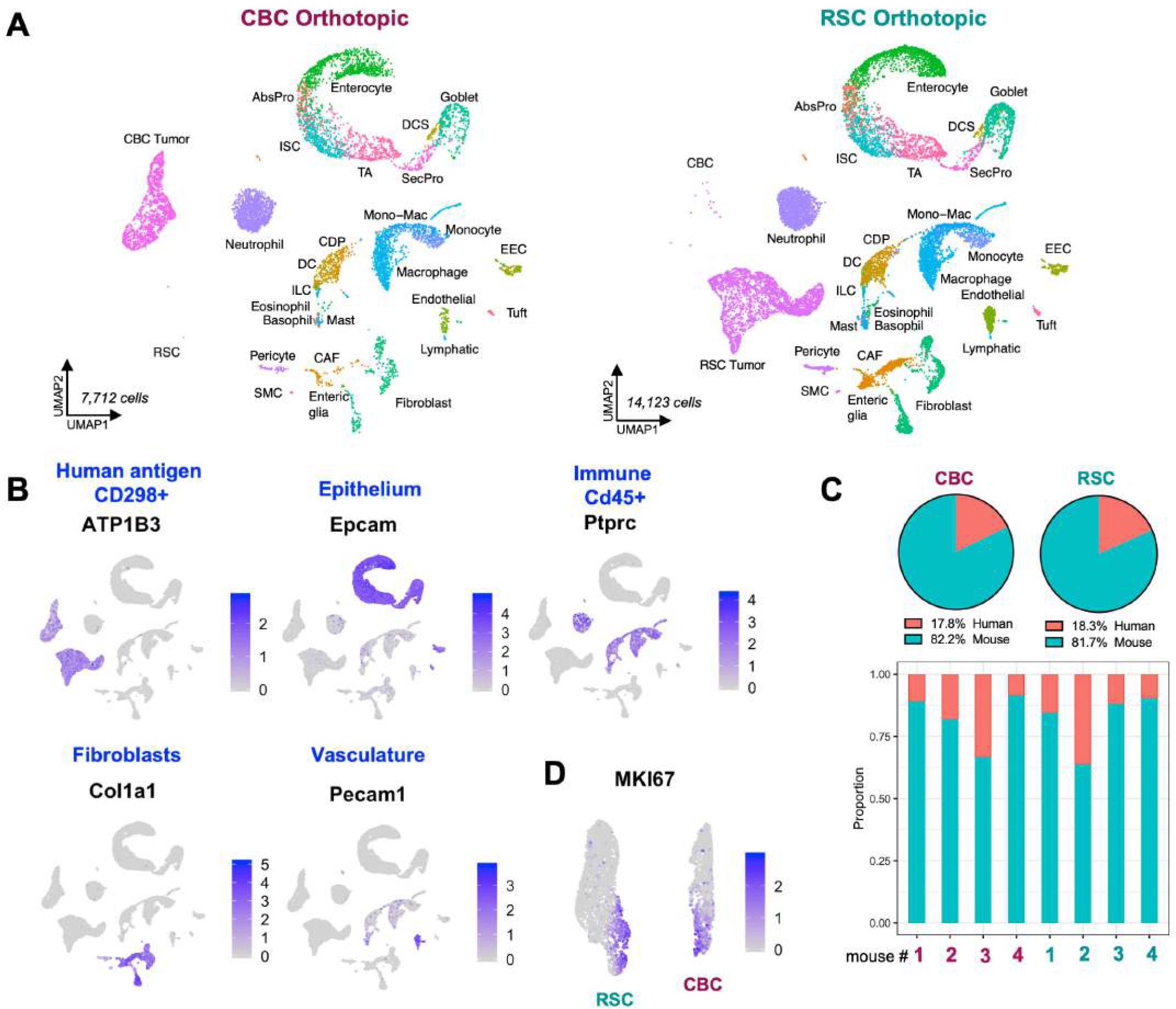
Related to Figure 4**: Single cell sequencing of orthotopic xenografts reveals a diverse and expanded tumor microenvironment that dominates the tumor setting.** A: UMAP clustering of all mouse and human cells captured in CBC and RSC orthotopic tumors, n = 4 each subtype. B: Gene expression plots of marker genes for the major cellular compartments in orthotopic tumors. SW480 human tumor cells are identified by human specific antigen CD298 (A*TP183).* C: Proportion of human tumor to mouse TME on average (pie chart) and in each replicate (below). D: Gene expression plot of a cell cycling biomarker, *MK/67,* in the human tumor compartment.

**Figure S15:**
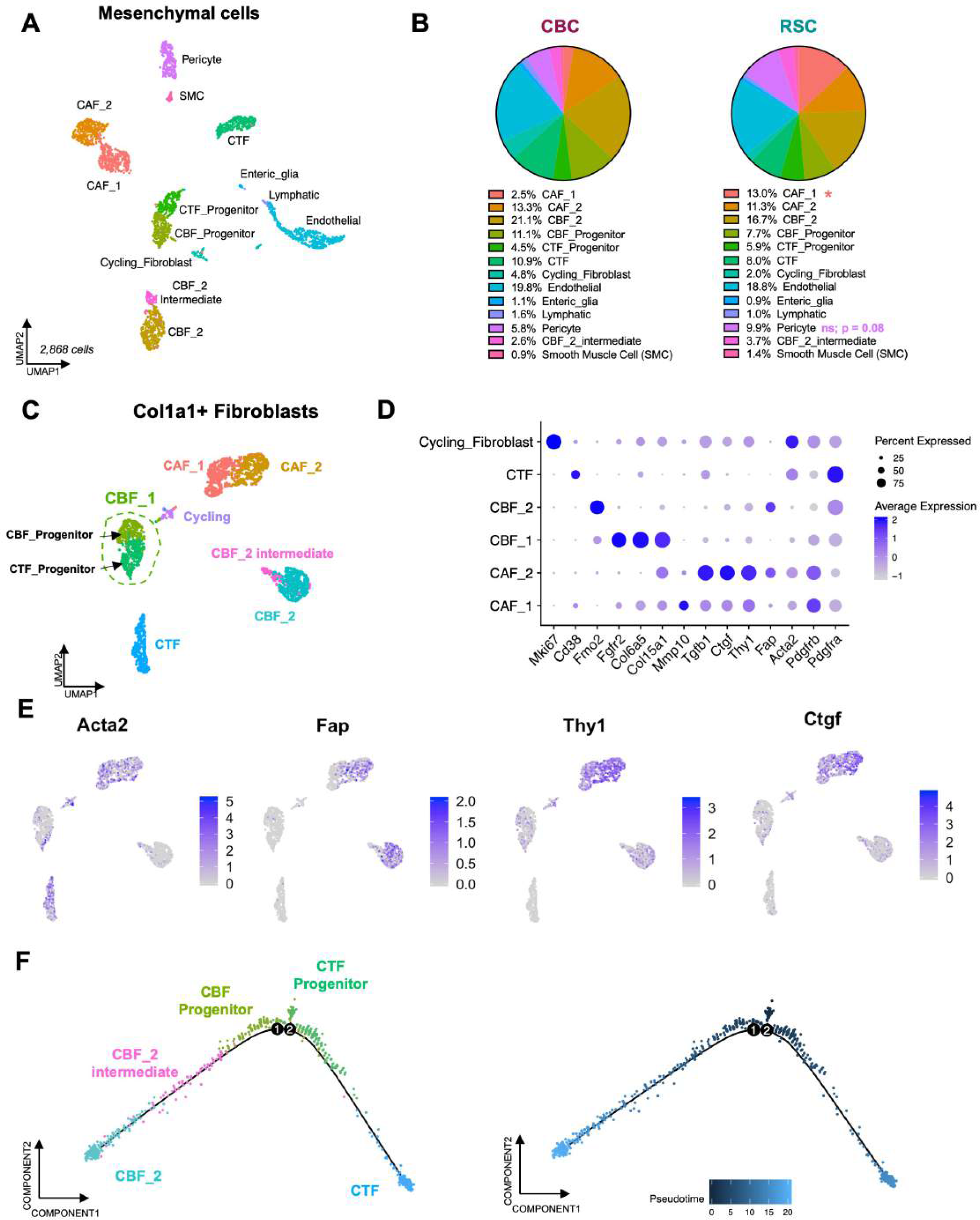
Related to Figure 5**: Identification of normal and cancer-associated fibroblast populations.** A: UMAP clustering of all mesenchymal cells in orthotopic tumors. B: Pie charts showing proportional compositions of mesenchymal cell types in CBC and RSC orthotopic tumors; P value = * < 0.05 (Student’s t test). C: UMAP clustering of *Co/1a1+* mesenchymal cells subsets all normal and cancer-associated fibroblast (CAF) populations. D: DotPlot of intestinal fibroblast markers: differential *Pdgfra* expression distinguishes crypt top fibroblast (CTF) from crypt bottom fibroblast (CBF) populations (Brugger et al., 2020; ref. 29). The universal fibroblast progenitor marker *Co/15a1* (Buechler et al., 2021; *Nature* 593(7860):575-579) identifies the CBF_1 cluster as a bi-potent progenitor to CTF and CBF_2 clusters. E: Gene expression plots of CAF and myofibroblast marker genes (Acta2/SMA, *Fap, Thy1, Ctgf).* Since *Fap* is co-expressed by CAFs and normal CBF_2 fibroblasts, *Thy1* and *Ctgf* are better, exclusive gene markers of CAFs. F: Monocle2 analysis orders normal crypt fibroblast populations by “pseudotimefl with the two CBF_1 sub-clusters distinctly divided: the darker green cluster appears to be closely connected to the CTF population and is placed at the beginning of ’pseudotime’, therefore we refer to this initiating Monocle population as “CTF_Progenitor”; the lime green cluster is situated on a path that leads to the CBF_2 populations and is therefore referred to as “CBF_Progenitor”.

**Figure S16:**
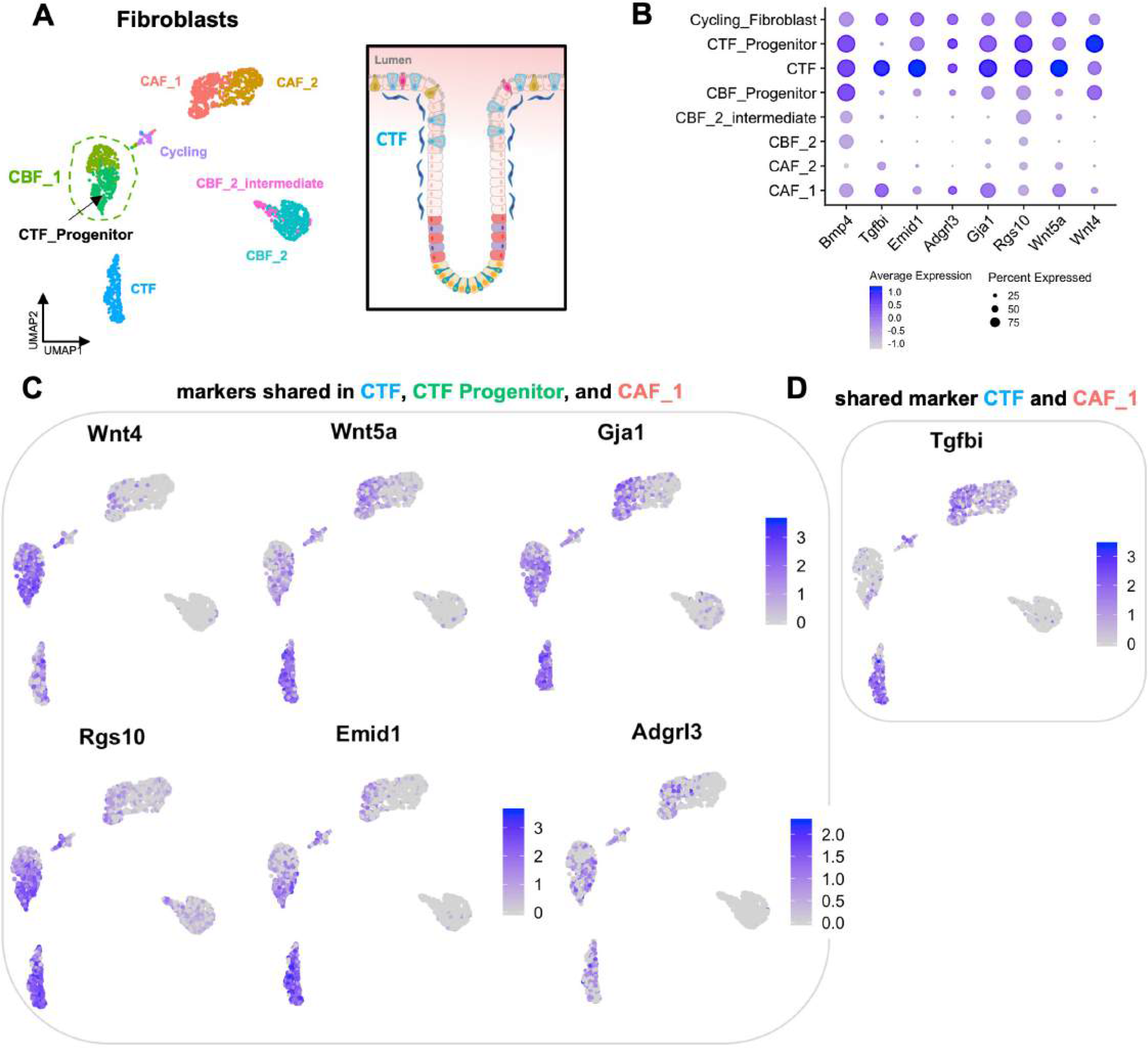
Related to Figure 5**: Shared gene expression relationships between Crypt Top Fibroblasts and CAF**_**1 cells.** A: UMAP clustering of fibroblast states, highlighting the two transcriptionally distinct sub-populations within the CBF_1 cluster, indicating here the CTF_Progenitor state. B: DotPlot of genes co-expressed between CTF, CTF_Progenitor and CAF_1 populations. C: Gene expression plots of biomarkers co-expressed in CTF, CTF_Progenitor and CAF_1 populations *(Wnt4, Wnt5a, Gja1, Rgs10, Emid1, Adgrl3).* D: Gene expression plot of shared *Tgfbi* expression in CTF and CAF_1 populations.

**Figure S17:**
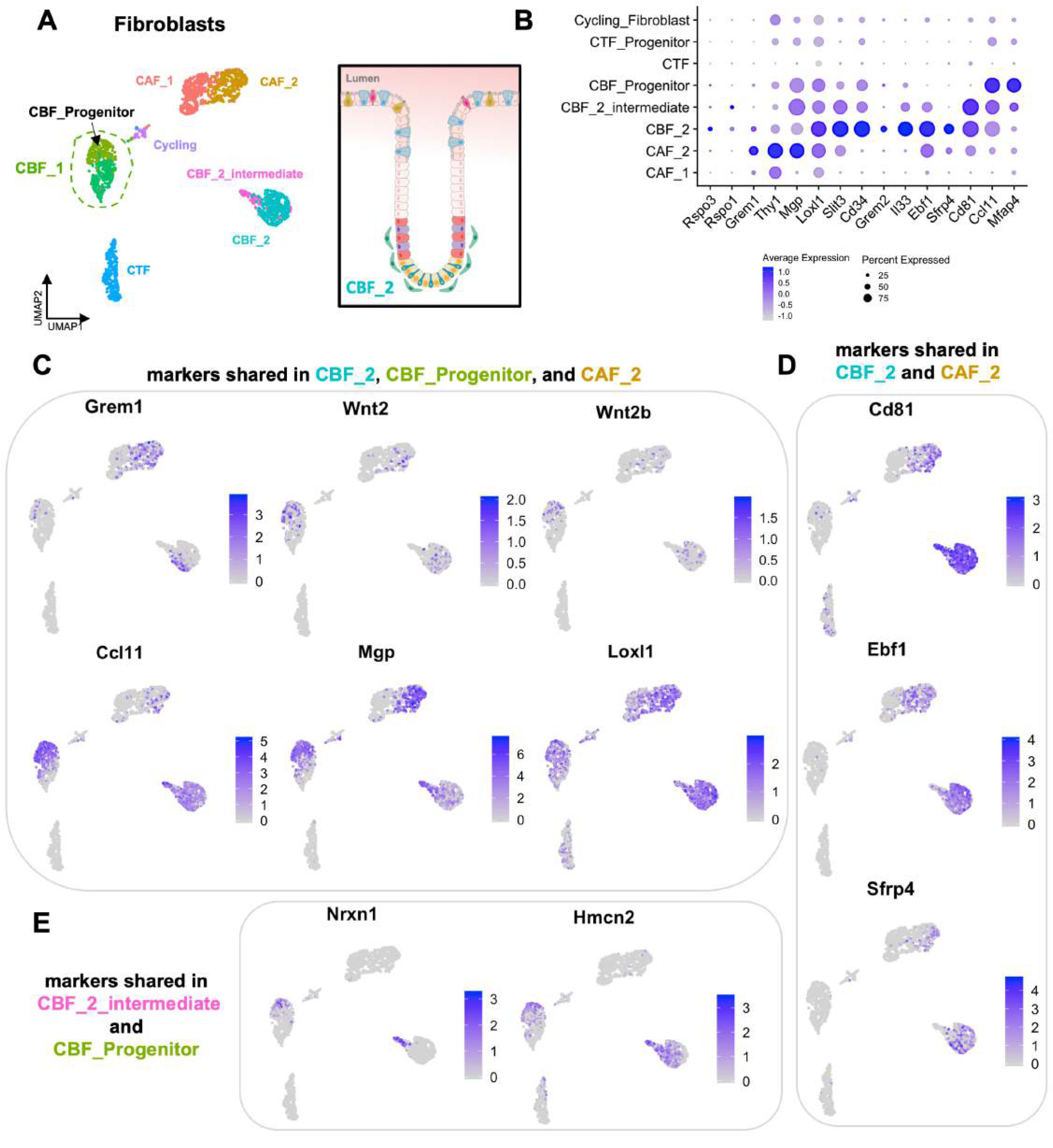
Related to Figure 5**: Shared gene expression relationships between Crypt Bottom Fibroblasts and CAF_2 cells.** A: UMAP clustering of fibroblast states, highlighting the two transcriptionally distinct sub-populations within the CBF_1 cluster, indicating here the CBF_Progenitor state. B: DotPlot of genes co-expressed in CBF_2 sub-populations and CAF_2 cells. C: Gene expression plots of genes co-expressed in CBF_Progenitor, CBF_2, and CAF_2 populations include genes for secreted factors *(Grem1, Wnt2, Wnt2b, Cc/11, Mgp).* D: Gene expression plots of genes co-expressed in CBF_2 and CAF_2 populations *(Fap, Cd81, Ebf1, Sfrp4).* E: Gene expression plots of genes co-expressed in CBF_2_intermdiate and CBF_Progenitor populations *(Nrxn1, Hmcn2, Gucy1a1, Ces1d).* CBF_2 is considered to be the telocyte population due to expression of biomarkers: *Cd34, Grem1, Grem2, Wnt2, Wnt2b, Rspo1* and *Rspo3*.

**Figure S18:**
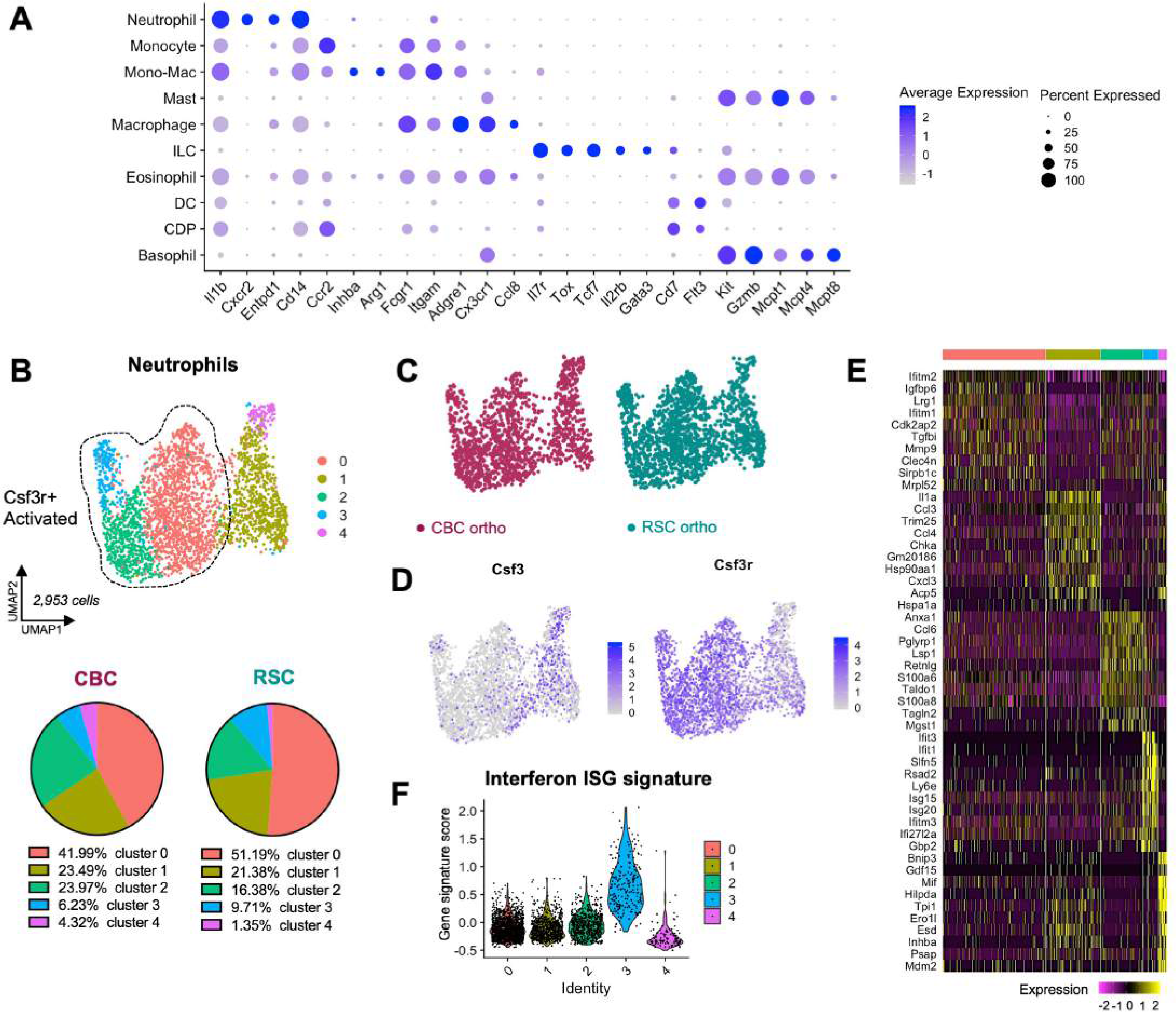
Related to Figure 6**: Neutrophils are the most abundant immune cell type in the orthotopic setting but little difference is observed between tumor subtypes.** A: DotPlot of marker genes for the immune cell types found in the orthotopic tumor microenvironment. B: UMAP clustering of neutrophils, with pie charts showing siimilar proportional compositions of neutrophil subclusters between tumor subtypes. C: UMAP clustering of neutrophils colored by subtype-specific xenograft. D: Gene expression plots of resting *(Csf3+)* and activated *(Csf3r+)* neutrophil states (Park et al., 2022; *Front* Onco/.12:932608). E: Heatmap of marker genes distinguishing neutrophil clusters. F: Violin plot of the Interferon Stimulated Gene (ISG) signature scoring amongst neutrophil clusters, the *Csf3r+* population (cluster 3) has enriched expression.

**Figure S19:**
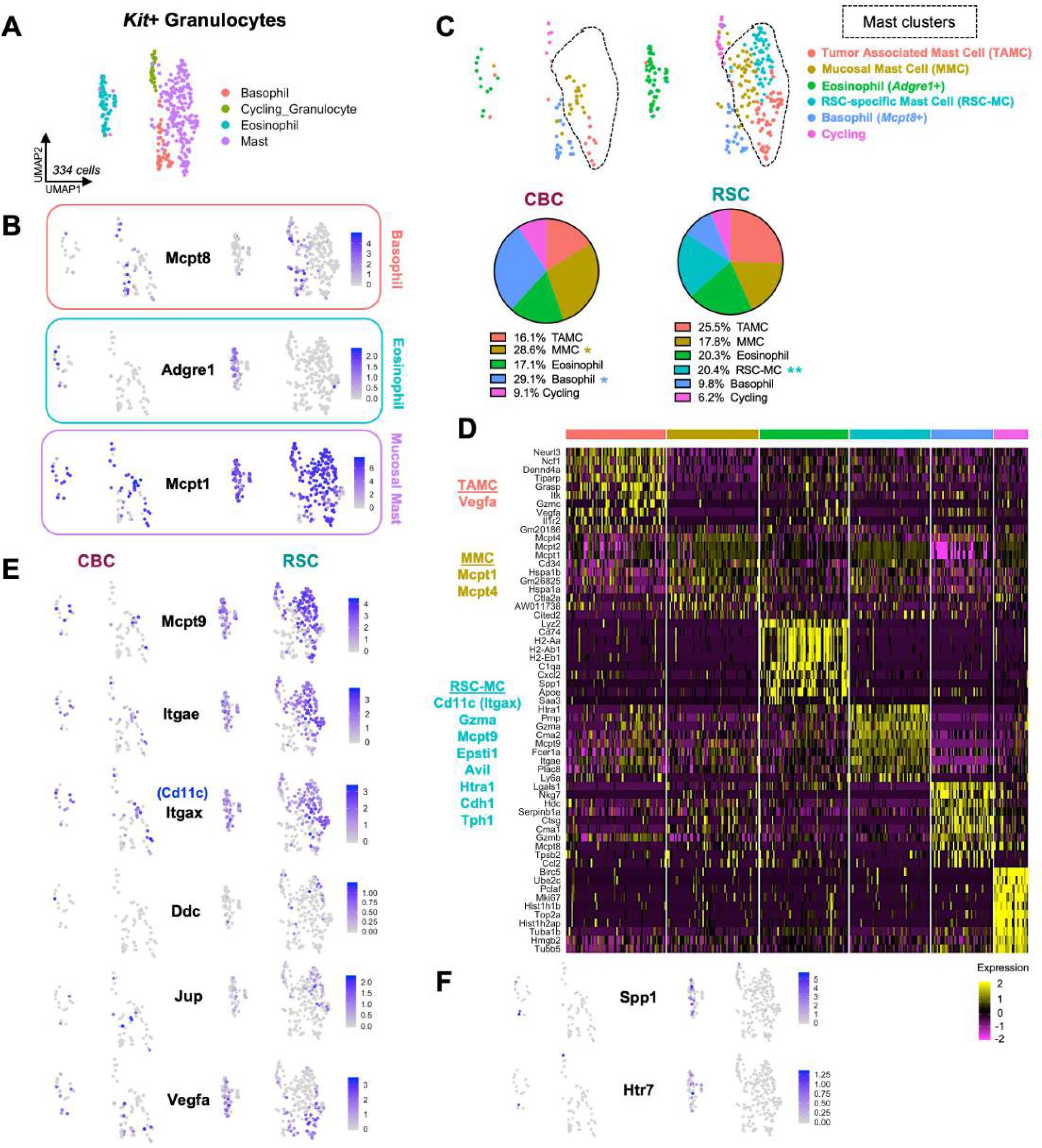
Related to Figure 6**: Mast cells exhibit multiple phenotypes in the orthotopic tumor microenvironment with a novel population found exclusively in RSC tumors.** A: UMAP clustering of *Kit+* granulocytes (mast, eosinophil, basophil) present in orthotopic tumors. B: Gene expression plots of marker genes identifying basophil, eosinophil and intestinal mucosa! mast cell populations in each subtype-specific xenograft. C: UMAP of *Kit+* granulocytes colored by transcriptional state, with pie charts highlighting proportional differences of populations in tumor subtypes. Mast cells comprise three distinct transcriptional populations, one of which is found exclusively in RSC tumors (RSC-MC). D: Heatmap of marker genes distinguishing mast cell states with key marker genes of the novel RSC-MC population is listed in teal. E: Gene expression plots of upregulated genes enriched in RSC mast cells. F: Gene expression plots of genes upregulated in RSC eosinophils. P values = * < 0.05, ** < 0.01 (Student’s t test).

**Figure S20:**
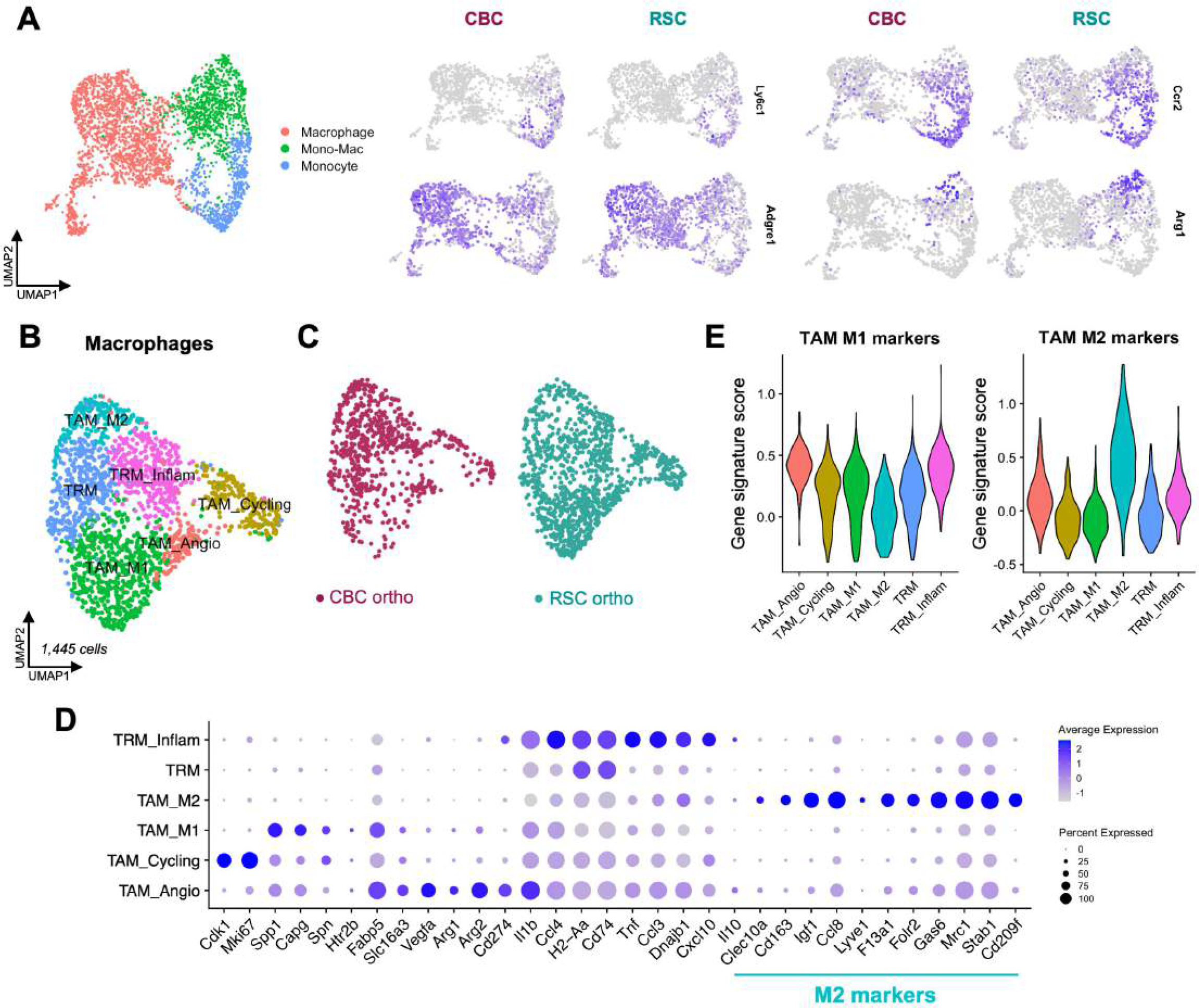
Related to Figure 6**: Monocytic immune cells are a large, expanded compartment** in **the orthotopic microenvironment including several tumor associated macrophage states.** A: UMAP clustering and gene expression plots showing *Ly6c1+* monocytes, *Adgre1+* macrophages, and an intermediate *Ccr2+* monocyte-derived macrophage (Mono-Mac) population, with a subset expressing *Arg1.* B: UMAP clustering of tumor associated macrophage (TAM) and tumor resident macrophage (TRM) states. C: UMAP of macrophages states colored by subtype-specific xenograft. D: DotPlot of marker genes distinguishing each TAM and TRM population. Tissue Resident Macrophages (TRMs), identified by a lack of polarized M1/M2 markers and heightened expression of MHC-11 markers *H2-Aa* and *Cd74,* were a distinct population found in both tumor subtypes with a subset expressing multiple inflammatory cytokines (TRM_lnflam; *111b, Cc/3, Cc/4, Tnf, Cxc/10, Dnajb1)*. E: Violin plot of gene signature scoring of M1 and M2 associated TAM marker genes (see Supplemental Data 3 for gene lists).

**Figure S21:**
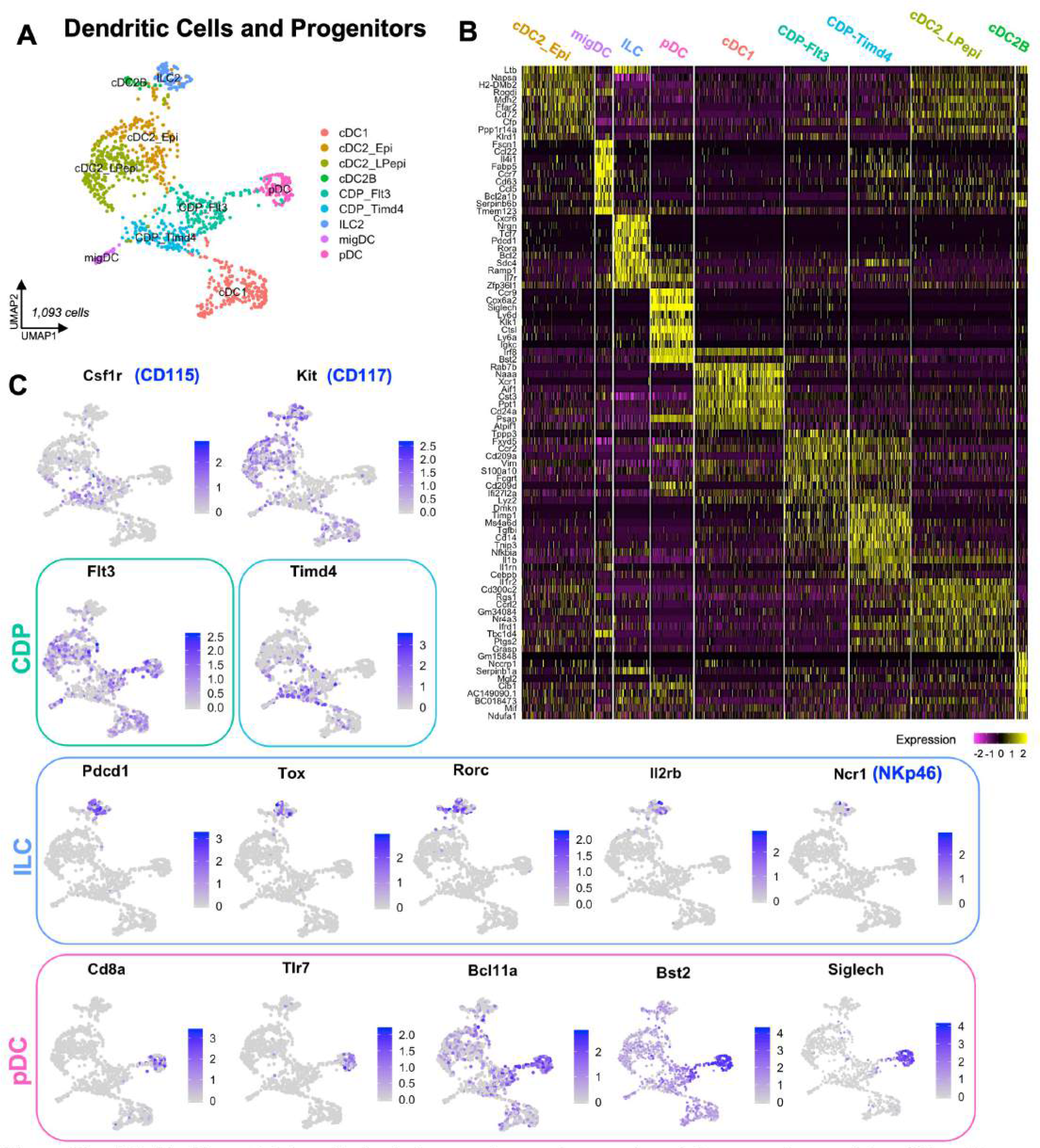
Related to Figure 6**: The orthotopic tumor microenvironment contains many types of dendritic cells with distinct transcriptional profiles.** A: UMAP clustering of dendritic cell (DC) subtypes: conventional DC 1 and 2 (cDC1, cDC2) with additional cDC2 heterogeneity, migratory DC (migDC), plasmacytoid DC (pDC), common DC progenitor (CDP), and innate lymphoid group 2 cells (ILC2). B: Heatmap of marker genes identifying each dendritic cell and progenitor subpopulation. C: Gene expression plots of biomarkers of dendritic cell (DC) subpopulations: CDP cells marked by CD115+;CD11710 express different levels of *Flt3* and *Timd4.* Innate lymphoid cells (ILCs) are an early precursor to the adaptive immune compartment and express genes involved with T-cell *(Pdcd1, Tox, Rorc)* and NK cell *(l/2rb,* Ncr1/NKp46) functions. Plasmacytoid DCs (pDC) have unique markers *(Cd8a, Siglech* and *Tlr7)* and more strongly express *Bst2* and *Bcl11a* compared with other DC subpopulations.

**Figure S22:**
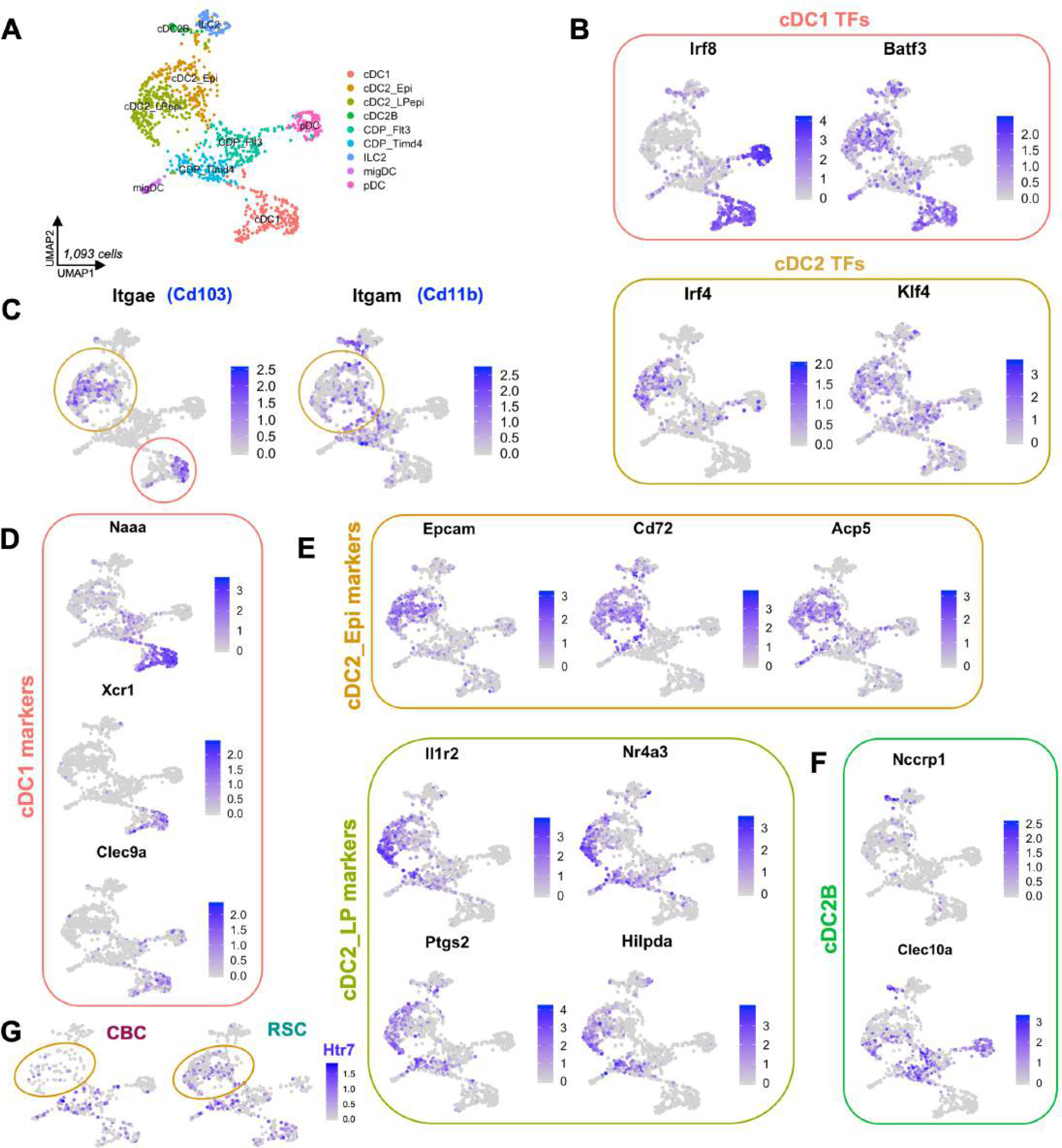
Related to Figure 6**: Conventional dendritic cells exhibit the most diversity amongst dendritic cell subtypes in the orthotopic tumor microenvironment.** A: UMAP clustering of dendritic cell (DC) subtypes: conventional DC 1 and 2 (cDC1, cDC2, cDC2B) with additional cDC2 heterogeneity defined in (E), migratory DC (migDC), plasmacytoid DC (pDC), common DC progenitor (CDP), and innate lymphoid group 2 cells (ILC2). B: Gene expression plots of master transcription factor regulators for cDC1 (/rf8, *Batf3)* and cDC2 *(lrf4, Klf4}* lineages. C: Gene expression plots of intestinal specific cDC marker genes for cDC1 (Cd103+ Cd11b-) and cDC2 (Cd103+ Cd11b+) populations. D: Gene expression plots of cDC1 marker genes with distinctive expression of migratory marker *(Xcr1).* E: Gene expression plots of genes expressed in cDC2 subtypes associated with Epithelial (cDC2_Epi) or Lamina Propria (cDC2_LP) compartmental localization (markers from Rivera et al. 2022, ref. 32). F: Gene expression plots of genes identifying cDC2B cells *(C/ec10a and Rorc).* G: Gene expression plot of Htr7 expression in cDC2 cells, enriched in RSC tumors.

**Figure S23:**
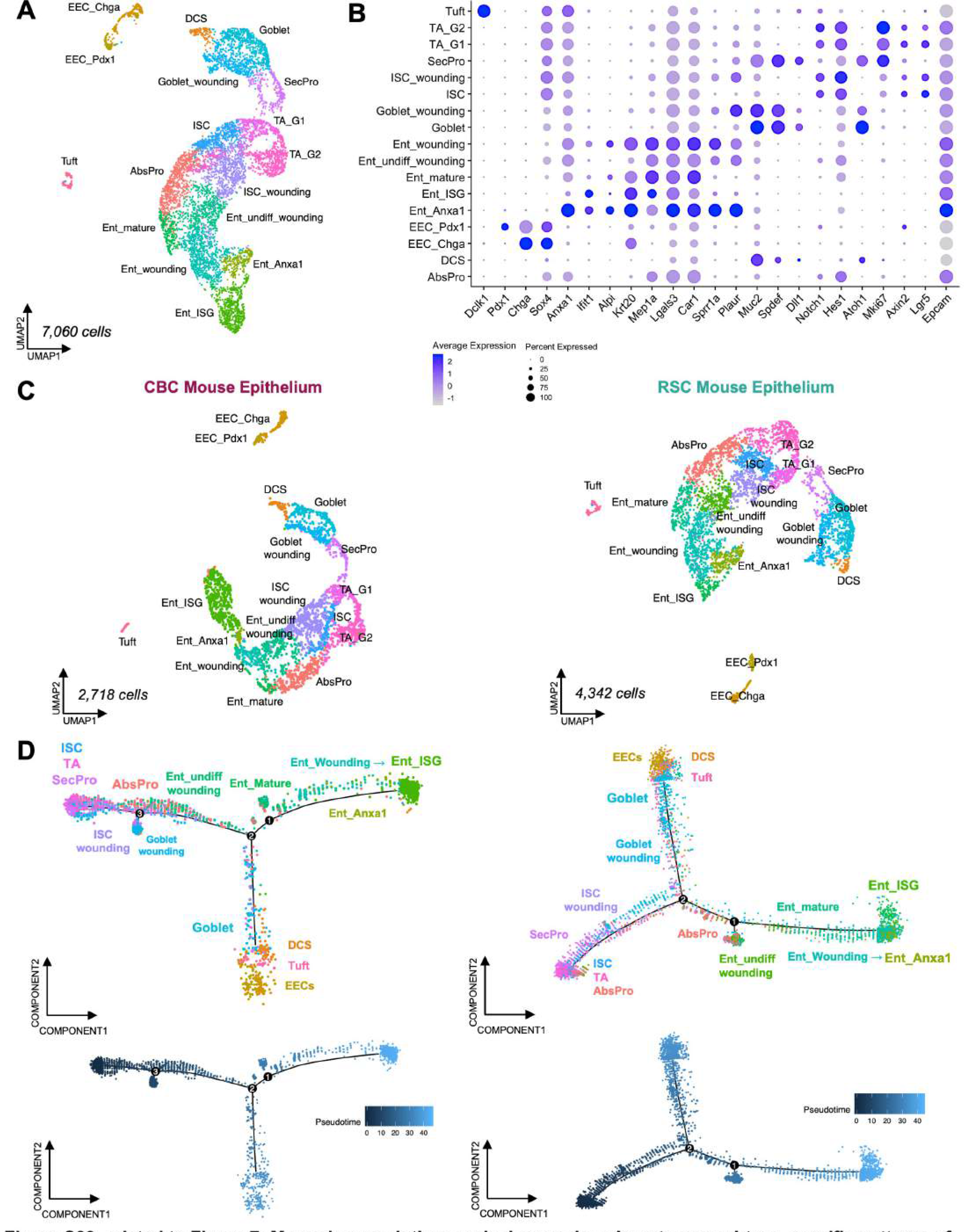
Related to Figure 7**: Monocle pseudotime analysis reveals unique tumor subtype-specific patterns of epithelial wounding and differentiation.** A: UMAP clustering of *Epcam+* mouse epithelial cells and wounded counterparts in orthotopic tumors: intestinal stem cells (ISCs), Absorptive Progenitors (AbsPro), Secretory Progenitors (SecPro), immature *Krt20-* enterocytes (Ent_undiff_wounding), mature *Krt20+* non-wounded (Ent_mature) and wounded (Ent_wounding) enterocytes, and wounded-specific populations of *Anxa1+* (Ent_Anxa1) and interferon stimulated gene expressing (Ent_lSG) enterocytes, normal and wounded goblet cells, tuft cells, and *Chga+* enteroendocrine (EEC_Chga) and *Pdx1+* enteroendocrine (EEC_Pdx1) cells. B: DotPlot showing biomarkers of intestinal epithelial cell populations and genes involved in lineage specification and differentiation. C: UMAP clustering of mouse epithelial cells separated by CBC (left) and RSC (right) orthotopic tumor identity. D: Monocle2 analysis of CBC (left) and RSC (right) tumor associated mouse epithelial populations orders cells by transcriptional similarities creating a lineage differentiation trajectory, measured by ’pseudotime’ (bottom graphs). The ISC compartment and early progenitors mark the beginning of ’pseudotime’ and wounded differentiated populations in CBC tumors generally lie closer in pseudotime to the ISC populations, whereas wounded differentiating populations (enterocytes and goblet cells) in RSC tumors are farther along the pseudotime trajectory indicating less damage and an earlier stage of tumor­ induced wounding.

**Figure S24:**
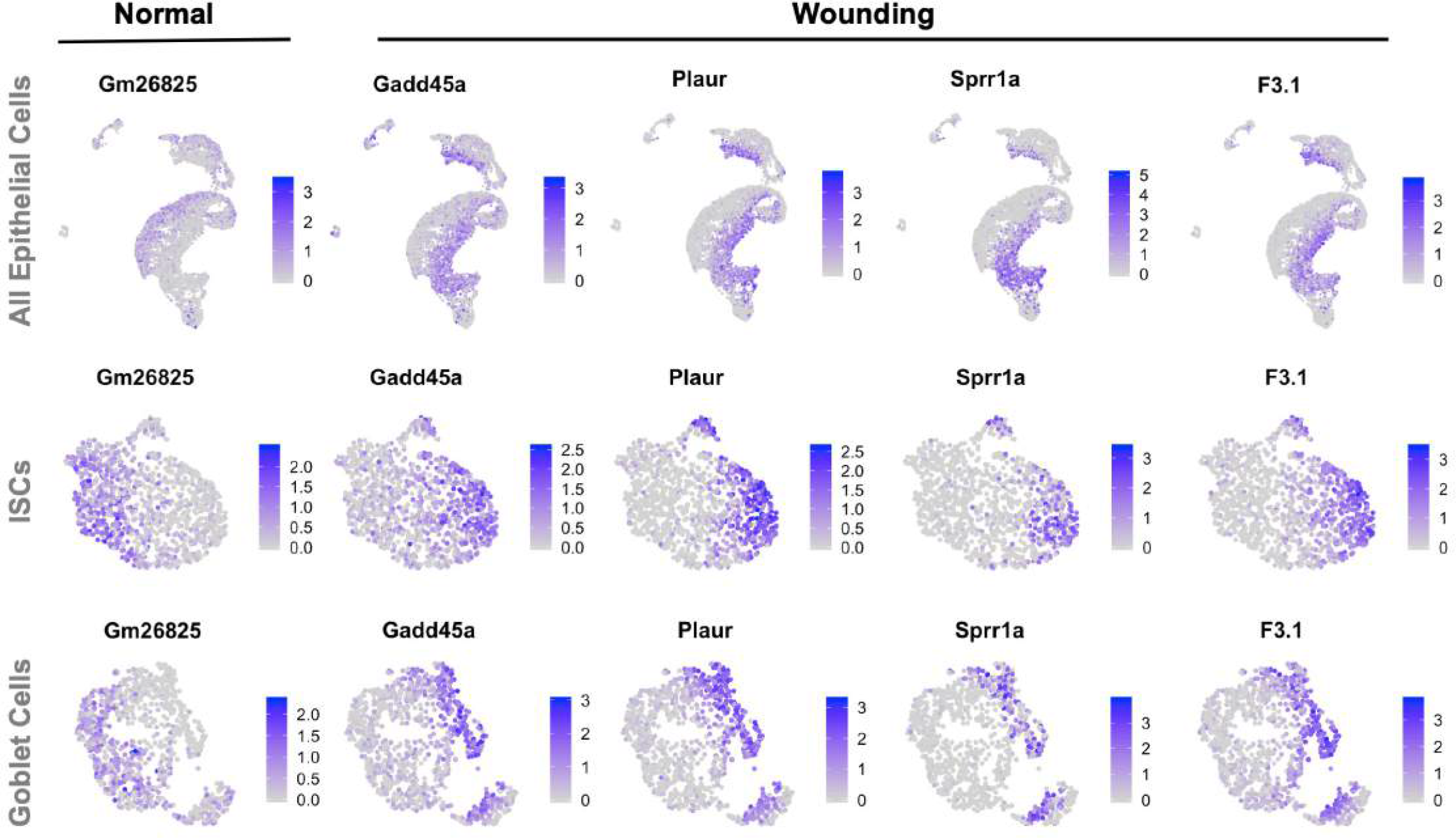
Related to Figure 7**: Biomarkers of normal and wounding-associated genes expressed in multiple mouse epithelial cell populations.** UMAPs of all mouse epithelial cells (top), the ISC compartment (middle), and the goblet population (bottom) have distinct expression of normal and wounding-associated genes. *Gm26825,* a lncRNA, is expressed in non-wounded (normal tissue) populations of epithelial cells from both tumor subtypes, while *Gadd45a, Plaur Sprr1a* and *F3.1* are examples of genes upregulated in tumor-induced wounding states of mouse epithelia. The large wounded enterocyte population is evident in the lower right of the UMAP of all epithelial cells.

**Figure S25:**
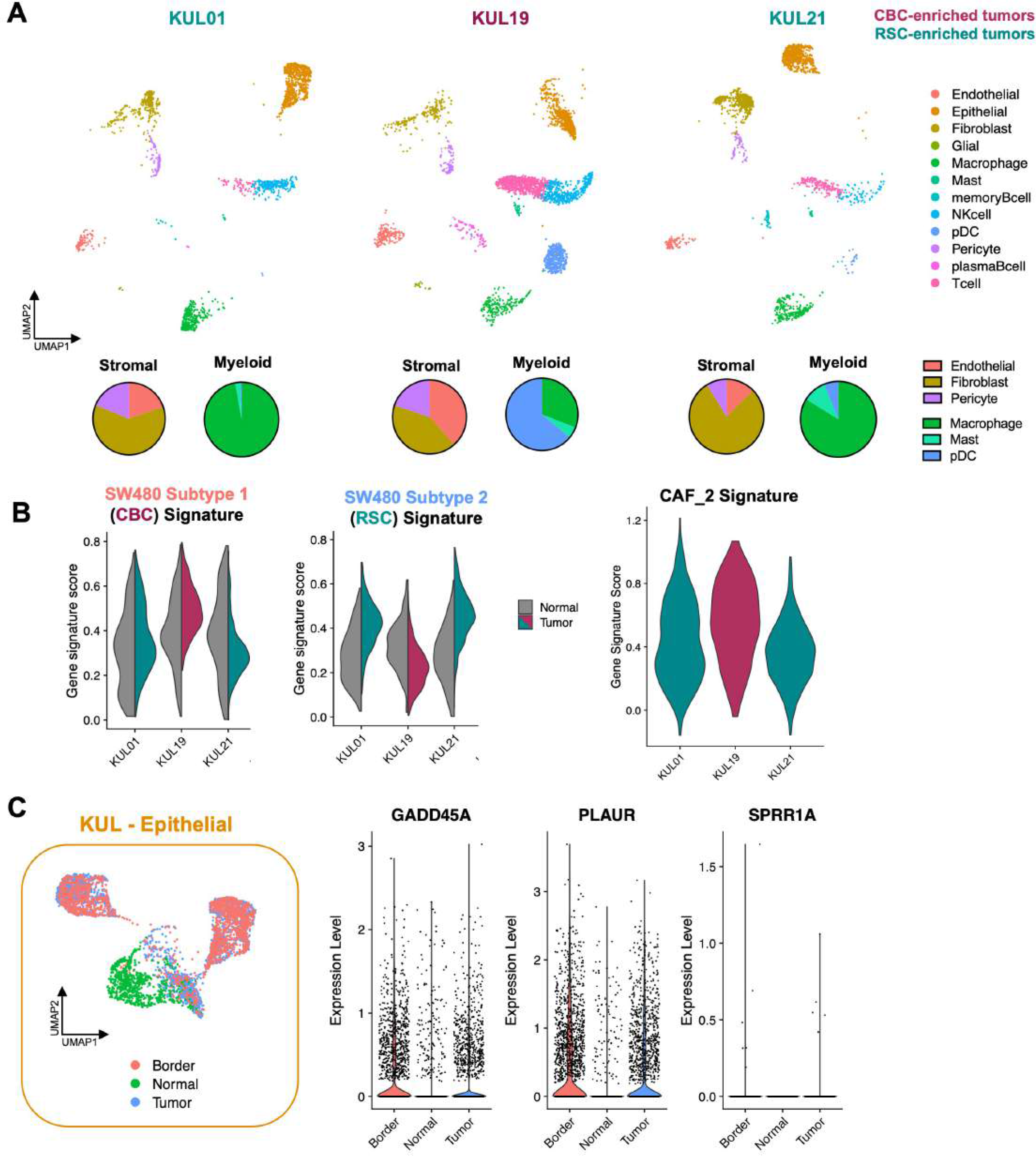
Related to Figure 2**: CRC patient TME correlates with SW480 subtype-specific TME compositions and phenotypes.** A: UMAP clustering of the tumor microenvironment from Belgian (KUL) patient tumor biopsies. Pie charts indicate differing compositions of stromal and myeloid cells in CBC- and RSC-enriched patient tumors. B: Violin plots of gene scoring for SW480-derived CBC and RSC signatures show that CBC-enriched patient tumors have CAFs with a higher CAF_2 signature, which correlates with a worse prognosis. C: Epithelial cells from normal, border, and tumor biopsies of Belgian (KUL) patients show that bystander border but not distant normal tissue expresses wound-specific biomarkers *GADD45A, PLAUR,* and *SPRR1A*.

**Figure S26:**
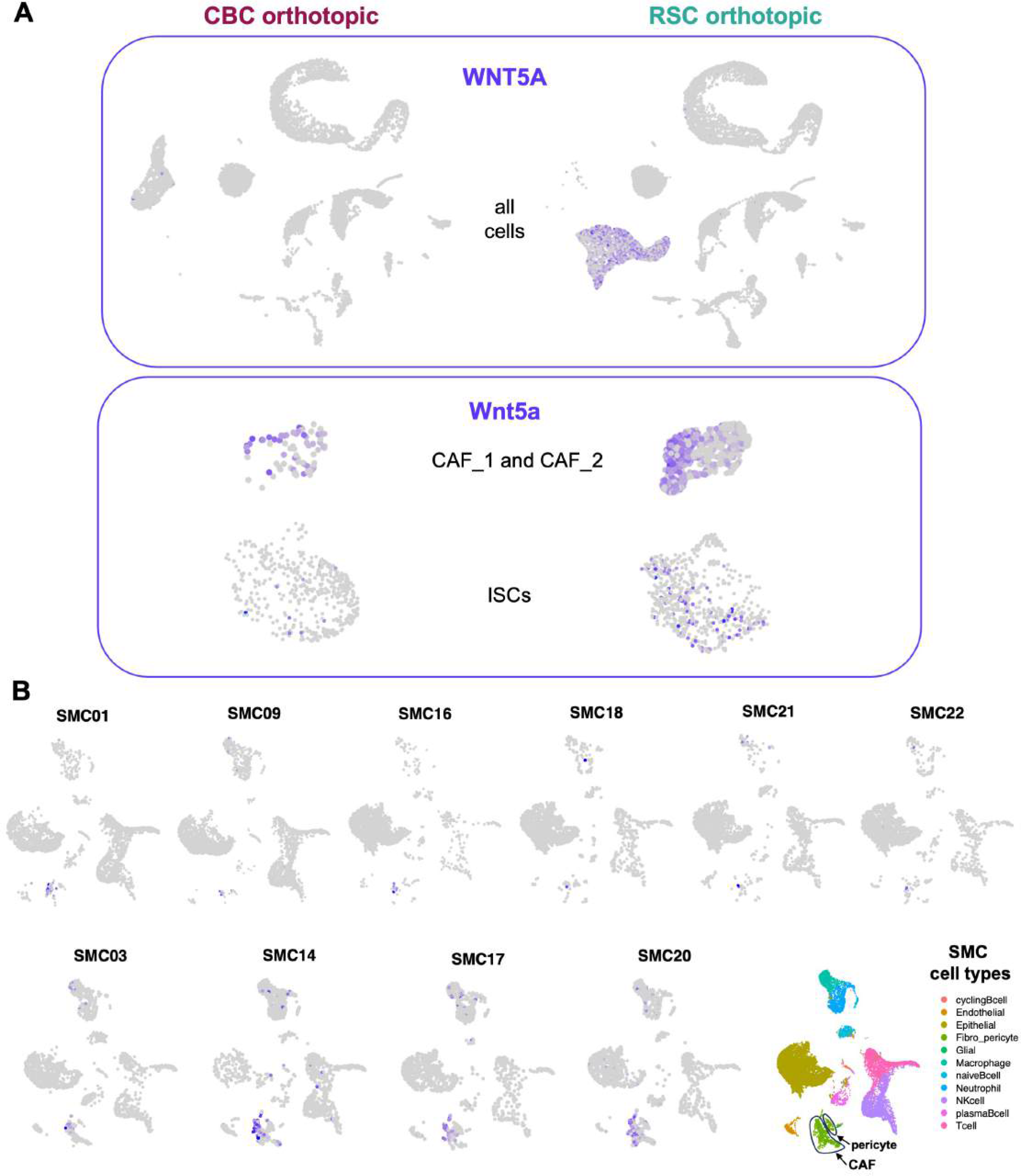
Inflammatory ligand WNT5A/Wnt5a is predominantly expressed in RSC tumors. Gene expression plots of non-canonical WNT expression in the tumor microenvironment show the major sources of this inflammatory signal come from tumor cells (specifically WNTSA from the RSC subtype), cancer associated fibroblasts (CAFs; specifically CAF_1 in the RSC TME), and Intestinal Stem Cells (ISCs; predominantly in the RSC TME) in the orthotopic tumor model (A), and WNTSA is expressed by CAFs in RSC enriched patient (SMC14, SMC17, SMC20) tumors (B).

## Notes

https://www.ncbi.nlm.nih.gov/geo/query/acc.cgi?acc=GSE254890

https://www.ncbi.nlm.nih.gov/geo/query/acc.cgi?acc=GSE254891

### Competing Interest Statement

The authors have declared no competing interest.

## References

1. Roerink, S.F., Sasaki, N., Lee-Six, H., Young, M.D., Alexandrov, L.B., Behjati, S., Mitchell, T.J., Grossmann, S., Lightfoot, H., Egan, D.A., et al. (2018). Intra-tumour diversification in colorectal cancer at the single-cell level. Nature 556, 437–462. 10.1038/s41586-018-0024-3.

2. Guinney, J., Dienstmann, R., Wang, X., de Reyniès, A., Schlicker, A., Soneson, C., Marisa, L., Roepman, P., Nyamundanda, G., Angelino, P., et al. (2015). The consensus molecular subtypes of colorectal cancer. Nat. Med. 2015 2111 21, 1350–1356. 10.1038/nm.3967.

3. Isella, C., Brundu, F., Bellomo, S.E., Galimi, F., Zanella, E., Porporato, R., Petti, C., Fiori, A., Orzan, F., Senetta, R., et al. (2017). Selective analysis of cancer-cell intrinsic transcriptional traits defines novel clinically relevant subtypes of colorectal cancer. Nat. Commun. 8, 15107. 10.1038/ncomms15107.

4. Cheung, P., Xiol, J., Dill, M.T., Yuan, W.C., Panero, R., Roper, J., Osorio, F.G., Maglic, D., Li, Q., Gurung, B., et al. (2020). Regenerative Reprogramming of the Intestinal Stem Cell State via Hippo Signaling Suppresses Metastatic Colorectal Cancer. Cell Stem Cell 27, 590–604.e9. 10.1016/J.STEM.2020.07.003.

5. Pearson, J.D., Huang, K., Pacal, M., McCurdy, S.R., Lu, S., Aubry, A., Yu, T., Wadosky, K.M., Zhang, L., Wang, T., et al. (2021). Binary pan-cancer classes with distinct vulnerabilities defined by pro- or anti-cancer YAP/TEAD activity. Cancer Cell 39, 1115–1134.e12. 10.1016/j.ccell.2021.06.016.

6. Joanito, I., Wirapati, P., Zhao, N., Nawaz, Z., Yeo, G., Lee, F., Eng, C.L.P., Macalinao, D.C., Kahraman, M., Srinivasan, H., et al. (2022). Single-cell and bulk transcriptome sequencing identifies two epithelial tumor cell states and refines the consensus molecular classification of colorectal cancer. Nat. Genet. 54, 963–975. 10.1038/s41588-022-01100-4.

7. Vazquez, E.G., Nasreddin, N., Valbuena, G.N., Mulholland, E.J., Belnoue-Davis, H.L., Eggington, H.R., Schenck, R.O., Wouters, V.M., Wirapati, P., Gilroy, K., et al. (2022). Dynamic and adaptive cancer stem cell population admixture in colorectal neoplasia. Cell Stem Cell 29, 1213–1228.e8. 10.1016/J.STEM.2022.07.008.

8. Mustata, R.C., Vasile, G., Fernandez-Vallone, V., Strollo, S., Lefort, A., Libert, F., Monteyne, D., Pérez-Morga, D., Vassart, G., and Garcia, M.I. (2013). Identification of Lgr5-Independent Spheroid-Generating Progenitors of the Mouse Fetal Intestinal Epithelium. Cell Rep. 5, 421–432. 10.1016/j.celrep.2013.09.005.

9. Solé, L., Lobo-Jarne, T., Álvarez-Villanueva, D., Alonso-Marañón, J., Guillén, Y., Guix, M., Sangrador, I., Rozalén, C., Vert, A., Barbachano, A., et al. (2022). P53 Wild-Type Colorectal Cancer Cells That Express a Fetal Gene Signature Are Associated With Metastasis and Poor Prognosis. Nat. Commun. 13. 10.1038/s41467-022-30382-9.

10. Merlos-Suárez, A., Barriga, F.M., Jung, P., Iglesias, M., Céspedes, M.V., Rossell, D., Sevillano, M., Hernando-Momblona, X., Da Silva-Diz, V., Muñoz, P., et al. (2011). The Intestinal Stem Cell Signature Identifies Colorectal Cancer Stem Cells and Predicts Disease Relapse. Cell Stem Cell 8, 511–524. 10.1016/J.STEM.2011.02.020.

11. Muñoz, J., Stange, D.E., Schepers, A.G., Van De Wetering, M., Koo, B.K., Itzkovitz, S., Volckmann, R., Kung, K.S., Koster, J., Radulescu, S., et al. (2012). The Lgr5 intestinal stem cell signature: Robust expression of proposed quiescent ′ +4′ cell markers. EMBO J. 31, 3079–3091. 10.1038/emboj.2012.166.

12. Alderdice, M., Richman, S.D., Gollins, S., Stewart, J.P., Hurt, C., Adams, R., Mccorry, A.M., Roddy, A.C., Vimalachandran, D., Isella, C., et al. (2018). Prospective patient stratification into robust cancer-cell intrinsic subtypes from colorectal cancer biopsies. J. Pathol. 245, 19–28. 10.1002/PATH.5051.

13. Petrova, T. V., Nykänen, A., Norrmén, C., Ivanov, K.I., Andersson, L.C., Haglund, C., Puolakkainen, P., Wempe, F., von Melchner, H., Gradwohl, G., et al. (2008). Transcription Factor PROX1 Induces Colon Cancer Progression by Promoting the Transition from Benign to Highly Dysplastic Phenotype. Cancer Cell 13, 407–419. 10.1016/j.ccr.2008.02.020.

14. Pate, K.T., Stringari, C., Sprowl-Tanio, S., Wang, K., TeSlaa, T., Hoverter, N.P., McQuade, M.M., Garner, C., Digman, M.A., Teitell, M.A., et al. (2014). Wnt signaling directs a metabolic program of glycolysis and angiogenesis in colon cancer. EMBO J. 33, 1454–1473. 10.15252/embj.201488598.

15. Lee, M., Chen, G.T., Puttock, E., Wang, K., Edwards, R.A., Waterman, M.L., and Lowengrub, J. (2017). Mathematical modeling links Wnt signaling to emergent patterns of metabolism in colon cancer. Mol. Syst. Biol. 13, 912. 10.15252/msb.20167386.

16. Barker, N., van Es, J.H., Kuipers, J., Kujala, P., van den Born, M., Cozijnsen, M., Haegebarth, A., Korving, J., Begthel, H., Peters, P.J., et al. (2007). Identification of stem cells in small intestine and colon by marker gene Lgr5. Nature 449, 1003–1007. 10.1038/nature06196.

17. Ishikawa, K., Sugimoto, S., Oda, M., Fujii, M., Takahashi, S., Ohta, Y., Takano, A., Ishimaru, K., Matano, M., Yoshida, K., et al. (2022). Identification of Quiescent LGR5+ Stem Cells in the Human Colon. Gastroenterology 163, 1391–1406.e24. 10.1053/j.gastro.2022.07.081.

18. Ayyaz, A., Kumar, S., Sangiorgi, B., Ghoshal, B., Gosio, J., Ouladan, S., Fink, M., Barutcu, S., Trcka, D., Shen, J., et al. (2019). Single-cell transcriptomes of the regenerating intestine reveal a revival stem cell. Nature, 1. 10.1038/s41586-019-1154-y.

19. Miyoshi, H., Ajima, R., Luo, C.T., Yamaguchi, T.P., and Stappenbeck, T.S. (2012). Wnt5a potentiates TGF-β signaling to promote colonic crypt regeneration after tissue injury. Science (80-.). *338*, 108–113. 10.1126/science.1223821.

20. Leibowitz, B.J., Zhao, G., Wei, L., Ruan, H., Epperly, M., Chen, L., Lu, X., Greenberger, J.S., Zhang, L., and Yu, J. (2021). Interferon b drives intestinal regeneration after radiation. Sci. Adv. 7, 1–13. 10.1126/sciadv.abi5253.

21. Lee, H.-O., Hong, Y., Etlioglu, H.E., Cho, Y.B., Pomella, V., Van den Bosch, B., Vanhecke, J., Verbandt, S., Hong, H., Min, J.-W., et al. (2020). Lineage-dependent gene expression programs influence the immune landscape of colorectal cancer. Nat. Genet. 2020 526 52, 594–603. 10.1038/s41588-020-0636-z.

22. Tomita, N., Jiang, W., Hibshoosh, H., Warburton, D., Kahn, S.M., Weinstein, I.B., and Development, [D W (1992). Isolation and Characterization of a Highly Malignant Variant of the SW480 Human Colon Cancer Cell Line.

23. Yoon (2008). The tumorigenic, invasive and metastatic potential of epithelial and round subpopulations of the SW480 human colon cancer cell line. Mol. Med. Rep. 10.3892/mmr_00000026.

24. Ke, J., Wu, X., He, X., Lian, L., Zou, Y., Wang, H., Luo, Y., Wang, L., and Lan, P. (2012). A subpopulation of CD24 + cells in colon cancer cell lines possess stem cell characteristics. Neoplasma 59. 10.4149/neo_2012_036.

25. Wang, Y., Xu, X., Maglic, D., Dill, M.T., Mojumdar, K., Ng, P.K.S., Jeong, K.J., Tsang, Y.H., Moreno, D., Bhavana, V.H., et al. (2018). Comprehensive Molecular Characterization of the Hippo Signaling Pathway in Cancer. Cell Rep. 25, 1304–1317.e5. 10.1016/j.celrep.2018.10.001.

26. Yui, S., Azzolin, L., Maimets, M., Pedersen, M.T., Fordham, R.P., Hansen, S.L., Larsen, H.L., Guiu, J., Alves, M.R.P., Rundsten, C.F., et al. (2018). YAP/TAZ-Dependent Reprogramming of Colonic Epithelium Links ECM Remodeling to Tissue Regeneration. Cell Stem Cell 22, 35–49.e7. 10.1016/j.stem.2017.11.001.

27. Tan, M.S.Y., Sandanaraj, E., Chong, Y.K., Lim, S.W., Koh, L.W.H., Ng, W.H., Tan, N.S., Tan, P., Ang, B.T., and Tang, C. (2019). A STAT3-based gene signature stratifies glioma patients for targeted therapy. Nat. Commun. 2019 101 10, 1–15. 10.1038/s41467-019- 11614-x.

28. Busslinger, G.A., Weusten, B.L.A., Bogte, A., Begthel, H., Brosens, L.A.A., and Clevers, H. (2021). Human gastrointestinal epithelia of the esophagus, stomach, and duodenum resolved at single-cell resolution. Cell Rep. 34, 108819. 10.1016/J.CELREP.2021.108819.

29. Brügger, M.D., Valenta, T., Fazilaty, H., Hausmann, G., and Basler, K. (2020). Distinct populations of crypt-associated fibroblasts act as signaling hubs to control colon homeostasis. PLOS Biol. 18, e3001032. 10.1371/JOURNAL.PBIO.3001032.

30. Puram, S. V., Tirosh, I., Parikh, A.S., Patel, A.P., Yizhak, K., Gillespie, S., Rodman, C., Luo, C.L., Mroz, E.A., Emerick, K.S., et al. (2017). Single-Cell Transcriptomic Analysis of Primary and Metastatic Tumor Ecosystems in Head and Neck Cancer. Cell 171, 1611–1624.e24. 10.1016/J.CELL.2017.10.044.

31. Varricchi, G., Galdiero, M.R., Loffredo, S., Marone, G., Iannone, R., Marone, G., and Granata, F. (2017). Are mast cells MASTers in cancer? Front. Immunol. 8, 424. 10.3389/FIMMU.2017.00424/BIBTEX.

32. Rivera, C.A., Randrian, V., Richer, W., Gerber-Ferder, Y., Delgado, M.G., Chikina, A.S., Frede, A., Sorini, C., Maurin, M., Kammoun-Chaari, H., et al. (2022). Epithelial colonization by gut dendritic cells promotes their functional diversification. Immunity 55, 129–144.e8. 10.1016/J.IMMUNI.2021.11.008.

33. Schuijs, M.J., and Halim, T.Y.F. (2018). Group 2 innate lymphocytes at the interface between innate and adaptive immunity. Ann. N. Y. Acad. Sci. 1417, 87–103. 10.1111/NYAS.13604.

34. Rodrigues, P.F., Alberti-Servera, L., Eremin, A., Grajales-Reyes, G.E., Ivanek, R., and Tussiwand, R. (2018). Distinct progenitor lineages contribute to the heterogeneity of plasmacytoid dendritic cells. Nat. Immunol. 2018 197 19, 711–722. 10.1038/s41590-018-0136-9.

35. Aspord, C., Leccia, M.T., Charles, J., and Plumas, J. (2013). Plasmacytoid dendritic cells support melanoma progression by promoting Th2 and regulatory immunity through OX40L and ICOSL. Cancer Immunol. Res. 1, 402–415. 10.1158/2326-6066.CIR-13-0114-T.

36. Zhou, B., Lawrence, T., and Liang, Y. (2021). The Role of Plasmacytoid Dendritic Cells in Cancers. Front. Immunol. 12, 1–10. 10.3389/fimmu.2021.749190.

37. Leoni, G., Neumann, P.A., Kamaly, N., Quiros, M., Nishio, H., Jones, H.R., Sumagin, R., Hilgarth, R.S., Alam, A., Fredman, G., et al. (2015). Annexin A1’containing extracellular vesicles and polymeric nanoparticles promote epithelial wound repair. J. Clin. Invest. 125, 1215–1227. 10.1172/JCI76693.

38. Ratziu, V., Lalazar, A., Wong, L., Dang, Q., Collins, C., Shaulian, E., Jensen, S., and Friedman, S.L. (1998). Zf9, a Kruppel-like transcription factor up-regulated in vivo during early hepatic fibrosis. Proc. Natl. Acad. Sci. U. S. A. 95, 9500–9505. 10.1073/pnas.95.16.9500.

39. Isella, C., Terrasi, A., Bellomo, S.E., Petti, C., Galatola, G., Muratore, A., Mellano, A., Senetta, R., Cassenti, A., Sonetto, C., et al. (2015). Stromal contribution to the colorectal cancer transcriptome. Nat. Genet. 47, 312–319. 10.1038/ng.3224.

40. Lotti, F., Jarrar, A.M., Pai, R.K., Hitomi, M., Lathia, J., Mace, A., Gantt, G.A., Sukhdeo, K., DeVecchio, J., Vasanji, A., et al. (2013). Chemotherapy activates cancer-associated fibroblasts to maintain colorectal cancer-initiating cells by IL-17A. J. Exp. Med. 210, 2851–2872. 10.1084/jem.20131195.

41. Deng, L., Jiang, N., Zeng, J., Wang, Y., and Cui, H. (2021). The Versatile Roles of Cancer-Associated Fibroblasts in Colorectal Cancer and Therapeutic Implications. Front. Cell Dev. Biol. 9, 1–16. 10.3389/fcell.2021.733270.

42. Brügger, M.D., and Basler, K. (2023). The diverse nature of intestinal fibroblasts in development, homeostasis, and disease. Trends Cell Biol. xx, 1–16. 10.1016/j.tcb.2023.03.007.

43. Murakami, M., Ikeda, T., Ogawa, K., and Funaba, M. (2003). Transcriptional activation of mouse mast cell protease-9 by microphthalmia-associated transcription factor. Biochem. Biophys. Res. Commun. 311, 4–10. 10.1016/j.bbrc.2003.09.148.

44. Sharma, M.D., Baban, B., Chandler, P., Hou, D.Y., Singh, N., Yagita, H., Azuma, M., Blazar, B.R., Mellor, A.L., and Munn, D.H. (2007). Plasmacytoid dendritic cells from mouse tumor-draining lymph nodes directly activate mature Tregs via indoleamine 2,3- dioxygenase. J. Clin. Invest. 117, 2570–2582. 10.1172/JCI31911.

45. Chen, W., Liang, X., Peterson, A.J., David, H., Blazar, B.R., and Munn, D.H. (2008). The Indoleamine 2,3-Dioxygenase Pathway Is Essential for Human Plasmacytoid Dendritic Cell-Induced Adaptive T Regulatory Cell Generation. J. Immunol. 181, 5396–5404.

46. Nguyen, Q.H., Pervolarakis, N., Blake, K., Ma, D., Davis, R.T., James, N., Phung, A.T., Willey, E., Kumar, R., Jabart, E., et al. (2018). Profiling human breast epithelial cells using single cell RNA sequencing identifies cell diversity. Nat. Commun. 9, 1–12. 10.1038/s41467-018-04334-1.

47. Stuart, T., and Satija, R. (2019). Integrative single-cell analysis. Nat. Rev. Genet. 20, 257–272. 10.1038/s41576-019-0093-7.

48. Subramanian, A., Tamayo, P., Mootha, V.K., Mukherjee, S., Ebert, B.L., Gillette, M.A., Paulovich, A., Pomeroy, S.L., Golub, T.R., Lander, E.S., et al. (2005). Gene set enrichment analysis: A knowledge-based approach for interpreting genome-wide expression profiles. Proc. Natl. Acad. Sci. U. S. A. 102, 15545–15550. 10.1073/pnas.0506580102.

49. Liberzon, A., Birger, C., Thorvaldsdóttir, H., Ghandi, M., Mesirov, J.P., and Tamayo, P. (2015). The Molecular Signatures Database Hallmark Gene Set Collection. Cell Syst. 1, 417–425. 10.1016/j.cels.2015.12.004.

